# An atlas of transcription factor cooperation reveals how motif readers shape regulatory output

**DOI:** 10.64898/2026.09.14.751590

**Authors:** Hanbei Xiong, Jieyuan Liu, Wei Wang

## Abstract

Regulatory motifs are conventionally associated with named transcription factors (TFs), yet a motif label need not identify the protein that reads the sequence or the regulatory consequence that follows in a given cell. We analyzed 1,552 TF binding datasets in 10 cell types using ARES, a multi-agent system that tests competing mechanisms of TF-motif dependencies in a specific cellular context against multi-omic data. We found that the inferred mechanisms converged on three operating routes: direct sequence recognition, protein-mediated recruitment or exclusion, and regulatory context. Importantly, the predictive motifs of the target TF binding were read by their conventionally “canonical” TFs in only one third of resolved dependencies, and these “canonical” TFs were expressed much less often than the inferred readers. Furthermore, we observed that motif similarity was associated with shared regulatory region type but not shared transcriptional outcome, whereas reader identity was associated with both and the only feature among the examined associated with outcome. In validation case studies where an inferred reader was perturbed, target TF occupancy fell in proportion to reader binding before perturbation, and a natural variant disrupting the predictive motif altered target TF binding at every intermediate step of the inferred mechanism. These observations were further supported by single-cell perturbation, in vitro cooperativity and evolutionary constraint. Together, these results separate motif identity from reader identity and regulatory output, suggesting that a motif acts as an address whose regulatory consequence is shaped in trans by the protein that interprets it.

## Introduction

Transcription factors (TFs) rarely regulate genes in isolation. Their genomic occupancy and regulatory activity emerge from interactions with other DNA-binding proteins, cofactors, chromatin regulators and the local regulatory environment.^1–7^ A target factor may recognize DNA directly, be recruited through a protein bound at a nearby motif, or respond to chromatin accessibility, histone state or three-dimensional genome organization. These mechanisms are well established in individual systems, but how they are distributed across transcription factors and cellular contexts, and what determines which one operates for a given TF-motif dependency in a specific cellular context, has not been systematically resolved.

Sequence-based analyses have greatly improved the prediction of TF occupancy and regulatory activity.^8–13^ Motif enrichment, machine-learning models and genomic foundation models can identify sequence features associated with binding and estimate the molecular consequences of sequence variation.^13–16^ However, a predictive association between a motif and a TF does not reveal the mechanism connecting them. The target factor may recognize the motif directly; a separate protein may bind the motif and recruit the target, or the motif may mark a regulatory environment that indirectly permits target occupancy. These alternatives can generate similar statistical relationships between sequence and binding while implying fundamentally different molecular mechanisms and different responses to perturbation.

Mechanistic interpretation is further complicated by the way regulatory motifs are represented. Motifs are conventionally assigned to “canonical” TFs that are assumed to universally bind to that motif. Yet related TFs often share sequence preferences, and proteins lacking sequence-specific DNA-binding activity can be recruited to sites through other factors.^17^ The protein that functionally reads a motif can therefore differ from its canonical reader and may depend on the proteins expressed in a particular cell. Determining which protein reads a predictive motif of a TF binding, and how that protein influences the target factor, requires evidence extending beyond DNA sequence. Hereinafter we refer to the “canonical” TFs of motifs as **namesake TFs** to distinguish the JASPAR-assigned TFs from the true readers of a motif.

The necessary evidence is distributed across heterogeneous data types. TF ChIP-seq^18^ measures protein occupancy, chromatin-accessibility^19^ and histone-modification assays describe regulatory state, Hi-C^20^ captures three-dimensional genome organization, expression and perturbation data reveal functional consequences, and protein-interaction networks nominate possible recruitment mechanisms. No individual assay is sufficient to distinguish direct sequence recognition, protein-mediated recruitment and context dependence.^6,7,19,20^ Resolving these alternatives requires generating competing hypotheses, gathering evidence for and against each hypothesis, identifying contradictions and selecting experiments capable of distinguishing the remaining possibilities. Large language model agents can plan analyses, invoke computational tools and reason over heterogeneous evidence, making them a candidate instrument for this task.^21–25^ Their application to biology, however, is non-trivial, such as when fluent model output can assert results that were never computed or mechanisms unsupported by any analysis, and recent surveys of biological agents identify such unfaithful reporting as a central obstacle to trustworthy automated discovery.^26–29^

Here we developed ARES (Automated Regulatory Explanation System), a framework for mechanistic inference of TF cooperation that integrates multimodal genomic and protein interaction evidence to propose, test and refine alternative explanations for why a partner motif predicts the occupancy of a target factor. Its reported results are based on real computation and concrete evidence, and its inferences are constrained by deterministic checks to avoid model hallucination, an approach to trustworthy inference that we adapt here to genomic mechanism discovery.^30–32^ We applied ARES to 1,552 TF-motif dependencies in specific cellular contexts, each derived from a distinct TF ChIP-seq experiment,^33^ across ten human cell types. We first identified motifs predictive of the target TF binding in each ChIP-seq, referred to as partner motifs thereinafter, based on which we constructed a mechanistic atlas of motif-associated TF cooperation. To place these dependencies on a common quantitative scale, we defined a partner-dependence coefficient measuring the extent to which a partner motif’s predictive effect depends on its namesake TF’s binding, which varied continuously rather than falling into discrete classes.

Despite the diversity of individual relationships, the mechanisms ARES inferred converged on three recurrent mechanistic routes. In the SEQUENCE route, the target factor directly interprets the predictive sequence. In the PROTEIN route, a distinct motif-bound protein recruits, stabilizes, co-binds with or excludes the target. In the CONTEXT route, motif-associated chromatin or genome organization influences target occupancy without a motif-bound protein acting directly on the target. These routes mapped onto the partner-dependence coefficient continuum, reproduced across cellular contexts, and were supported by orthogonal evidence: promoter reporter activity, evolutionary conservation, in silico motif perturbation, in vitro biochemical cooperativity, motif-centered ChIP-seq, single-cell perturbation and allele-specific binding from human genetic variation.

The atlas also allowed us to quantify, systematically and genome-wide, how often motif identity fails to specify protein identity. The namesake TFs were inferred to be readers of the motifs in only a third of resolved dependencies; where the two differed, the inferred reader, rather than the namesake, was expressed in the relevant cell type, occupied motif-centered loci and tracked target binding. Depleting an inferred reader reduced target occupancy in proportion to how much of that reader had been bound at each site. Because the same motif can be read by different proteins in different cells, the reader, not the sequence alone, shapes transcriptional output. These findings recast the partner motif as an address read in trans rather than a fixed instruction, and show how agentic integration of genomic evidence, when constrained to remain faithful to the underlying computation, can bridge the gap between accurate regulatory prediction and testable molecular explanation.

## Results

### TF cooperation spans a continuous spectrum of partner dependence

Transcription-factor (TF) binding, or more generally DNA-binding protein occupancy (for simplicity all the factors are referred to as generally defined TFs), is commonly interpreted through cognate DNA motifs, yet sites containing the same target motif often differ substantially in binding intensity. We asked whether additional sequence features could explain this variation. For each of 1,552 ChIP-seq experiments and each targeting a specific TF (i.e. **target TF**) across 10 human cell types, we trained a random-forest model to predict continuous target signal intensity from JASPAR motif scores and quantified the contribution of each motif using TreeSHAP (Fig. 1a, Supplementary Fig. 1).^35^ We used motif scores as features so that each feature is a named motif whose contribution can be read directly, at accuracy comparable to the state-of-the-art models^10,11,36–38^ (Supplementary Methods and Supplementary Fig. 8a). For more complex models such as those based on convolutional neural network (CNN), it requires a non-standard architecture to accommodate variable-length JASPAR motifs to evaluate each motif’s contribution to the prediction, yet this added complexity produced at most a marginal median accuracy gain over the random forest (Δr ≤ 0.02, Supplementary Fig. 8a).

**Figure 1.**
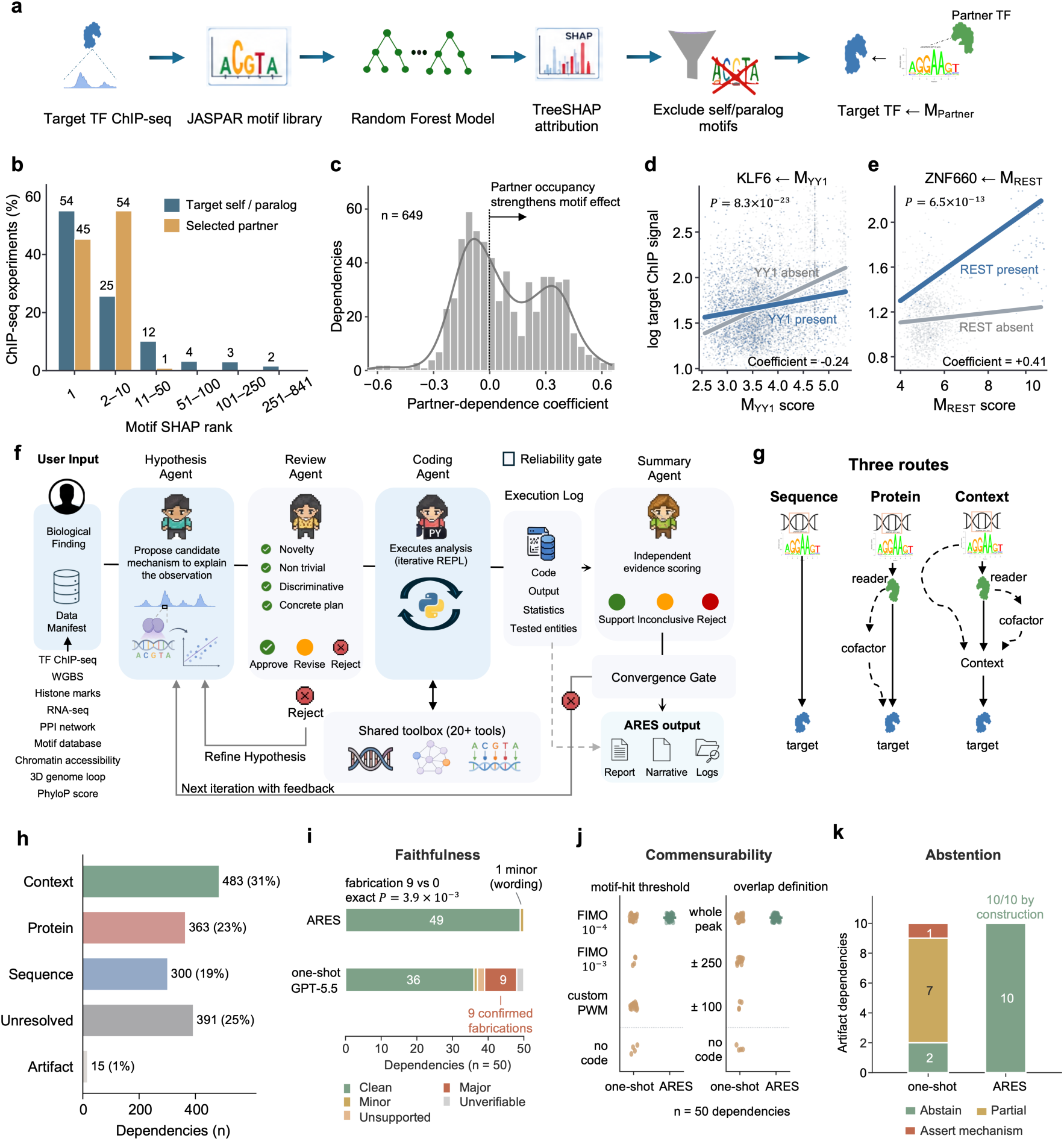
ARES converts motif-based binding predictions into an auditable mechanistic atlas. **a**, Partner-motif discovery workflow. A random-forest model predicts continuous target-TF occupancy from 841 JASPAR motif scores; motifs are ranked by TreeSHAP, and the top non-self, non-paralogous motif is selected as the partner motif. **b**, SHAP rank of the target’s own or paralogous motif versus the selected partner motif, across the 825 of 1,552 datasets whose target TF is also named in the JASPAR database (Methods). Ranks are out of the 841 motifs scored. **c**, Partner-dependence coefficients for the 649 of 1,552 dependencies for which the coefficient can be defined (Methods). The continuous, non-discrete distribution indicates graded motif-namesake dependence; positive values denote namesake-strengthened, negative values namesake-weakened motif effects. **d**, Negative-coefficient example: the YY1 motif predicts KLF6 occupancy more strongly at YY1-unbound loci (interaction, P = 8.3 × 10^-23^). **e**, Positive-coefficient example: the REST motif predicts ZNF660 occupancy primarily at REST-bound loci (interaction, P = 6.5 × 10^-13^). **f**, ARES framework. Multi-omic evidence is integrated through an agentic hypothesis–test–review loop to infer and audit the mechanism of each dependency, producing a mechanistic report, iteration narrative and traceable logs. **g**, Three mechanistic routes: SEQUENCE (motif read directly through DNA), PROTEIN (a reader protein binds the motif and acts on the target, often via a cofactor) and CONTEXT (the motif marks a regulatory context rather than a direct binding site). **h**, Atlas composition of 1,552 dependencies, CONTEXT 483 (31%), UNRESOLVED 391 (25%), PROTEIN 363 (23%), SEQUENCE 300 (19%) and 15 (1%) technical artifacts. **i**, Per-dependency faithfulness audit against saved code and logs (n = 50, 10 per category; scoring in Methods). The one-shot baseline (GPT-5.5) produced 9 reports with confirmed fabrications versus 0 for ARES (two-sided exact test on discordant dependencies, P = 3.9 × 10^-3^). **j**, Analytic parameters across dependencies for two helper-enforced operations (motif-hit and overlap definition). The baseline re-selected parameters per dependency, whereas ARES used fixed shared routines (FIMO 10^-4^; whole-peak overlap); each point is one dependency. Commensurability is assessed for helper-enforced operations only (Methods). **k**, Abstention on 10 artifact dependencies (target TPM < 0.5; a non-expressed factor cannot generate specific ChIP signal). ARES abstained on all 10 via a built-in artifact gate; the baseline assigned a context role or mechanism to 8 of 10, including TRIM22, an E3 ubiquitin ligase that does not bind DNA. Artifact criteria in Methods.

As expected, the target TFs’ own motif and close paralogs dominated the top-ranked predictors, consistent with canonical motif recognition.^8,9^ After excluding the motifs of the target factors and close paralogs, we identified the top motifs that contributed substantially to predict target TF binding as **partner motifs** (Fig. 1b). We define each retained cell-type-specific target TF–motif relationship as a **motif-mediated dependency**, represented by a unique (target TF, partner motif, cell type) tuple. The resulting atlas contained 1,552 motif-mediated dependencies derived from the corresponding ChIP-seq experiments, involving 864 target regulatory proteins and 309 partner motifs (Supplementary Fig. 1b). Unlike conventional motif discovery, which separates bound from unbound loci,^39,40^ our analysis here identifies motifs that explain why target TF binding strength varies among its bound regions. While the target TF binding is decided by many factors other than DNA sequence, the partner motifs captured regulatory information beyond canonical self-motif recognition and provided an opportunity to study TF cooperation mechanisms.

A partner motif could predict target occupancy due to reasons such as that its sequence is informative on its own or a protein bound to that motif influences the target. To quantify these contributions directly from genomic data, we modeled target TF binding as a function of partner-motif score, **motif-namesake-TF** (i.e. JASPAR named TF binding to that partner motif) binding, and their pairwise interaction, computed for the dependencies with available motif-namesake ChIP-seq data (Methods). We summarize each dependency by a partner-dependence coefficient (Methods): it compares how much target binding increases per unit of partner-motif score, at peaks where the motif-namesake is bound versus where it is absent. A positive coefficient means the motif predicts target occupancy more strongly where the motif namesake is bound; a coefficient near zero means the motif’s predictive effect is the same whether or not the protein is bound; and a negative coefficient means the motif predicts target occupancy less well where the motif namesake is bound, as expected when the partner competes with or occludes the target TF rather than assisting it.

The partner-dependence coefficient varied continuously across the atlas (Fig. 1c). Because a few partner motifs recur across many dependencies, we assessed multimodality after collapsing each motif to a single representative; the distribution was strongly unimodal (Hartigan’s dip test^41^, P = 0.99), supporting a continuous spectrum rather than discrete quantitative classes (Methods). Dependencies ranged from relationships in which partner-motif strength predicted target occupancy largely independently of motif-namesake binding to relationships in which the partner motif was informative almost exclusively at namesake co-bound loci. Thus, partner dependence is a graded property of TF occupancy rather than a small number of sharply separated binding states, revealing an analog rather than digital tuning of target TF binding.

The continuum was evident at individual dependencies. The YY1^42^ motif predicted KLF6 occupancy most strongly at loci where YY1 protein was absent, and less strongly where YY1 was bound (interaction P = 8.3 × 10^-23^; Fig. 1d). By contrast, the REST^43^ motif predicted ZNF660 occupancy almost exclusively at loci where REST protein was bound (interaction P = 6.5 × 10^-13^; Fig. 1e). Thus, the same statistical relationship, a partner motif predicting target binding, can arise either from sequence information that remains predictive without motif-namesake occupancy or from a namesake-dependent mechanism requiring a bound intermediary.

### ARES resolves three mechanistic routes underlying the quantitative continuum

The partner-dependence coefficient quantitatively describes motif-namesake dependence but does not identify the mechanism that produces it. A low or negative coefficient could arise because the target itself binds the partner motif, for instance where the motif is a degenerate form of a sequence the target TF recognizes or part of a composite element or sequence grammar the target reads, or because the partner motif marks a permissive chromatin environment rather than recruiting any protein. A strong positive partner-dependence coefficient could arise from cooperative binding, tethering or stabilization by the motif-namesake protein. Resolving which mechanism operates for a given dependency requires evidence beyond the coefficient itself.

We therefore developed ARES (Automated Regulatory Explanation System), a multi-agent framework that infers TF cooperation mechanisms from genomic evidence (Fig. 1f). It integrated TF and histone ChIP-seq, chromatin accessibility^44,45^, DNA methylation, ChromHMM states^46^, RNA-seq, evolutionary conservation^47^, Hi-C contacts and protein–protein interaction networks^48^. For each cell-type-specific dependency, ARES iteratively generated and tested competing mechanistic hypotheses linking the partner motif to the target TF binding, rejecting unsupported alternatives before producing a report summarizing the interpretation best supported by the available data. We implemented deterministic checks to constrain each judgement of large language model (LLM) and govern termination of inference, thus reducing fabrication and hallucination. ARES therefore uses LLMs to propose and interpret hypotheses, but requires mechanism assignments to be supported by traceable code output, standardized measurements, discriminative tests against simpler co-occupancy explanations and independent functional-genomic evidence for convergence (Methods; Supplementary Note 1).

ARES identified diverse mechanisms, including direct sequence recognition, embedded motifs, composite sequence grammar, cooperative co-binding, tethering, stabilization, competitive exclusion, accessibility dependence, chromatin-state dependence and three-dimensional genome organization. Despite this diversity, the paths connecting partner motif to target TF binding could be organized into three recurrent operating routes (Fig. 1g). In the SEQUENCE route, the predictive signal is carried by the DNA sequence itself: the partner motif may be read directly by the target TF, embedded within the target’s motif, part of a homotypic motif cluster, or embedded in a composite grammar or local DNA-shape feature that modulates target occupancy. In the PROTEIN route, a separate protein reads the partner motif and acts on the target by recruiting, tethering, co-binding with, stabilizing or excluding it. This protein need not be the TF for which the motif is named in JASPAR (i.e. the namesake TF) as different proteins can recognize similar motifs. In the CONTEXT route, the motif does not primarily act as a binding site for a protein that directly controls the target TF; instead, it marks or shapes a chromatin context, such as accessibility, promoter state, repressive chromatin, nucleosome positioning, three-dimensional architecture or local sequence composition, to which the target TF responds. Proteins may still bind at these loci, but the inferred path from partner motif to target TF is mediated through the shared context rather than through a direct motif-reader-to-target chain.

Across the atlas, CONTEXT was the most common route (483), followed by PROTEIN (363) and SEQUENCE (300), with 391 unresolved (i.e. the available evidence did not support any single route; Supplementary Table S1) and 15 artifacts (Fig. 1h). Each resolved dependency was therefore represented at two levels: a broad operating route describing the predominant path from partner motif to target TF binding, and a more specific molecular mechanism implementing that path (e.g. cooperative co-binding, tethering, competitive exclusion, accessibility proxy and chromatin state, Supplementary Fig. 1c).

### ARES generates faithful and commensurable evidence

Before drawing biological conclusions from these route assignments, we asked whether ARES’s reports can be trusted at the scale of the atlas. We benchmarked ARES against a one-shot agentic baseline built on a frontier model (GPT-5.5), across 50 dependencies sampled evenly from the five atlas categories (SEQUENCE, PROTEIN, CONTEXT, UNRESOLVED and ARTIFACT). Every report was audited against its saved code and execution logs (Methods).

The systems differed most in the faithfulness of the reported numbers to the analyses actually performed. The baseline produced nine reports containing confirmed **fabrications**, defined as statistics that contradicted its own saved output or were attributed to analyses it never ran. In one instance, it reported that motif-positive target TF peaks carried higher signal and dominated the top signal decile, while its saved output showed the opposite in both comparisons. ARES produced none (Fig. 1i; two-sided exact test on discordant dependencies, P = 3.9 × 10^-3^).

Because every ARES quantitative claim is written from logged computation rather than recalled by the model, a reported result cannot depart from executed analysis. The baseline’s fabrications fell into a small number of recurring failure modes: phantom analyses, fabricated constants and sign reversals, each structurally blocked by a corresponding safeguard in ARES (Supplementary Table S4).

For the core measurement operations that ARES routes through shared helpers with fixed default parameters — such as motif scanning and peak overlap — the baseline re-selected parameters for nearly every dependency. Motif thresholds ranged from custom percentiles to FIMO^49^ P < 10^-4^, and overlap definitions ranged from ±100 bp to whole peaks. Quantities such as motif deviance and co-occupancy were therefore not directly comparable across reports. ARES routed each operation through a fixed shared routine, generating evidence under common definitions before interpretation (Fig. 1j).

Most directly, we tested whether the systems reached the correct conclusion in cases where a specific regulatory mechanism was biologically untenable. Ten artifact dependencies involved factors that are not expressed (TPM < 0.5) and therefore cannot generate specific ChIP signal. The correct interpretation was thus not merely uncertain, but rather no regulatory route should be assigned at all. ARES identified and abstained on all ten through its built-in artifact gate, whereas the baseline assigned a regulatory route to eight. In seven the baseline reinterpreted non-specific signal as genuine sequence or chromatin context, and in one it asserted an explicit mechanism for an E3 ubiquitin ligase that does not bind DNA (Fig. 1k). Thus, in these cases where report-level mechanistic accuracy could be directly assessed, ARES made the expected abstention call for all ten dependencies, whereas the one-shot baseline converted absent biological signal into erroneous mechanisms.

### Mechanistic routes recur across cells and partner motifs

The three mechanistic routes showed biologically meaningful distribution along the partner-dependence-coefficient continuum. SEQUENCE-route dependencies were concentrated at low and negative values, PROTEIN-route dependencies at high values, and CONTEXT-route dependencies across an intermediate and broader range (Fig. 2a).

**Figure 2.**
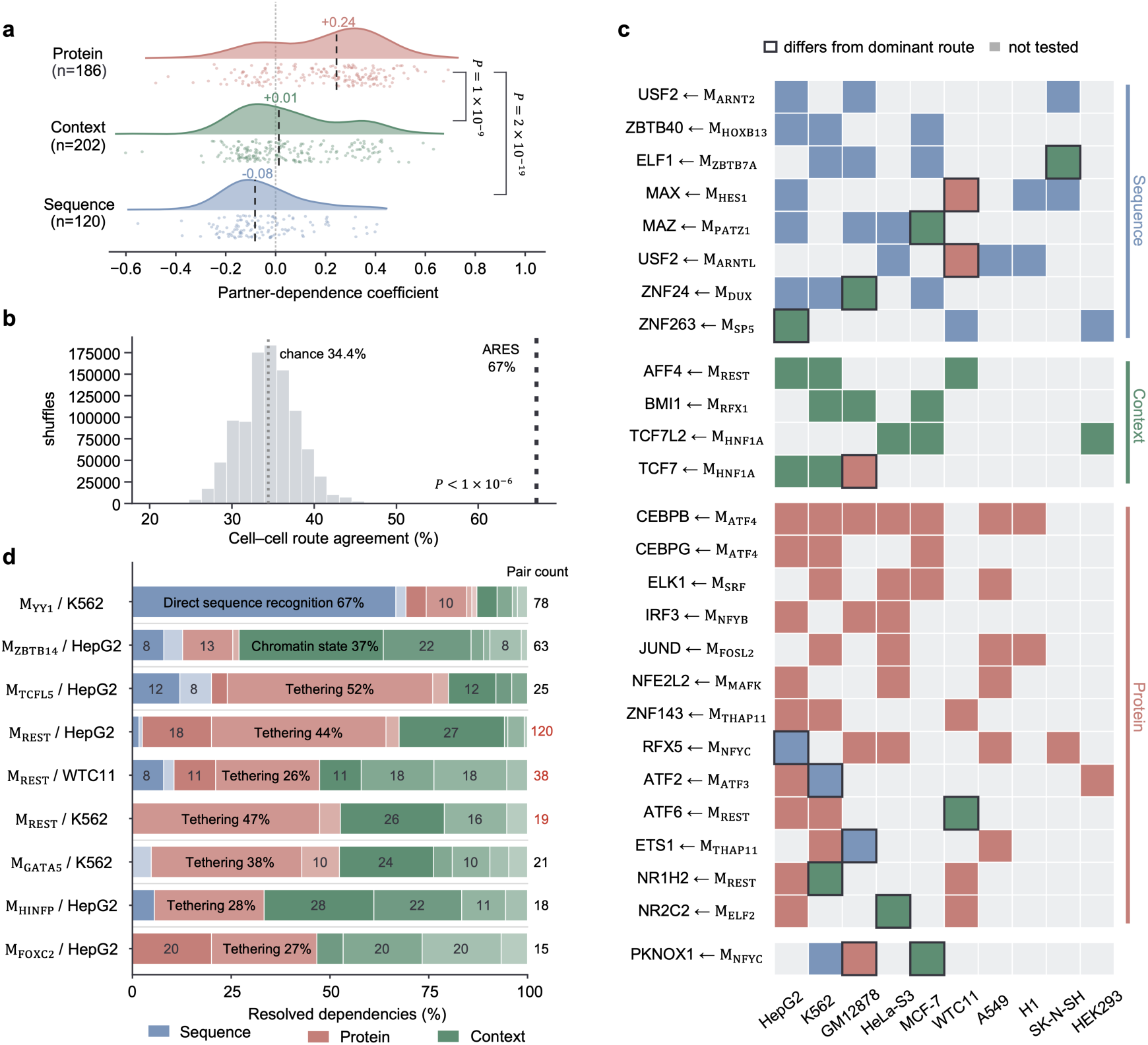
Mechanistic routes map onto the continuum and reproduce across cells. **a**, Distribution of partner-dependence coefficient across ARES routes for dependencies with available motif-namesake ChIP-seq data. Points represent dependencies; density curves show route-level distributions; vertical lines indicate medians. **b**, Cross-cell recurrent ARES routes. Route agreement between cell-type-specific dependencies sharing the same target TF and partner motif reached 67.0% concordance across 188 pairwise cross-cell comparisons from 90 recurrent TF–partner-motif groups. This exceeded a permutation null in which route labels were shuffled across dependencies while retaining each dependency’s recurrent-group membership and recomputing concordance over the same 188 within-group comparisons (null mean, 34.4%; z = 9.5; P < 1 × 10^-6^). **c,** Route assignments for recurrent TF-partner-motif groups observed in three or more cell types. Each row represents a recurrent TF–partner-motif group and columns represent cell types. Colored cells indicate assigned routes; grey indicates not tested. Outlined cells indicate assignments that differ from the dominant route for that recurrent group. **d,** Route composition of recurrent partner motifs across cell types. Bar widths indicate the fraction of resolved cell-type-specific dependencies assigned to each route; numbers at right indicate the number of dependencies represented for each motif–cell-line combination.

We next asked whether the inferred routes recur across cellular contexts. If a TF–partner-motif group was observed in k cell types, it contributed k observations and thus allowed in theory C(k, 2) pairwise cross-cell comparisons for concordance check. There were 90 recurrent TF–partner-motif groups with 219 cell-type-specific dependencies and 188 pairwise cross-cell comparisons, among which route concordance reached 67.0%. To estimate chance concordance, we permuted route labels across the 219 cell-specific dependencies while retaining their assignment to recurrent TF-partner-motif groups, and recomputed concordance over the same 188 within-group comparisons. The observed concordance substantially exceeded the resulting permutation mean of 34.4% (z = 9.5, P < 1 × 10^-6^; Fig. 2b and Methods). Recurrent TF-partner-motif groups observed in three or more cell types showed similar stability: 25 of 26 retained a dominant route in at least two-thirds of the cell types examined, whereas only a single group lacked a prevailing assignment (Fig. 2c). Thus, ARES route assignments were not arbitrary labels applied independently to each dependency, but captured recurrent biological properties of TF-partner-motif relationships across cellular contexts.

Partner motifs also recur across cellular contexts. A small number of motifs repeatedly emerged as the strongest predictors of TF occupancy across diverse targets and cell types, yet each tended to be preferentially associated with one operating route (Fig. 2d and Supplementary Fig. 2b). The REST motif provided a prominent example. Across multiple cell types and target factors, REST-associated dependencies were overwhelmingly assigned to PROTEIN route and rarely to SEQUENCE route. This enrichment is consistent with the established function of REST. REST binds RE1 elements through its zinc-finger DNA-binding domain^43,50^ and recruits corepressor assemblies, including CoREST^51^ and Sin3-HDAC^52^, so proteins localized to REST-bound loci need not recognize the RE1 sequence themselves; instead, the REST motif predicts their occupancy through recruitment into a REST-centered complex. REST-associated dependencies were therefore most frequently assigned to tethering mechanisms within the PROTEIN route (Supplementary Fig. 2c). RCOR1 illustrates this mechanism. RCOR1 is a core component of the CoREST complex that lacks sequence-specific DNA recognition,^51^ yet its occupancy was correlated with the REST motif, and only at loci where REST protein was bound. ARES inferred that the REST motif predicted RCOR1 occupancy through recruitment into the CoREST complex rather than through direct motif recognition (Supplementary Note 2).

### Partner motifs mark regulatory sequences and encode mechanism-associated perturbation signatures

The recurrent association of partner motifs with target occupancy suggested that they mark biologically relevant regulatory sequences rather than merely improving statistical prediction. We evaluated this possibility using three lines of orthogonal data: promoter activity^53^, evolutionary conservation^47^ and predicted molecular responses to sequence perturbation^15^.

Partner motifs showed regulatory activity in an independent promoter MPRA generated by PARM.^53^ PARM tiles each promoter into multiple overlapping fragments, some of which contain the partner motif (motif+) and some of which do not (motif−). We first balanced the motif+ and motif− fragment pools genome-wide for GC content, length and TSS distance by matched down-sampling, removing systematic sequence-composition differences between the two pools. To additionally control for locus-level activity, we then assessed the motif-associated activity effect within each parent promoter, computing the difference between the median activity of its motif+ and motif− fragments; because fragments from a common promoter are highly correlated, this within-promoter contrast isolates the motif effect from baseline differences between promoters (Methods). A target–partner dependency was considered supported if these per-promoter differences were consistently non-zero across promoters after Benjamini–Hochberg correction (Wilcoxon, q < 0.05). Across 989 testable dependencies from four PARM-covered cell lines, 358 (36%) showed a significant motif effect (BH-adjusted q < 0.05; Fig. 3a): 42% of HepG2 dependencies, 37% of HEK293 dependencies, 33% of MCF-7 dependencies and 25% of K562 dependencies. Thus, partner motifs are cis-active regulatory elements that alter transcriptional output where the target binds. This is a conservative lower bound: many partner motifs activate transcription at some promoters and repress it at others, and the signed test averages these opposing effects towards zero (Supplementary Fig. 3a,b).

**Figure 3.**
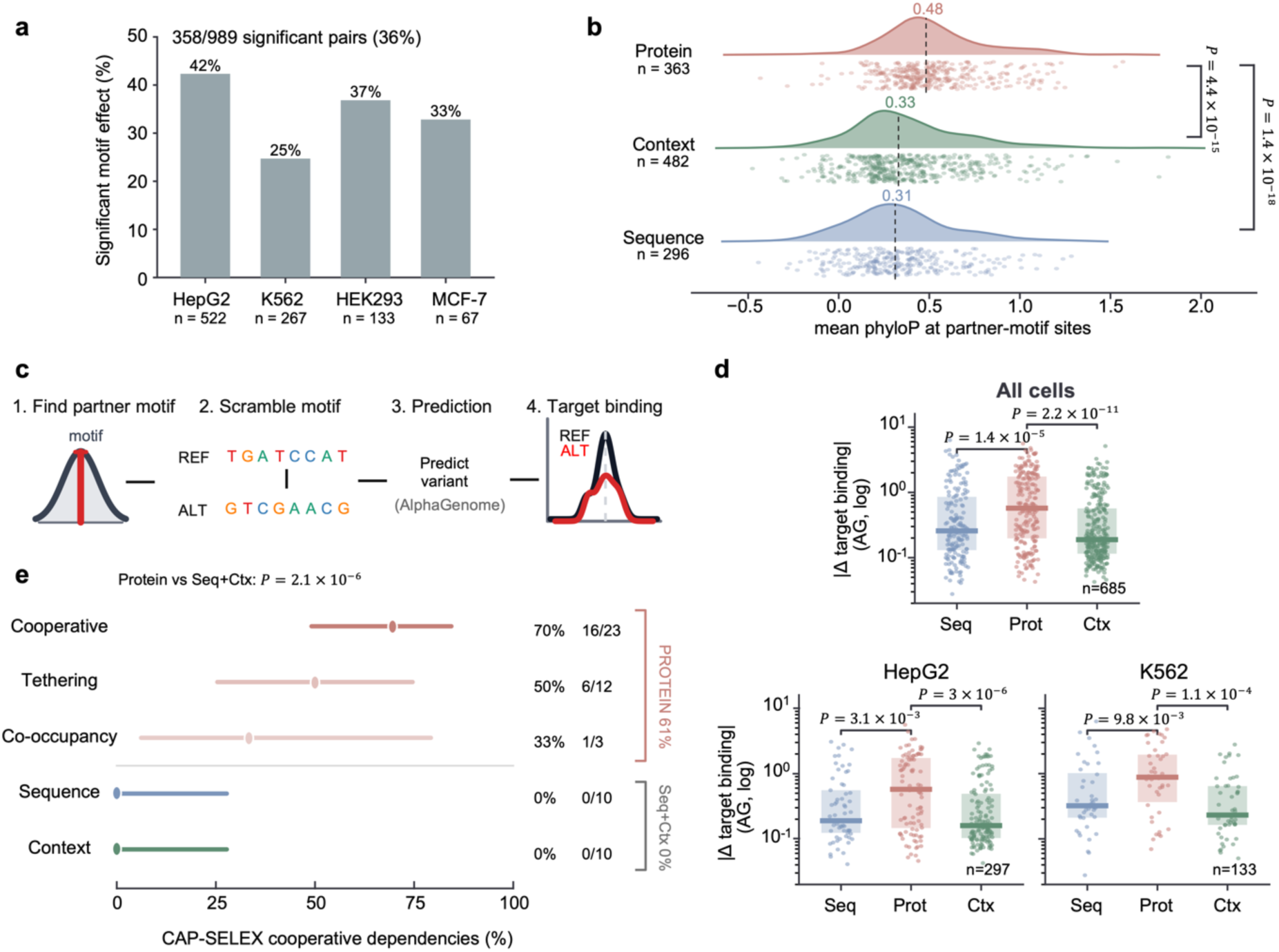
Partner motifs are functional, conserved and mechanistically perturbable. **a,** Fraction of testable dependencies with a significant partner-motif effect in promoter MPRA data after parent-locus aggregation and multiple-testing correction (q < 0.05). **b,** Mean phyloP conservation at partner-motif sites across ARES routes. Points show dependencies; density curves show route-level distributions; vertical lines indicate medians. **c,** AlphaGenome motif-perturbation workflow. Partner motifs were scrambled in silico and evaluated across predicted TF-binding, accessibility, chromatin and transcriptional tracks. **d,** Absolute predicted change in target binding (log_2_ fold-change) after partner-motif scrambling, measured relative to a matched substitution 60 bp away in the same peak (Methods). Each point is one dependency (median across up to 50 motif-positive target peaks). The pooled panel includes all eligible dependencies across cell lines. Individual-cell panels are shown for cell lines with more than 100 testable dependencies; full per-cell-line results are provided in Supplementary Fig. 4a. Boxes show interquartile ranges and medians. **e,** CAP-SELEX support for protein-mediated dependencies. Fractions indicate testable dependencies supported by cooperative in vitro DNA binding among assay-covered motif-associated factor dependencies, stratified by ARES route or PROTEIN-route mechanism. CAP-SELEX tests route-level biochemical cooperativity rather than the exact cellular reader for every dependency.

Partner motifs also showed evolutionary constraint that differed across operating routes. PROTEIN-route motifs showed the highest conservation, with a median phyloP score of 0.48, compared with 0.33 for CONTEXT and 0.31 for SEQUENCE-route motifs (SEQUENCE n = 296, PROTEIN n = 363, CONTEXT n = 482 dependencies with a scorable motif instance; Methods) (Fig. 3b; PROTEIN versus SEQUENCE P = 1.4 × 10^-18^). Because ARES uses conservation only to evaluate functional importance and not to determine the operating route itself, this gradient is independent of the classification procedure it tests. The ordering matches a direct mechanistic prediction: a SEQUENCE-route motif must satisfy the sequence preference of a single protein (the target or a close family member), whereas a PROTEIN-route motif must simultaneously support recognition by a separate reader and productive recruitment of the target — two independent evolutionary constraints rather than one, leaving less freedom for neutral sequence variation. The conservation gradient is therefore consistent with the number of proteins binding the motif must accommodate, matching the mechanistic hierarchy ARES infers.

We next tested whether partner motifs showed route-dependent sensitivity to sequence perturbation. For each dependency, the partner motif was scrambled and the resulting sequence was evaluated across molecular tracks using AlphaGenome for both the reference and scrambled sequences (Methods; Fig. 3c). Across cell lines, motif disruption reduced predicted target occupancy, with the strongest effects observed for PROTEIN-route dependencies (Fig. 3d). In HepG2, K562 and the pooled analysis, PROTEIN dependencies lost significantly more predicted target binding than SEQUENCE dependencies (pooled P = 1.4 × 10^-5^). This observation is consistent with conservation analysis and indicates that intermediary-protein-mediated recruitment is more sensitive to motif integrity than direct sequence recognition.

Together, MPRA activity, route-ordered evolutionary constraint and mechanism-specific perturbation signatures establish that partner motifs are functional cis-regulatory elements whose molecular behavior is specific to the mechanisms ARES infers.

### Biochemical cooperativity distinguishes protein-route mechanisms

We next asked whether the operating routes corresponded to distinct biochemical behavior. CAP-SELEX measures cooperative DNA binding by purified transcription-factor pairs in the absence of chromatin and most cellular cofactors.^17^ It directly tests cooperative co-binding and provides a selective, biochemical assay for protein-mediated mechanisms such as tethering and co-occupancy. Because CAP-SELEX assays purified factor pairs, whereas ARES infers cellular mechanisms that may involve motif-family members, hidden readers, cofactors or chromatin-dependent recruitment, we used CAP-SELEX to test whether PROTEIN-route dependencies are enriched for biochemical cooperativity among assay-covered motif-associated factor pairs, rather than to validate the exact ARES-inferred reader in every cellular context.

Fifty-eight atlas dependencies met the requirement that both the target and the motif-namesake TFs were represented in the CAP-SELEX compendium. CAP-SELEX detected cooperative binding in 23 of 38 protein-route dependencies (61%), compared with none of the 10 sequence-route and none of the 10 context-route dependencies (23 of 38 versus 0 of 20; Fisher’s exact P = 2.1 × 10^-6^; Fig. 3e). Thus, within the subset of the TF families covered by the assay, biochemical cooperativity was strongly enriched in PROTEIN-route dependencies and sharply distinguished them from the other operating routes.

Support differed across mechanism leaves. Cooperativity was highest among cooperative co-binding dependencies (70%, 16 of 23), and then in 50% of tethering (6 of 12) and 33% of co-occupancy dependencies (1 of 3). Support rates thus tracked the mechanism-leaf hierarchy, with the highest support where ARES explicitly inferred participation of two proteins in a shared DNA-bound complex.

The tethering cases clarified the distinction between biochemical cooperativity detected by CAP-SELEX and cellular motif interpretation revealed by ARES. CAP-SELEX-positive tethering assignments predominantly involved well-characterized heterodimer systems, including CEBPB–ATF4, CEBPG–ATF4, and FOSL1–JUN (Supplementary Table S6).^54,55^ These pairs are capable of cooperative DNA binding in vitro. In contrast, CAP-SELEX-negative tethering assignments included classical recruitment or ternary-complex systems such as ELK1–SRF, STAT2–IRF2 and GATA-associated recruitments,^56–58^ which may depend on cofactors, chromatin organization or binding geometries not reproduced under purified-protein conditions on naked DNA. CAP-SELEX therefore provides biochemical support for a subset of PROTEIN-route mechanisms, especially cooperative co-binding, but does not exhaust the cellular mechanisms through which motif-bound readers can influence target occupancy.

### The partner-motif reader is frequently distinct from the motif’s namesake factor

Each partner motif is named for the TF whose JASPAR profile it matches, but this name need not identify the protein that binds to the motif and cooperates with the target TF in a given cell. In fact, in many cases the motif-namesake factors were not expressed in the corresponding cell lines: in 121 (25.4%) of 477 namesake-cell-type combinations, the namesake-factor expressions were below 0.5 TPM in the cell type where the motif was identified (Fig. 4a, 19.3% below 0.1 TPM and 28.7% below 1 TPM). Furthermore, many of these unexpressed namesakes are also lineage-incompatible with the cell in which their motif was found: the meiosis-specific factor PRDM9 in HepG2, the neural factors OLIG2 and NHLH2 in the erythroleukemia line K562, and the autonomic-nervous-system factor PHOX2B in the lymphoblastoid line GM12878. For these dependencies the motif label cannot identify the protein occupying the site, because that protein is not there. Thus, motif names provide sequence annotations, but not necessarily cell-specific reader annotations.

**Figure 4.**
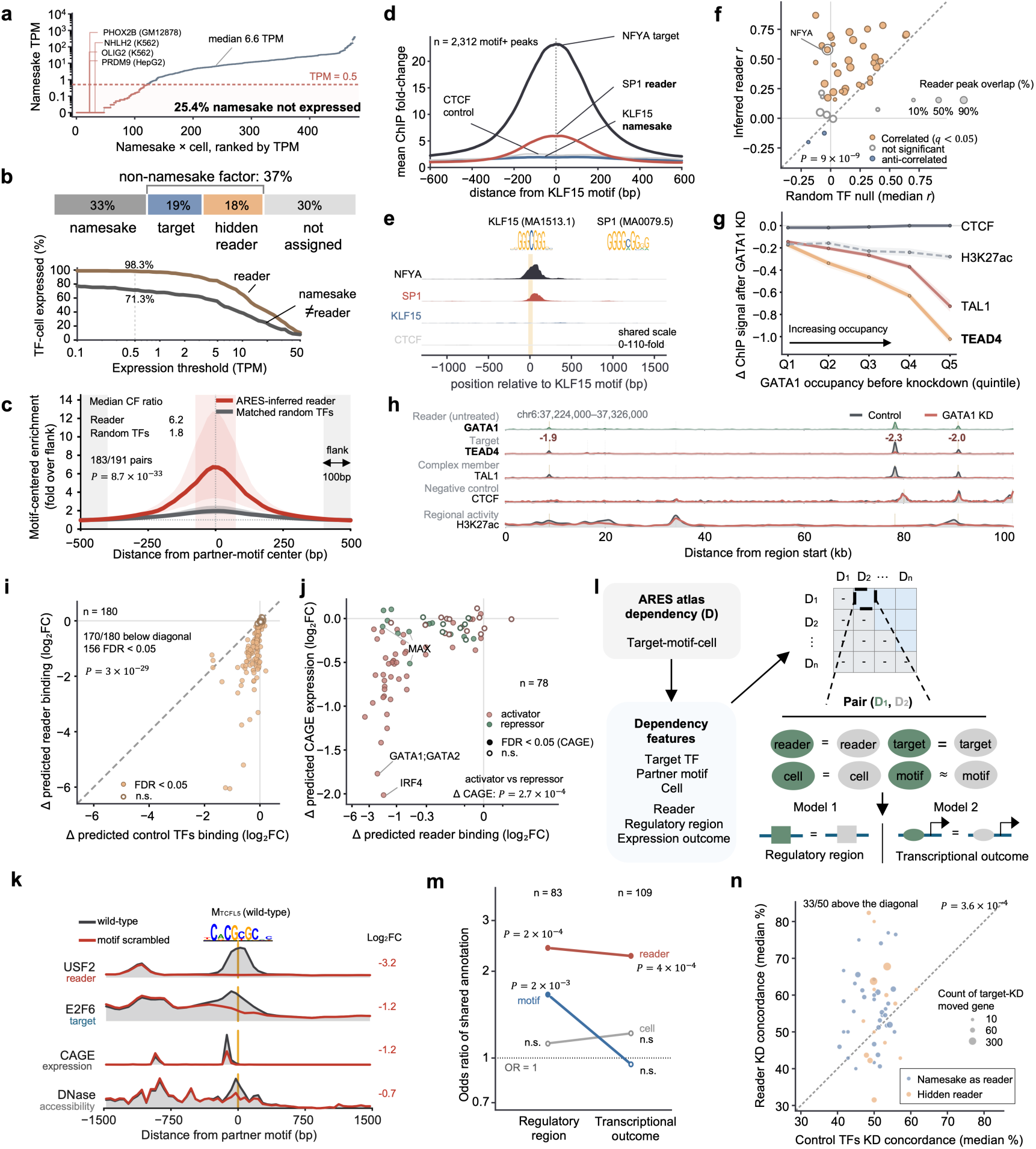
ARES identifies motif readers that explain target binding and regulatory output. **a,** Expression of the motif-namesake factors, in the cell type where that motif was identified. Namesakes are frequently not expressed in the cell. Three cell types (HEK293, HeLa-S3 and SK-N-SH) lacked matched total RNA-seq in ENCODE and are excluded; 1,258 of 1,552 dependencies remain, representing 477 unique namesake-cell-type combinations, so that a namesake factor recurring across many dependencies contributes once per cell type rather than once per dependency. Points are these 477 combinations, ranked by expression and plotted as TPM + 0.01 on a log scale; the lowest tier corresponds to TPM = 0 (45 combinations). Dashed line, TPM = 0.5, the expression threshold used throughout this study. 121 of 477 combinations (25.4%; shaded) fall below it, including lineage-incompatible namesakes such as PRDM9 in HepG2, OLIG2 and NHLH2 in K562, and PHOX2B in GM12878 — factors restricted to germ cells, the central nervous system and the autonomic nervous system, respectively. Median expression of motif-namesake factors across all combinations was 6.6 TPM. The fraction below cut-off varies with threshold: 19.3%, 25.4%, 28.7%, 32.9% and 43.4% at 0.1, 0.5, 1, 2 and 5 TPM. **b,** Reader assignments across the three-route-resolved atlas (n = 1,146 dependencies). For each dependency, ARES inferred the factor that recognizes the partner motif within target peaks, when one could be assigned. Reader assignments fell into four categories: the motif’s namesake partner (375 dependencies, 33%), the target factor itself recognizing the partner motif (216, 19%; Supplementary Note 3), a distinct third factor that was neither the target nor the namesake partner (“hidden reader”; 208, 18%), or no assigned reader (347, 30%). Lower panel, fraction of factors expressed as a function of expression threshold, counted per TF × cell-type combination rather than per dependency (same as in **a**). Brown, the assigned reader pooled across the three reader-assigned categories (n = 361); gray, the namesake in dependencies where a different factor (target or hidden reader) reads the motif (n = 178). At 0.5 TPM (dotted line) 98.3% of assigned readers are detected versus 71.3% of displaced namesakes; the gap widens with threshold (97.0 vs 68.0 at 1 TPM, 84.8 vs 55.6 at 5, 64.8 vs 38.2 at 10). Median expression 14.1 TPM for readers, 5.8 for displaced namesakes. **c,** Inferred readers are enriched at the partner motif above a matched null. Among the 208 hidden-reader dependencies, 191 were testable with both reader and target TF ChIP-seq data available and at least 30 partner-motif-positive target peaks present. For each dependency, reader ChIP-seq signal was quantified in a ±500-bp window centered on the partner motif within target peaks and expressed as fold enrichment over the mean signal in the flanking regions (computed as the average signals in the 400–500 bp from the motif center on each side to provide a stable background estimate). Curves show the median profile and interquartile range across dependencies. To estimate the background motif-centered enrichment expected at active regulatory elements, four TFs assayed in the same cell type were randomly selected for each dependency, balancing a more stable matched null estimate against the limited number of eligible TF profiles available across cell types; the target, inferred reader and partner-motif namesake were excluded. These factors were quantified at the identical loci, and their median profile was used as the matched null for that dependency. Both inferred readers and matched random TFs showed higher signals at the motif center, reflecting the fact that the analyzed loci are active regulatory elements rather than random genomic positions. However, inferred readers exhibited substantially stronger motif-centered enrichment than the matched random TFs — the center-to-flank ratio (mean signal within ±75 bp of the motif center divided by mean signal in the 400-500-bp flanks) was 6.2 versus 1.8 (median); the inferred reader exceeded its matched null in 183 of 191 dependencies (one-sided paired Wilcoxon signed-rank test, P = 8.7 × 10^-33^). Using a predefined centering criterion—a center-to-flank ratio of at least 2 and signal apex within 50 bp of the motif center—177 of 191 inferred readers (93%) were classified as motif-centered. **d,** Aggregate motif-centered ChIP enrichment for the KLF15-motif reassignment example (NFYA target in HepG2; n = 2,312 motif-positive NFYA peaks). Mean ChIP fold-change over flanking signal, centered on the KLF15 motif, is shown for the inferred reader SP1, the namesake KLF15, the NFYA target and a CTCF control. Additional reassignment examples in Supplementary Fig. 5c. **e,** A representative NFYA locus containing the KLF15 motif. Tracks share a common scale (0–110 fold). **f,** Hidden reader tracks the target at partner-motif loci where the namesake does not occupy. Each point is one hidden-reader dependency (n = 42). y, Spearman correlation between reader and target signal across those loci; x, the median of the same correlation for ten randomly chosen TFs with ChIP-seq in that cell line, over the identical loci. 37 of 42 lie above the diagonal (one-sided Wilcoxon signed rank on the paired values, P = 9 × 10^-9^; median r 0.43 versus 0.10). Fill indicates whether the reader–target correlation is itself significant (Benjamini–Hochberg q < 0.05): 34 significant positive, 6 not significant, 2 significant negative. Marker area scales with the fraction of tested loci inside a reader ChIP-seq peak (key: 10%, 50%, 90%; median 59%, range 7–99%). **g**, Perturbation experiments supported that GATA1 knockdown preferentially reduced TEAD4 occupancy at sites with stronger prior GATA1 binding. y, log_2_ fold-change of ChIP-seq signal after GATA1 knockdown; x, GATA1 occupancy before knockdown (n = 8,094 GATA5-motif-positive TEAD4 peaks), in quintiles (Q1 lowest, Q5 highest). TEAD4 is the target; CTCF (GATA1-independent) and H3K27ac (local enhancer activity) are controls, and TAL1 is a GATA1 complex member. TEAD4 is measured against a MAFF-knockdown control from the same experiment; the remaining factors against wild-type in the same experiment. **h**, Genome-browser example of a GATA5-labeled motif within a TEAD4 peak occupied by GATA1 before perturbation. Tracks show TEAD4, GATA1, TAL1, CTCF and H3K27ac at the same locus before and after GATA1 knockdown, illustrating reader-dependent loss of TEAD4 occupancy without comparable collapse of local chromatin activity. **i,** AlphaGenome-predicted change in reader binding after scrambling the partner motif, computed as in Fig. 3d. Each point is one dependency (n = 180). x, median predicted binding change for twelve control TFs drawn from the same cell line, excluding the target, partner, reader and their family members at the same loci; y, predicted reader binding change (both median log_2_ fold-change). **j,** AlphaGenome-predicted CAGE change after scrambling the partner motif, colored by reader function (activator versus repressor, assigned from GO; Methods). Each point is one dependency (n = 78). x, predicted reader binding change; y, predicted CAGE change (both median log_2_ fold-change). **k**, Example of AlphaGenome-predicted CAGE change in the direction following activator reader after scrambling the repressor-named motif. A TCFL5-named partner motif is inferred to be read by activator USF2 at an E2F6 target locus. Predicted tracks for the wild-type (unscrambled genomic sequence; grey) and motif-scrambled (red) sequences are shown for the reader USF2, the target E2F6, CAGE expression and DNase accessibility; values at right are log_2_ fold-change (scrambled vs wild-type). Scrambling reduces predicted reader binding (−3.2), target binding (−1.2), expression (−1.2) and accessibility (−0.7), as expected for an activator-read site. **l**, Schematic of the pairwise annotation-sharing analysis to identify features important for regulatory regions or transcriptional outcome. Each atlas dependency (D) is defined by a target TF, partner motif and cell type (observed), to which ARES adds an inferred reader, regulatory-region type and transcriptional-outcome type. ARES derived the transcriptional outcome type from RNA-seq (expression at motif-positive versus motif-negative target genes) independently of reader assignment (Methods), so any reader outcome association is not built into the annotations. The analysis is restricted to dependencies in which the ARES reader differs from the partner-motif namesake, so that reader identity carries information beyond the motif label. Every pair of dependencies (D_1_, D_2_) is scored for whether they share each of four features (represented as a binary variable for each feature comparison) — partner-motif similarity (Tomtom), reader, target and cell type — and, separately, whether they share an ARES annotation: the regulatory-region type (model 1) or the transcriptional outcome (model 2). Testing whether feature-sharing (e.g. common readers) predicts annotation-sharing (e.g. same transcriptional outcome) asks which property, the motif or the protein reading it, is more informative about where regulation occurs and about what it does. **m**, Features important for regulatory regions or transcriptional outcome. Odds ratios from logistic models were fit separately for each annotation (regulatory-region type (left) and transcriptional-outcome type (right)), with all four shared-feature terms included so each was adjusted for the others (n = 83 dependencies, 3,403 pairs for regulatory region; n = 109 dependencies, 5,886 pairs for transcriptional outcome). Points are odds ratios on a log scale; filled points are significant (P < 0.05) and open points are not significant, assessed by a node-permutation null in which the focal feature is permuted with the others held fixed. Lines connect the two annotations for each feature. A shared target TF was entered as a covariate but was not identifiable (quasi-complete separation from too few target-sharing dyads) and is omitted from the plot. Dotted line indicates OR = 1 (no association). **n,** Reader TF knockdown reproduces the target TF’s transcriptional response more often than knockdown of other TFs at the same sites. Each point is one dependency (n = 50, 19 readers). **y**, percentage of the target TF’s responsive genes moved in the same direction by reader TF knockdown; **x**, median of the same quantity across ten control TFs matched on occupancy of the motif-positive target peaks (Methods). An unrelated TF is expected at 50%. Point area, number of genes tested. 33 of 50 lie above the diagonal (paired median difference +5.6 percentage points; one-sided Wilcoxon P = 3.6 × 10^-4^).

Consistent with this distinction, ARES frequently assigned readers that differed from the motif namesake. Across 1,146 three-route-resolved dependencies, ARES assigned the namesake as the reader in only 33%; the target TF itself was inferred to recognize the motif in 19%, consistent with SEQUENCE-route interpretation; a **hidden reader** (neither the target TF nor the namesake TF) was assigned to 18%; and the remaining 30% lacked an assigned reader (Fig. 4b). In the direct recognition cases, ARES inferred the partner motif to be an alternate binding site for the target TF. This can occur because JASPAR motif labels are assigned to specific factors or fine-grained families, whereas DNA-binding specificity is often shared across broader DNA-binding-domain classes, including bHLH E-boxes, SP/KLF GC-boxes and nuclear-receptor half-sites (Supplementary Note 3). Most reader-unassigned dependencies belonged to the CONTEXT route, in which the partner motif is not interpreted through a single assigned reader but instead acts by marking chromatin state, accessibility or higher-order architecture (Supplementary Fig. 5a). In these cases, the absence of a reader is an expected property of the mechanism rather than a failure of annotation.

The inferred readers were also more compatible with the cellular context than the displaced namesakes (non-reader). Across expression thresholds, the factor ARES named as the reader was expressed far more often than the displaced namesake: at 0.5 TPM, 98.3% of assigned readers were expressed, compared with 71.3% of namesakes in dependencies where a different factor (target TF or hidden reader) was inferred to recognize the motif, and this gap widened at higher expression thresholds (Fig. 4b, lower panel). Median expression of readers was 14.1 TPM versus 5.8 TPM for displaced namesakes. Thus, the inferred reader was almost always present in the relevant cell type, whereas the factor the motif is named for frequently was not.

Next, we asked whether ARES-inferred readers were localized to the partner motifs. We focused on dependencies with a hidden reader because the nominated reader is neither the namesake nor the target factor and is therefore least obvious from the motif label alone. Across 191 testable hidden-reader assignments, reader ChIP-seq signal showed substantially stronger motif-centered enrichment than matched random TFs assayed in the same cell type and quantified at the same loci (median center-to-flank ratio 6.2 versus 1.8; reader greater than matched null in 183 of 191 assignments; one-sided paired Wilcoxon signed-rank test, P = 8.7 × 10^-33^; Fig. 4c). Using the predefined centering criterion, 177 of 191 reader profiles (93%) were centered on the partner motif. This supports that the inferred hidden reader occupies loci centered on the partner motif.

We next examined 42 hidden-reader dependencies where the namesake is expressed (TPM > 0.5) and its ChIP-seq is available. The inferred reader gave stronger motif-centered signal than the namesake in aggregate (mean peak enrichment 20.7-fold versus 8.4-fold, against 4.3-fold for a CTCF control; Supplementary Fig. 5b). NFYA illustrates this directly. The partner motif associated with NFYA (target) occupancy was best matched in JASPAR to the KLF15 motif.

ARES inferred that SP1 recognizes the motif and influences NFYA. Because KLF15 and SP1 both recognize the GC box motif, the motif label alone cannot identify which TF reads it. In aggregate ChIP-seq at NFYA peaks containing the KLF15 motif: only SP1 shows enrichment beyond the target itself where KLF15 and CTCF control do not bind at all (Fig. 4d, single-locus view in Fig. 4e).

Within each of the 42 dependencies, we restricted the analysis to target peaks containing the partner motif but lacking an overlapping namesake peak. Across these loci, reader signal correlated with target signal more strongly than did randomly chosen TFs quantified at the same positions in 37 of 42 dependencies (88%, median Spearman r 0.43 versus 0.10; one-sided Wilcoxon signed-rank test, P = 9 × 10^-9^; Fig. 4f). When removing the background signals, reader-target correlation remains robust where the random TFs collapse (Supplementary Fig. 5d). Therefore, hidden-reader signal continued to track target occupancy even at motif-positive loci where the expressed namesake factor was not detectably bound, supporting reader reassignment beyond simple namesake absence. NFYA is one of these dependencies (circled in Fig. 4f): at motif-positive NFYA peaks lacking a KLF15 peak, SP1 tracked NFYA occupancy, so the motif’s relationship to NFYA does not require the namesake to be bound there.

We then tested if target occupancy depends on the reader. We examined a K562 dependency in which the target was TEAD4 and the partner motif matched GATA5, whose expression was below the 0.5 TPM threshold in K562 (0.22 TPM). ARES inferred GATA1/GATA2 as the reader, consistent with the shared recognition preference of the GATA family and with GATA1 being the erythroid master regulator of K562.^59,60^ The ARES report proposed knockdown/knockout of the reader as the decisive test of whether TEAD4 binding collapses. Using publicly available ENCODE and GEO ChIP-seq datasets (Methods),^33,61,62^ we analyzed TEAD4 ChIP-seq after GATA1 knockdown, restricting the analysis to TEAD4 peaks containing the GATA5 motif and binning these sites into quintiles by GATA1 peak strength before knockdown. After GATA1 knockdown, TEAD4 occupancy declined in proportion to prior GATA1 occupancy (Fig. 4g, genome browser view in Fig. 4h). Since GATA1 is a master regulator in K562, we examined additional factors at the same sites to exclude a global collapse of these regions. CTCF, a GATA1-independent factor, showed no generic loss of signal; H3K27ac declined only slightly, far less than TEAD4, indicating the surrounding enhancer activity was largely preserved; and TAL1, a GATA1 complex member, showed an intermediate decline consistent with its well-characterized interaction with GATA1.^57^ TEAD4 declined most steeply of all, supporting its dependence on GATA1 while excluding global inactivation of the surrounding chromatin. A reader nominated by ARES, and named by neither the motif nor the target, is therefore required for full target occupancy, which is the test ARES itself proposed as decisive.

### Partner motifs provide the address and readers shape the outcome

Identifying readers is biologically meaningful if the same motif can produce different regulatory outcomes depending on the protein that interprets it. We tested this in silico with AlphaGenome by scrambling partner motifs within target TF peaks and measuring predicted changes in both reader binding and CAGE expression at the same loci. We focused on dependencies with a hidden reader, since the motif’s effect on target TF binding was already supported (Fig. 3d) and a namesake reader is trivial to test. Among 208 hidden-reader dependencies, 180 had an AlphaGenome–predicted reader track and sufficient target peaks to test. Motif disruption was predicted to significantly reduce reader binding in 156 of 180 dependencies (FDR < 0.05; Fig. 4i). To test whether this reflected reader-specific motif dependence rather than a generic response of the surrounding regulatory context, we scored twelve random TFs per dependency under the same scrambling. In 170 of 180 dependencies, reader binding dropped more than the random-TF median (P = 3 × 10^-29^), indicating reader responds to the motif itself rather than to the context.

To ask whether reader identity tracks the regulatory output, we examined the subset of the 180 dependencies whose CAGE track is covered by AlphaGenome. We used Gene Ontology (GO) to assign reader function (activator/repressor) and removed readers with ambiguous or mixed annotations (Methods). This left 78 testable dependencies, 58 with an activator reader and 20 with a repressor. Among them, 57 showed a significant predicted CAGE change (FDR < 0.05). Importantly, activator-reader dependencies showed significantly stronger CAGE decreases than repressor-reader ones (P = 2.7 × 10^-4^; Fig. 4j). Note that context-insensitive GO labels might not reflect the real function of a factor at an individual site: MAX, for example, is annotated as a repressor but acts as an activator as part of MYC-MAX heterodimer.^63^ Thus, the GO-based analysis supports a relative reader-valence effect but also shows why static activator/repressor labels are insufficient. This limitation is informative: it suggests that a factor’s regulatory direction is not a fixed property of the protein alone, but depends on the site and partners through which it acts, motivating the context-specific, data-derived outcome analysis below.

To illustrate why identifying the reader matters, we examined a dependency whose motif namesake is annotated as repressor TCFL5 in GO, but for which ARES inferred the activator USF2^64^ as the reader acting on the target E2F6. Scrambling the motif was predicted to lower the E2F6 binding, abolish USF2 binding, and reduce CAGE expression at the locus (Fig. 4k). Therefore, the direction of the transcriptional change is consistent with the activator USF2 instead of the repressor namesake.

We next tested, independently of AlphaGenome and of static GO annotations, whether reader identity was more informative than partner-motif identity about regulatory outcome across the atlas. We focused on dependencies in which the ARES-inferred reader differed from the partner-motif namesake, so that reader identity provided information beyond the motif label. The logic is the following: if readers shape the regulatory output (activation or repression), two dependencies sharing readers are expected to have similar transcriptional outcome, even if their partner motifs differ; sharing a similar partner motif might place two dependencies in the same class of regulatory regions (e.g. promoters and enhancers)^65^ but not necessarily produce the same transcriptional outcome.

ARES used RNA-seq to define transcriptional outcome (activation or repression) by comparing expressions at motif-positive versus motif-negative target genes. Since the ARES annotation is purely based on the data, it captures the direction of regulation actually observed at motif-positive target genes, independent from GO annotation of the reader as activator/repressor, which avoids circular argument (Supplementary Note 4). We considered all dependencies with both regulatory region and transcriptional outcome annotations by ARES. We then used two separate logistic regression models to respectively predict whether any pair of these dependencies share either regulatory region or transcriptional outcome using four binary features: similar partner motifs (Tomtom^66^ q < 0.05), same inferred reader, same target factor, and same cell type (Methods; Fig. 4l). Sharing the same reader was the strongest predictor of a shared transcriptional outcome and regulatory region type among the three identifiable features (regulatory region odds ratio (OR) 2.386, permutation P = 2 × 10^-4^; outcome OR 2.301, permutation P = 4 × 10^-4^; n = 83 dependencies, 3,403 pairs and n = 109 dependencies, 5,886 pairs, respectively; Fig. 4m). Similar partner motifs increased the odds that two dependencies occupied the same type of regulatory region (OR 1.69, permutation P = 2 × 10^-3^) but did not increase the odds that they had the same transcriptional outcome (OR 0.975, permutation P = 0.82). A shared cell type was not significantly associated with either regulatory region (OR 1.116, permutation P = 0.34) or transcriptional outcome (OR 1.166, permutation P = 0.057), and a shared target factor was not identifiable (too few target-sharing dependency pairs; Methods). Together, these analyses support that partner-motif similarity carried information about where regulation occurred but not what followed, whereas reader identity carried information about both and the only feature among the examined associated with transcriptional outcome.

Furthermore, if the reader acts on transcription through the target TF, knocking down the reader should move the target TF’s regulated genes in the same direction as knocking down the target TF itself. We tested this in genome-wide CRISPRi Perturb-seq in K562.^67^ For each dependency in which the reader is either the motif namesake or a hidden third factor, we took the genes whose promoters fall under motif-positive target peaks and that respond to knockdown of the target (median 25 genes per dependency to reduce noise; Methods), and compared their response to knockdown of the reader with their response to knockdown of ten control TFs that occupy the same peaks at a comparable rate (median 47.6% peak overlap for readers, 42.0% for controls). Matching occupancy ensured that a control TF does not differ from the reader merely by being absent from these sites. The matched controls of unrelated factors agreed with the target at 50.0% (IQR 47.1–53.1), whereas the reader agreed at 55.9% (IQR 50.0–61.6), exceeding its own controls in 33 of 50 dependencies spanning 19 readers (paired median difference +5.6 percentage points, and +4.3 in the 34 dependencies with a single nominated reader; one-sided Wilcoxon P = 3.6 × 10^-4^; Fig. 4n). This analysis suggested that the target’s own transcriptional response depends more on the ARES-inferred readers than on other factors occupying the same motif-marked sites.

### Human genetic variation supports route-specific motif function

Human genetic variation offers natural perturbations of regulatory sequences that are independent of the functional genomic evidence used to build the atlas, providing an orthogonal test of how partner motifs act.^68^ A binding quantitative trait locus (bQTL) is associated with allele-specific TF binding;^69,70^ when it falls within a partner motif, it directly alters the sequence feature predictive of target TF occupancy. If the operating routes reflect genuine differences in how a motif is interpreted, e.g. a direct reader binding site versus a marker of regulatory context, then naturally occurring variants should perturb target binding in a route-dependent manner.

We first characterized partner-motif bQTLs by operating routes. Starting from ∼270,000 allele-specific binding variants catalogued by ADASTRA,^68^ 41,393 overlapped target TF binding sites represented in the atlas. Among the 288 atlas dependencies whose target TF had allele-specific-binding data in ADASTRA (across the eight cell lines with ADASTRA coverage), 73 contained at least one bQTL within the partner motif (Fig. 5a). On-motif bQTL frequency decreased across routes as the motif’s role shifted from a direct reader binding site toward a marker of regulatory context: 40.0% of SEQUENCE-route, 24.5% of PROTEIN-route and 15.7% of CONTEXT-route dependencies carried an on-motif bQTL (Fig. 5b). This ordered trend was significant by a Cochran–Armitage test (P = 2.8 × 10^-4^), which does not adjust for confounders. To control for the two most likely confounders, we fitted a logistic regression for whether each dependency carried an on-motif bQTL, entering route as an ordinal score (SEQUENCE = 0, PROTEIN = 1, CONTEXT = 2) and including partner-motif width and the log-transformed number of target ASB variants (a proxy for testing power) as covariates (Methods); the route trend remained significant (P = 1.2 × 10^-3^). It was driven primarily by the two extremes of the axis, SEQUENCE versus CONTEXT, with PROTEIN-route dependencies intermediate, consistent with their motifs read indirectly through a separate reader rather than by the target TF itself. Thus, variants within partner motifs were most often associated with altered target binding when the motif was read directly as a binding site and progressively less often as its role shifted toward marking regulatory context.

**Figure 5.**
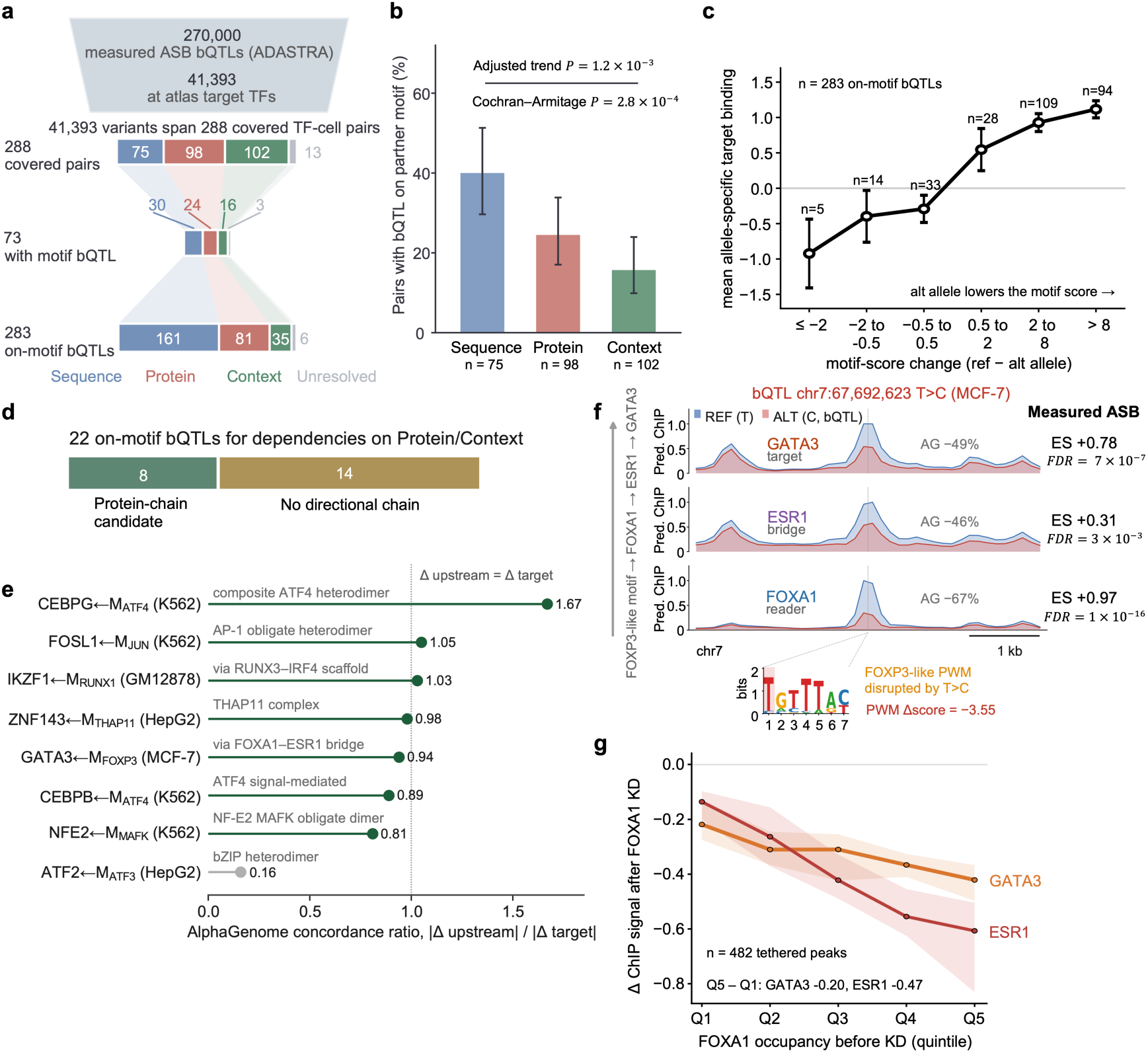
Allele-specific binding supports the inferred routes and reader chains. **a,** Overlap between ADASTRA allele-specific binding variants and the ARES atlas. Of 288 covered dependencies, 73 contained at least one bQTL within the partner motif. **b**, On-motif bQTL frequency declined monotonically across routes (SEQUENCE 40.0%, PROTEIN 24.5%, CONTEXT 15.7%). The ordered trend was significant by a Cochran–Armitage test (P = 2.8 × 10^-4^) and remained significant after adjusting for motif width and bQTL density in a logistic regression with route entered as an ordinal score (adjusted trend P = 1.2 × 10^-3^). Error bars indicate 95% confidence intervals. **c**, Stronger disruption of the partner motif was associated with larger allele-specific target TF binding effects. Points show the mean allele-specific target-binding effect across 283 on-motif bQTLs, binned by motif-score change between alleles; both axes are oriented reference-minus-alternative, so a positive value indicates that the reference allele has the higher motif score (x) or the stronger target binding (y). A variant that lowers the motif score (moving right) is thus associated with reduced binding on the same allele. Error bars indicate s.e.m. Dose- dependence held where the motif is read as a binding site but not in CONTEXT-route dependencies (per-route Spearman calculated from the unbinned variants: SEQUENCE r = 0.39, P = 2.8 × 10^-7^, n = 161; PROTEIN r = 0.41, P = 1.3 × 10^-4^, n = 81; CONTEXT r = 0.12, P = 0.51, n = 35). **d**, Evidence-tier classification of 22 on-motif bQTLs, one per PROTEIN- or CONTEXT-route dependency. In eight cases the report nominated a directional protein-chain intermediate (a reader, bridge or heterodimeric partner) whose allele response could be traced; these were evaluated in **e**. The remaining 14 were consistent with co-occupancy without a direct protein interaction, and did not constitute a testable directional chain. **e**, AlphaGenome concordance for the eight protein-chain bQTL candidates from **d**. For each candidate, AlphaGenome was used to predict how the motif-altering variant changes the binding of every factor in the ARES-nominated chain (the reader or cofactor and the downstream target). The concordance ratio is the absolute predicted change in binding at the weakest (limiting) upstream node divided by the predicted change at the target; a ratio near 1 indicates the perturbation propagates through the chain to the target, whereas a low ratio indicates it does not. Seven candidates showed concordant intermediate–target effects (ratios > 0.8); the exception fell at 0.16, with no case in between. Because both quantities derive from the same predictive model, this is an in silico consistency check, not independent validation. **f**, A FOXP3-like dependency in MCF-7 in which the variant is itself an allele-specific binding QTL at every node of the ARES-inferred chain. The bQTL chr7:67,692,623 T>C disrupts the FOXP3-like partner motif (PWM Δscore = −3.55) and shows measured allele-specific binding (ADASTRA) at FOXA1 (reader; ES = +0.97, FDR = 1 × 10^-16^), ESR1 (bridge; ES = +0.31, FDR = 3 × 10^-3^) and GATA3 (target; ES = +0.78, FDR = 7 × 10^-7^), each preferring the reference allele and thus losing binding as the motif is weakened. Shaded tracks show AlphaGenome-predicted REF and ALT occupancy (AG % = predicted change, alt vs ref); ES and P report the measured ADASTRA allele effect. Both indicate reduced binding on the motif-disrupting allele, consistent with coordinated allele-sensitive occupancy across the ARES-nominated assembly. This was the only protein-route chain whose intermediate nodes carried allele-specific binding data; each track is normalized to its own maximum. **g**, Change in ChIP signal after FOXA1 knockdown in T-47D at the 482 inferred tethered loci, binned into quintiles of FOXA1 occupancy before knockdown. Lines, per-quintile medians; bands, 95% bootstrap intervals. GATA3 was profiled as an HA-tagged protein; the empty-vector anti-HA control is shown in Supplementary Fig. 7.

We next asked whether the magnitude of motif disruption predicted the magnitude of allele-specific binding. If partner motifs contribute directly to target TF binding, stronger disruption of the motif should produce larger binding effects, but only where the motif is read as a binding site. Consistent with this, allele-specific binding tracked the signed change of motif score across 283 on-motif bQTLs (Fig. 5c). This relationship was pronounced where the motif acts as a binding site: SEQUENCE-route (Spearman r = 0.39, P = 2.8 × 10^-7^, n = 161) and PROTEIN-route (r = 0.41, P = 1.3 × 10^-4^, n = 81) dependencies both showed strong, significant correlation. In contrast, CONTEXT-route dependencies showed a substantially weaker correlation that did not reach significance (r = 0.12, P = 0.51, n = 35), consistent with their partner motifs acting as markers of regulatory context rather than as directly read binding sites, though the smaller number of CONTEXT bQTLs limits this comparison. The remaining six on-motif bQTL variants belong to unresolved dependencies and are not analyzed by route.

### ARES nominates mechanistic chains that can be traced and perturbed

Beyond this aggregate route-level pattern, ARES can nominate a specific hidden mechanism for an individual variant, generating a traceable hypothesis. We used AlphaGenome^15^ to test these mechanistic chains and first calibrated against the measured allele-specific binding effects. Of the 73 on-motif dependencies, 67 had a target TF with AlphaGenome coverage, together spanning 273 bQTL variants; across these variants, AlphaGenome-predicted and measured allele effects agreed in direction for 236 of 273 variants (86%; Spearman r = 0.64; Supplementary Fig. 6a). At the dependency level, where each dependency is represented by the most significant variant, 60 of the 67 (90%) were direction-concordant and were retained.

This calibration applied only to the chain-level analysis here; the dosage relationship above used all 283 variants without AlphaGenome. Of the 60 calibrated dependencies, 32 fell in PROTEIN or CONTEXT route; SEQUENCE-route dependencies were excluded because direct sequence recognition involves no trans-acting chain to test. For 22 of the 32, AlphaGenome covered the chain components (the target plus at least one ARES-named reader or cofactor; see filtering steps in Supplementary Fig. 6b). We classified these 22 by the interpretation their ARES report provided (Fig. 5d and Supplementary Table S5): eight nominated a directional protein-chain intermediate, a reader, bridge or heterodimeric partner, whose allele response could be traced; the remaining 14 were consistent with co-occupancy without a direct protein interaction, and did not constitute a testable directional chain.

For the eight protein-chain hypotheses, we asked whether the predicted allele effect propagated through the ARES-named intermediate, quantifying the predicted change at the upstream node relative to the target (Fig. 5e). The cases separated cleanly: seven showed concordant allele effects (ratios > 0.8), while the exception (an ATF2–ATF3 bZIP pair) fell at 0.16, with no case in between, so the distinction does not depend on the choice of cutoff.

In one case, every node of an inferred chain independently carried allele-specific binding data, allowing the full chain to be tested against measured genetics rather than prediction alone (Fig. 5f). In MCF-7, ARES had identified a forkhead-family motif labelled FOXP3 as a proxy for FOXA1 occupancy and proposed that GATA3 is recruited to FOXA1-bound loci by protein tethering where a canonical GATA3 motif is absent, with ESR1 as a candidate component of this assembly. A variant disrupting this motif is itself a binding QTL at all three nodes of the ARES-inferred mechanistic chain: the reader FOXA1, the bridge ESR1 and the target GATA3 each lose binding on the motif-weakening allele (all FDR < 0.01; Fig. 5f). Although the motif is named for FOXP3, the allele-specific binding data supported FOXA1 as the functional reader in this inferred chain. This was the only chain whose intermediate nodes carried allele-specific binding data, and, where such coverage existed, it supported the ARES interpretation directly, without recourse to a predictive model.

To establish the acting order from FOXA1 through ESR1 to GATA3, we used FOXA1-knockdown ChIP-seq in T-47D,^71,72^ a luminal breast cancer cell line different from the original MCF-7 in which ARES conducted inference. Considering FOXA1-overlapping GATA3 peaks with the partner motif but not canonical GATA3 motif (n = 482), both GATA3 and ESR1 binding declined upon FOXA1 knockdown, proportional to the prior FOXA1 binding (Q5 − Q1: GATA3 −0.20, ESR1 −0.47; Fig. 5g). ESR1 showed greater dependency on FOXA1 than GATA3, consistent with its established reliance on FOXA1 for chromatin access. Because GATA3 was profiled as an HA-tagged protein, we ensured that this gradient was not an artifact: empty vectors showed low anti-HA signal at the same loci (15.3% of HA-GATA3 signals), with uniform decrease upon FOXA1 knockdown rather than in proportion to prior FOXA1 binding, and subtracting empty vector signals did not weaken the GATA3 trend (Supplementary Fig. 7). Occupancy of both downstream nodes at these loci therefore showed a FOXA1 binding-dependent decline.

## Discussion

Transcription factor cooperation has largely been studied one partnership at a time, as a catalog of interactions established in individual systems. By inferring mechanisms for 1,552 TF-motif dependencies across ten cell types, we find that cooperation does not fall into rigid categories but varies along a continuum that can be summarized by a small number of recurrent mechanistic routes. A consistent principle emerges: a partner motif acts less often as a fixed regulatory instruction than as an address. The sequence helps specify where a protein can bind; the protein that reads the motif helps determine what regulatory action follows in trans. Because the reader is frequently not the factor for which the motif is named, the same sequence feature can carry different regulatory consequences in different cellular contexts.

This view helps organize observations previously described case by case. The regulatory output of a binding site is known to depend on context, including binding site number, neighboring factors, promoter architecture and chromatin state, and a factor’s genomic occupancy can vary between cell types. Our atlas resolves these principles systematically by separating three quantities that are often conflated: motif identity, reader identity and regulatory output. Similar motifs tended to mark similar regulatory regions, whereas shared reader identity was more closely associated with shared transcriptional outcome. Thus, the advance is not simply that TFs can be bifunctional or that motif effects are context dependent, but that the regulatory consequence of a motif tracks the identity of the protein reading it more closely than the motif label does itself. This reframes motif annotation from a one-to-one mapping between sequence and named factor to a context-dependent mapping among sequence, reader and regulatory action.

The strongest test of a reader assignment is to remove the reader, and two kinds of data do this at different scales. At the level of occupancy, public perturbation experiments existed for two dependencies. In K562, knockdown of GATA1, inferred as the reader of a GATA5-labelled motif, reduced TEAD4 occupancy in proportion to how much GATA1 had bound at each site, while CTCF at the same loci was unaffected. In T-47D, knockdown of FOXA1, inferred as the reader of a FOXP3-labelled motif, reduced both GATA3 and ESR1 occupancy at the loci that motif marks, again in proportion to prior FOXA1 binding. At the level of transcriptional output, single-cell perturbation across 50 dependencies and 19 readers showed that knocking down an inferred reader reproduced the direction of knocking down the target TF more often than knockdown of control factors matched for occupancy at the same peaks. The two lines of evidence are complementary: the knockdown experiments carry internal controls and site-level resolution, the single-cell data carry breadth across readers. A systematic search of public repositories recovered no further experiments pairing knockdown of an inferred reader with ChIP-seq of its target in a matched cell type.

ARES also illustrates a general strategy for moving from predictive sequence features to mechanistic hypotheses. Rather than interpreting a model only by inspecting its internal representation, ARES begins from a predictive dependency and asks what molecular path could explain it. It generates competing explanations, tests them against independent genomic and biochemical evidence, and rejects those that fail prespecified checks. The reliability of this process comes not from the large language model but from the surrounding discipline: fixed analysis routines, logged computation, deterministic gates, calibrated evidence thresholds and explicit abstention when the available data cannot support a mechanism. These safeguards address a specific failure mode of agentic analysis, in which reported quantitative claims are fabricated but not based on analyses actually executed. Against a one-shot agentic baseline, this design eliminated the confirmed fabrications observed in that baseline and produced more commensurable evidence across dependencies. More broadly, ARES suggests how agentic systems can be useful in biology when they are required to convert predictions into falsifiable, auditable, testable statements rather than free-form explanations.

A reader-centered view of regulatory motifs has implications for sequence-to-function models. Such models have become increasingly accurate at identifying motifs and predicting regulatory activity, yet a motif’s effect may depend on which proteins are available to read it in a given cell.

Our results suggest that some limits of cross-cell-type generalization may arise not only from insufficient training coverage, but also because the regulatory consequence of a motif is not fully specified by local sequence alone. Sequence encodes binding potential, grammar and affinity, whereas the cellular repertoire of readers helps determine what action follows. Models that represent reader availability or reader identity explicitly may therefore generalize differently from models that learn these effects implicitly from broader data. Testing this hypothesis will require showing that reader information improves prediction of sequence-to-function models across cell types, perturbations or variants beyond sequence-only baselines.

Approximately one quarter of dependencies remained unresolved because the available evidence did not support a single mechanistic route with sufficient confidence. This category records evidential limitation rather than an absence of biological mechanism: resolution was likely constrained in many cases by incomplete cell-matched coverage of candidate readers, cofactors, interactions or perturbations, although some dependencies may reflect genuinely mixed mechanisms or processes outside the present hypothesis space. By retaining these cases as unresolved, ARES avoids forcing unsupported explanations and identifies where additional data would be most informative.

Although developed here for transcription factor binding, ARES addresses a broader class of problems in which a predictive association is biologically informative but mechanistically unresolved. Our bQTL analysis provides one example of this extension: ARES connected motif-disrupting variants with route-specific allele effects on target binding and, where evidence and assay coverage permitted, generated explicit reader–cofactor–target hypotheses for individual variants. The present analysis was deliberately restricted to bQTLs overlapping the partner motif under investigation, allowing each variant to act as a direct perturbation of the predictive sequence feature. In principle, the same hypothesis-testing framework could be applied more generally to molecular QTL, GWAS loci or model-predicted regulatory variants by beginning from a variant–phenotype association and evaluating alternative paths through TF binding, chromatin state, regulatory contacts and downstream gene expression. In these settings, ARES would act as an additional interpretive layer that converts a predictive association into explicit, auditable mechanistic hypotheses supported by the available evidence, while identifying the experiments needed to distinguish, refine or validate them.

## Methods

### Data sources and preprocessing

Analyses were performed across ten human cell types: A549, GM12878, H1, HEK293, HeLa-S3, HepG2, K562, MCF-7, SK-N-SH and WTC11. All genomic coordinates were in hg38 and analyses were restricted to chromosomes 1–22 and X. Primary functional-genomic data were obtained from ENCODE4.^33^ TF ChIP-seq datasets were used as released narrowPeak peak calls together with fold-change-over-control bigWig signal tracks. Additional ENCODE tracks used by ARES included histone ChIP-seq, chromatin accessibility, RNA-seq, WGBS, and Hi-C contacts; chromatin-state segmentations were the Roadmap Epigenomics 18-state core model lifted to hg38.^73^ Conservation scores were obtained from the UCSC hg38 100-way phyloP track.^47^ Protein–protein interactions were obtained from STRING v12.0.^48^ Gene models were from GENCODE v38.^74^ Motif analyses used the JASPAR 2022 CORE vertebrate non-redundant v2 collection^34^ for partner-motif discovery and a combined JASPAR 2022, HOCOMOCO v12^75^ and CIS-BP^9^ motif database for downstream motif assignment and centering analyses.

For TF ChIP-seq peak processing, each ENCODE narrowPeak was represented by a fixed 200-bp window centered on the peak midpoint (±100 bp). Peaks were filtered to the analyzed chromosomes and peaks wider than 4 kb were removed; windows extending beyond contig boundaries or containing ambiguous bases were excluded, as were peaks with all-missing bigWig signal. All passing peaks were retained, without any top-N or signal-value threshold. For random-forest modelling, the response for each peak was log1p of the mean fold-change-over-control signal across the 200-bp window, extracted from the corresponding bigWig track.

### Partner-motif discovery

Each 200-bp target-TF peak (Data sources and preprocessing) was scored against all 841 motifs in the JASPAR 2022 CORE vertebrate non-redundant v2 collection, using the best-hit PWM log-odds convolution score over all window positions and both strands (Supplementary Methods), giving an 841-dimensional motif-score vector per peak. For each target TF and cell line, a random-forest regressor (scikit-learn; 300 trees, max_features = 0.3, seed 1337) predicted the log1p-transformed target ChIP-seq signal from these scores, with 10% of peaks held out for evaluation (full partition in Supplementary Methods).

Motif importance was quantified as the mean absolute TreeSHAP value computed over all peaks of the dataset, subsampled to a fixed 1,000 (seed 42) where larger, and motifs were ranked accordingly. Because importances use all peaks rather than the training partition alone, the motif ranking that defines the partner motif does not depend on the train/test split. To avoid rediscovering the target’s own recognition motif, self and paralogous motifs were removed before ranking: a motif was treated as self when its TF name matched the target, including the individual components of heterodimeric motifs (denoted ’::’), and as paralogous when the target and motif TF shared a family annotation in the JASPAR 2022 CORE vertebrate collection.

Because family annotations are keyed to the 841 motifs in that collection, the paralogue filter is inactive for targets absent from this set (731 of 1,552 dependencies; 727 of these have no JASPAR motif at all, and 4 appear only as components of heterodimeric motifs); the self filter matches the target’s name against motif components and does not depend on these annotations. The residual risk for these 731 is therefore a partner motif belonging to an unannotated factor with shared specificity, which cannot be excluded a priori. The proportion of target-as-reader assignments among 1,146 route-resolved dependencies was nevertheless similar between annotated and unannotated targets (18.2% versus 19.9%; Fisher’s exact P = 0.50), and a name-root audit of the 731 found no partner motif sharing a root with its target outside zinc-finger nomenclature (all 45 shared roots were ZNF or ZBTB, prefixes common to several hundred unrelated genes). The highest-ranking remaining motif was the partner motif for that target TF and cell line, yielding one candidate dependency per ChIP-seq dataset. Figure 1b compares these ranks across the 825 datasets whose target is named in the collection, directly or as a component of a heterodimeric motif.

### Partner-dependence coefficient

To quantify the extent to which a partner motif’s predictive effect on target-TF occupancy depends on binding of the motif-namesake TF, we fitted, for each TF-motif dependency in the cellular context, an ordinary least-squares model of target-TF ChIP signal as a function of partner-motif strength, motif-namesake-TF occupancy and their pairwise interaction.

Target-TF peaks were processed identically to the partner-motif discovery pipeline: each ENCODE narrowPeak was represented by a fixed 200-bp window centered on the peak midpoint, and peaks were discarded if the window extended beyond the contig, contained an ambiguous base, or had all-missing bigWig signal. All passing peaks were retained, without any top-N or signal-value threshold. The response (*y*) was log1p of the mean fold-change-over-control ChIP signal across the 200-bp window at each of the target’s peaks; the observations for each model were therefore the full set of the target factor’s own peaks. Partner-motif strength (*M*) was the best-hit partner-motif score at each target peak, computed as the maximum position-weight-matrix log-odds convolution score over all window positions and both strands (Supplementary Methods). Motif-namesake-TF occupancy (*P*) was a binary indicator of whether the target’s 200-bp window overlapped any of the motif-namesake factor’s 200-bp windows, defined by the same midpoint-centered procedure. The partner-dependence coefficient is therefore a descriptive quantity: it measures how the motif’s predictive effect depends on the presence of the namesake factor, and does not presuppose that the namesake is the protein that reads the motif. A coefficient near zero is itself informative on that question, indicating that the motif’s predictive information does not require the namesake factor to be bound. The model was

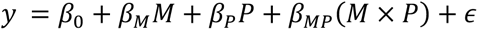

and the partner-dependence coefficient we report is βMP, which compares how much target binding increases per unit of partner-motif score, at peaks where the motif-namesake is bound versus where it is absent. Positive values indicate that the motif becomes more predictive of target occupancy when the namesake TF is bound, values near zero indicate a namesake-independent motif effect, and negative values indicate a reduced motif effect at namesake-bound peaks.

To make partner-dependence coefficients comparable across dependencies, we standardized the response and the motif-strength predictor within each dependency (z-scored y and M), retaining P as a binary variable, and refitted the model; the reported coefficient (Fig. 1c) is β_MP_ from this standardized fit. The significance of each motif-namesake dependence (for example Fig. 1d,e) was assessed by a two-sided t-test on β_MP_ in the joint model; because this test is scale-invariant, it is unaffected by standardization.

Partner-dependence coefficients require an occupancy indicator for the motif-namesake TF and are therefore defined only where that TF has its own ChIP-seq in the same cell line. Of the 1,552 dependencies, 847 lacked motif-namesake-TF ChIP-seq. A further 52 were excluded because the partner motif alone explained less than 0.5% of the variance in target TF signal (R² < 0.005), leaving the interaction term poorly identified, and 4 because the motif-namesake TF occupied fewer than 10 or more than n − 10 of a target TF’s n peaks, so that the occupancy indicator was near-constant. Coefficients are reported for the remaining 649.

To test whether the partner-dependence coefficient partitioned dependencies into discrete quantitative classes or formed a continuous spectrum, we applied Hartigan’s dip test. Because a _small_ number of partner motifs recur across many dependencies (REST, 181 of 649; YY1, 102), the per-dependency distribution is subject to pseudo-replication: applied to all 649 coefficients, the dip test was borderline (D = 0.021, P = 0.055) and a Gaussian-mixture model favored two components (ΔBIC = 66). This secondary component was dominated by a single recurrent motif rather than representing an independent class: 177 of its 245 dependencies (72%) were REST-associated, carrying consistently positive coefficients consistent with REST’s tethering-based, protein-dependent mechanism. Collapsing each partner motif to its median coefficient (*n* = 117) removed this pseudo-replication and yielded an unambiguously unimodal distribution (dip test P = 0.99; a single Gaussian preferred by BIC; Sarle’s bimodality coefficient 0.22). Collapsing reduces the number of observations and therefore the power of the dip test, but the same conclusion is supported by BIC and by the bimodality coefficient, neither of which depends on a significance threshold. We therefore describe the partner-dependence coefficient as a continuous spectrum rather than a set of discrete classes.

### ARES agentic mechanism inference

ARES was designed to convert each (target TF, partner motif, cell type) dependency into a mechanistic report explaining why the partner motif predicted target binding in the cellular context. Each ARES run was initialized with a target TF, partner motif, cell line, a statement of the motif-prediction finding and a structured data manifest listing the available genomic resources for that cell line. The manifest included TF and histone ChIP-seq peaks and signal tracks, chromatin accessibility, DNA methylation, ChromHMM states, RNA-seq, evolutionary conservation, Hi-C contacts, motif scans and STRING protein–protein interaction data.

ARES used an iterative multi-agent workflow whose defining principle is that language-model judgments are constrained by deterministic checks that can override them: agents propose hypotheses, code and interpretations, but faithfulness and termination are enforced by non-model guards rather than left to the model’s discretion. A hypothesis-generation agent (Gemini 3.1 pro) proposed candidate mechanisms; for each proposal it was required to state how the predicted signature would differ from mere co-occupancy and to redesign any test that could not distinguish the two. A review agent (GPT-5.4 mini) then screened each proposed mechanism for novelty, specificity, testability and discriminative value; a coding agent _(MiniMax_ M2.7) generated and executed analysis code in a dedicated computational environment; and a summary agent (Gemini 3.1 pro) evaluated the resulting evidence and assigned each hypothesis an outcome of SUPPORTS, REJECTS, INCONCLUSIVE, UNTESTABLE or ERROR. ERROR outcomes triggered retry rather than biological interpretation. The first hypothesis in each run was that the target’s ChIP-seq signal is a technical artifact, evaluated against the target’s expression and the enrichment of its own motif at its peaks.

Every executed code block and its standard output were persisted to disk, and every claim in the final report was evaluated against this logged execution output rather than model memory. Two mechanisms enforced fidelity to that log. At the level of named entities, a deterministic check compared every gene-like token in the summary against the raw execution output and flagged, with a confidence penalty, any entity absent from it—so a fabricated factor, having no log entry to trace to, could not enter the report unmarked even when the model failed to notice. At the level of reported numbers, the coding agent was required to re-derive every statistic in a final verification cell before emitting a solution, checking directionality, hardcoded constants and _contingency_ tables against the underlying variables. Measurement operations that recur across the atlas—motif scanning and peak overlap—were routed through shared helper routines with pinned parameters (FIMO at P < 1×10^-4^; a fixed overlap denominator), which the coding contract required in place of hand-rolled alternatives, so that quantities were computed under common definitions across all dependencies rather than under per-run choices.

After each tested hypothesis, ARES evaluated whether the run had converged, but the summary agent’s judgment was not final. A deterministic convergence check, evaluated in a context independent of the reasoning that produced the hypotheses, verified that at least one hypothesis (excluding the iteration-0 artifact check) had reached SUPPORTS, that convergence rested on at least two genuinely independent measurement modalities, and that any third-party factor named in a supported mechanism had itself been tested; if these conditions were unmet, a claimed convergence was overridden and the run continued. The hypothesis agent was not permitted to declare convergence at all—any such call was rewritten as a new hypothesis, reserving termination for the guarded summary step. Where the reported narrative and the recorded outcomes disagreed, an authoritative outcome table superseded the prose. For the artifact check specifically, a claimed SUPPORTS was clamped to INCONCLUSIVE when expression data were absent from the manifest, preventing a call resting on self-motif evidence alone. Runs stopped at guarded convergence, after repeated rejected hypotheses, or after a maximum of 15 iterations.

Each run produced a final report written to nine required sections (executive summary, methodology, key findings, evidence summary, mechanism comparison, speculative biological purpose, confidence, limitations and recommendations) and an investigation narrative recording every iteration: the hypothesis and its causal chain, the prediction and how it was made distinguishable from co-occupancy, the verification plan, the outcome with its confidence, and the reasoning for the next step. The code and standard output of every executed block, together with all intermediate code, were retained alongside them.

ARES reports were based only on the functional-genomic evidence supplied in the manifest. MPRA, CAP-SELEX, Perturb-seq, ADASTRA bQTLs, AlphaGenome predictions and the reader-perturbation ChIP-seq datasets were not used during ARES report generation or route assignment, and were used only as post hoc validations.

### Route definitions and report extraction

ARES reports were converted into structured annotations using a two-stage extraction procedure. The first stage assigned each dependency to one of five broad classes: SEQUENCE, PROTEIN, CONTEXT, UNRESOLVED or ARTIFACT. SEQUENCE was defined as a mechanism in which the predictive signal is carried by the DNA sequence or motif grammar itself, including direct recognition by the target, embedded motifs, homotypic clustering, or composite grammar. PROTEIN was defined as a mechanism in which a separate protein reads the partner motif and acts directly on the target by recruitment, tethering, cooperative co-binding, stabilization or exclusion. CONTEXT was defined as a mechanism in which the partner motif marks or shapes a chromatin or genomic context, such as accessibility, promoter state, repressive chromatin, nucleosome positioning, HOT-region co-occupancy, three-dimensional architecture or a DNA-intrinsic sequence feature (GC/CpG content, DNA shape), to which the target responds. UNRESOLVED was used when no mechanism was established, and ARTIFACT when the report indicated that the partner motif did not support a genuine dependency.

Route extraction used a trace prompt that followed the mechanistic path from partner motif to reader, actor and target (Supplementary Table S2). Each report was processed five independent times by the route extractor. The final route was assigned by majority vote; tied votes were assigned UNRESOLVED. Deterministic guards demoted calls to CONTEXT when the relevant actor was recruited by chromatin context rather than a motif-reader chain, and to UNRESOLVED when no report-supported mechanism was present. In addition to the broad route, the extractor recorded the inferred motif reader, actor, reader category, regulatory region and transcriptional outcome.

The motif reader was inferred using a route-conditioned majority procedure. Only extractor votes matching the final route were considered, and reader strings were parsed into individual proteins with expansion of shorthand and heterodimer notation. A reader was retained when it appeared in more than half of same-route votes. Readers were classified as the motif-namesake TF, the target itself, a hidden third factor or unassigned. Dimer-subunit corrections reassigned cases in which the extracted reader was a subunit of the named partner motif, such as AP-1 or NF-Y heterodimer components, from hidden reader to partner reader.

The second extraction stage assigned a specific mechanism leaf within the selected route. Five independent mechanism-extraction votes were generated for each report. A leaf was assigned when a unique non-abstaining leaf received at least two votes; otherwise the dependency was flagged for curator review. Mechanism leaves included direct sequence recognition and motif grammar for SEQUENCE; tethering, cooperative co-binding, co-occupancy and competitive exclusion for PROTEIN; and chromatin state, sequence-feature proxy, surrogate TF co-occupancy, accessibility proxy, HOT-region co-occupancy and 3D-loop context for CONTEXT (Supplementary Table S3).

Extractor performance was evaluated against blind curator labels. The curator reviewed 72 held-out reports, of which 70 received a confident SEQUENCE, PROTEIN or CONTEXT label; two could not be confidently classified and were excluded. Route assignment matched the curator label in 66 of 70 cases (94%; Cohen’s κ = 0.91; Supplementary Fig. 8b). The four discordances were one SEQUENCE to PROTEIN and three CONTEXT to PROTEIN calls. The final atlas comprised 1,552 dependencies across 10 cell lines, including 300 SEQUENCE, 363 PROTEIN, 483 CONTEXT, 391 UNRESOLVED and 15 ARTIFACT dependencies (Supplementary Table S7).

### Benchmark against a single-agent baseline

**Benchmark set.** Fifty dependencies were drawn from the atlas, ten per route class (SEQUENCE, PROTEIN, CONTEXT, UNRESOLVED and ARTIFACT). Within each class, dependencies were sampled randomly from the atlas under a constraint maximizing the number of distinct cell lines represented.

**Task specification.** Both systems received the same task file, which stated only the finding to be explained (“a systematic screen of 841 JASPAR PWMs identifies the {partner} motif score as the dominant feature for predicting the continuous signal intensity of {target} binding in {cell}”), the investigation objective, the availability of the full multi-omic dataset for that cell line, and the nine required report sections. It named neither the route taxonomy nor the pipeline, so the expected answer could not be recovered from the prompt.

**Baseline agent.** The baseline was the OpenAI Codex CLI v0.133.0 running GPT-5.5 at medium reasoning effort, invoked once per dependency with the same initial prompt as ARES and with no orchestration loop. It was granted access to every data file ARES receives, and could execute tools in the same coding environment as ARES. Each invocation used an isolated agent state directory, so no context carried between dependencies. The baseline agent was additionally instructed to save all executed code and output, a step ARES performs automatically; unverifiable classifications therefore reflect non-compliance with an explicit instruction rather than the absence of one.

**Method commensurability.** From the saved code of each report we extracted the motif-calling method and threshold, the peak-overlap window and the signal-averaging window actually used, recording file-and-line evidence for every value.

**Artifact designation and abstention.** Artifact status was established independently of both systems from two measured quantities per dependency, with no report consulted: the target’s expression in its own cell line (ENCODE RSEM quantification mapped through GENCODE), and enrichment of the target’s cognate motif within its own peaks, computed as the fraction of the top 1,500 peaks (summit ± 100 bp) carrying at least one FIMO hit at P < 10^-4^ divided by the same fraction in per-sequence dinucleotide-shuffled sequence. A dependency was designated an artifact when the target was effectively unexpressed, had no cognate motif in the motif database, or showed no enrichment of its cognate motif over the shuffled null. The ten artifact dependencies had target expression of 0.00–0.41 TPM (median 0.11), compared with a median of 16.6 TPM across the remaining forty. Of the ten, three had no cognate motif in the database and four showed cognate-motif enrichment near or below the shuffled null (0.37–1.19-fold).

Because all ten fell below the 0.5 TPM threshold of the ARES expression gate, ARES abstention on this set follows by construction, and the comparison is informative for the baseline, which had no artifact gate. Each report was then read by a human evaluator and classified by whether it declined to assert a mechanism (ABSTAIN), asserted one (ASSERT_MECHANISM), or diagnosed the artifact while nevertheless advancing a mechanistic account (PARTIAL).

**Faithfulness audit.** Every statistic in all 50 baseline reports and all 50 ARES reports was checked against the analysis scripts and output files saved in the report’s directory, under an identical two-stage procedure for both systems. Reports from both systems were pooled, stripped of source and infrastructure identifiers, relabeled with opaque identifiers, and the mapping was sealed until auditing was complete. A single auditing agent (Claude Opus 4.6), blind to provenance, received each report with its saved files and identified claims not supported by a saved computation. Every flagged claim was then manually verified against the saved code and outputs, and only issues confirmed at that stage were counted. Confirmed issues were classified as major where a reported number or analysis had no corresponding computation, where a number was absent from the saved outputs, where the reported direction contradicted the saved output, or where a result was attributed to the wrong entity; as minor where imprecise wording did not affect the conclusion; as unsupported where a number was untraceable but not conclusion-driving; and as unverifiable where neither code nor outputs were saved.

### Route mapping onto the partner-dependence coefficient

To relate ARES routes to the quantitative continuum (Fig. 2a), we compared the partner-dependence coefficient across routes among the dependencies with defined coefficients.

Because a small number of partner motifs recur across many dependencies, route-level comparisons could be dominated by hub-motif pseudo-replication; we further collapsed each partner motif to its median coefficient within each route before testing (n = 42 SEQUENCE-, 56 CONTEXT- and 40 PROTEIN-route motif representatives; Supplementary Fig. 2a). PROTEIN-route dependencies had significantly higher partner-dependence coefficients than both SEQUENCE (P = 0.020) and CONTEXT (P = 0.028) dependencies by a two-sided Mann–Whitney U test. SEQUENCE- and CONTEXT-route dependencies occupied the lower, protein-independent range and were not separated on this axis, consistent with the partner-dependence coefficient measuring dependence on motif-namesake-TF occupancy specifically rather than the full mechanistic distinction.

### Cross-cell reproducibility

To assess whether ARES route assignments capture reproducible rather than idiosyncratic properties of TF–motif relationships, we evaluated reproducibility across cell types.

We grouped cell-type-specific dependencies by their shared target TF and partner motif, and retained recurrent TF–motif groups observed in more than one cell type. Here, a cell-type-specific dependency observation refers to a unique (target TF, partner motif, cell type) tuple, whereas a recurrent TF–motif group refers to the set of observations sharing the same target TF and partner motif across cell types. Concordance was defined as agreement of the extracted route (SEQUENCE, PROTEIN or CONTEXT) between two observations from the same recurrent TF–motif group in different cell types.

A recurrent TF–motif group observed in k cell types contributed k observations and in theory C(k, 2) pairwise cross-cell comparisons. Because a TF-partner-motif group does not occur in each cell type, 90 recurrent TF–partner-motif groups yielded 219 cell-type-specific dependency observations and 188 pairwise cross-cell comparisons. Concordance was computed over the 188 pairwise comparisons.

To test whether concordance exceeded chance, route labels were permuted across the 219 observations while holding the assignment of observations to recurrent TF–motif groups fixed, and concordance was recomputed over the same 188 comparison set. This procedure was repeated 10^6^ times. The observed concordance was 67.0%, compared with a permutation mean of 34.4% (z = 9.5). No permutation reached or exceeded the observed concordance. Using the add-one estimator, (b + 1)/(m + 1), where b is the number of permutations with concordance at least as large as observed and m is the number of permutations, this corresponds to P = 9.99 × 10^-7^, which we report as P < 1 × 10^-6^ rather than relying on a parametric approximation to the tail of the null distribution.

We performed two robustness analyses. First, because the global permutation does not preserve cell-type-specific route frequencies, which varied appreciably across cell types (the fraction of PROTEIN-route observations ranged from 25% in HEK293 to 58% in HeLa-S3), we repeated the analysis by permuting route labels within each cell type separately. This gave a nearly identical null distribution (mean = 34.8%, z = 9.2, P < 1 × 10^-6^). Second, because pairwise cross-cell comparisons are not independent and the nine recurrent TF–motif groups observed in four or more cell types supplied 39% of all comparisons, we repeated the analysis after weighting each recurrent group equally. Specifically, concordance was first computed within each recurrent TF–motif group and then averaged across groups. This group-level concordance was 61.6%, compared with a null mean of 34.4% (z = 6.2, P < 1 × 10^-6^).

### Recurrent motif aggregation

To characterize recurring regulatory programs, dependencies were grouped by partner motif within each cell type. For every motif–cell-line combination, the fraction of dependencies assigned to each route and mechanism leaf was calculated. Motifs represented by at least 15 resolved dependencies were designated recurrent regulatory hubs. Route composition was visualized as stacked proportions across all associated target factors.

### MPRA validation of partner-motif cis-activity

We tested whether partner motifs contribute to local cis-regulatory activity using the PARM massively parallel reporter assay (MPRA; GEO accession GSE301246).^53^ PARM measured promoter-fragment activity across multiple cell lines and six sequence splits, consisting of five cross-validation folds and one held-out test split. MPRA data were withheld from ARES and were used neither during report generation nor during route assignment, so this validation was independent of the ARES dependency calls.

For each dependency, reporter fragments were first restricted to those overlapping the target-TF ENCODE ChIP-seq peaks. Retained fragments were scanned for the partner motif using FIMO from MEME Suite v5.5.5^39^ with the JASPAR 2022 CORE vertebrates non-redundant v2 motif collection at P < 1 × 10^-4^. When the motif-positive rate exceeded 0.5, the threshold was tightened to P < 1 × 10^-5^. Fragments containing at least one partner-motif hit were labelled motif-positive, and fragments without a hit were labelled motif-negative.

Because the six PARM splits are parent-locus-disjoint, they were concatenated within each cell line before testing, avoiding double-counting of promoters across splits. Because PARM tiles each promoter with many overlapping and highly correlated fragments, a fragment-level test would pseudo-replicate promoter-level activity and yield anti-conservative significance estimates. We therefore used parent promoter, rather than fragment, as the sample unit. Within each dependency, motif-positive and motif-negative fragments were balanced by 1:1 nearest-neighbor matching on standardized fragment length, GC content and distance to the transcription start site. Covariates were z-scored on the pooled motif-positive and motif-negative fragment distribution. Matching was performed greedily without replacement, and unmatched fragments were discarded.

Matched fragments were then grouped by parent promoter. For each promoter, we computed the median activity of matched motif-positive fragments minus the median activity of matched motif-negative fragments. These promoter-level motif effects were tested against zero using a two-sided Wilcoxon signed-rank test. A dependency was considered testable only if it contained both motif-positive and motif-negative fragments and passed predefined coverage and promoter-clustering power checks. Dependencies were excluded when they lacked sufficient matched fragments or parent promoters for a promoter-level test, had insufficient MPRA coverage, or had motif prevalence too narrow to support thresholded comparison.

P values were corrected across dependencies using the Benjamini–Hochberg procedure, and dependencies were called significant at q < 0.05. To assess whether significant effects were directionally consistent with the expected regulatory role of the motif-namesake factor, we compared the sign of each significant motif-associated activity effect with a curated conservative annotation of motif-namesake valence as activator or repressor. Partners without a confident valence annotation were left unscored for direction.

The analysis was restricted to dependencies in the canonical ARES atlas and to the four atlas cell lines assayed by PARM: K562, HepG2, HEK293 and MCF-7. Of 989 MPRA-testable dependencies, 358, or 36%, showed a significant partner-motif activity effect. The significant fraction was 42% in HepG2, 37% in HEK293, 33% in MCF-7 and 25% in K562.

### Evolutionary conservation analysis

Evolutionary constraint of partner motifs was quantified using vertebrate phyloP conservation scores (hg38 phyloP100way).^47^ For each dependency, the partner motif was scanned within the target-TF ChIP-seq peaks using FIMO, and the mean phyloP score across the bases of each motif instance was extracted. A per-dependency conservation score was then computed as the mean of these values across all partner-motif occurrences.

Of the 1,552 dependencies in the ARES atlas, the 1,146 assigned to a resolved cooperation route (SEQUENCE, PROTEIN, or CONTEXT; excluding UNRESOLVED and ARTIFACT) were carried into the conservation analysis. A conservation score was obtained for the 1,141 dependencies whose partner motif had at least one scorable instance within the target peaks (SEQUENCE n = 296, PROTEIN n = 363, CONTEXT n = 482), and the 5 dependencies lacking a scorable instance were dropped.

Conservation distributions were compared across cooperation routes using two-sided Mann–Whitney tests. Because conservation information was available to ARES only as supporting evidence and was not used by the route classifier, these analyses provided an independent assessment of the biological constraint associated with each route.

### AlphaGenome motif-perturbation analysis

To evaluate the contribution of partner motifs to target binding, reader binding, CAGE and other tracks, partner motifs were scrambled in silico using AlphaGenome.^15^ AlphaGenome predictions were never used during ARES inference.

For each dependency, target-TF ChIP-seq peaks were scanned for the partner motif with FIMO (P < 1 × 10^-4^) within ±500 bp of the peak summit, and the 50 motif-positive peaks with the highest partner-motif score were retained; dependencies with fewer than five motif-positive peaks were not analyzed. At each retained peak, the partner-motif hit was disrupted by a deterministic base substitution (A↔C, G↔T) that preserved sequence length while altering nucleotide identity, and a matched within-peak control was generated by applying the same substitution 60 bp from the motif. Because the control substitution follows the same rule, any systematic effect of the substitution itself on local base composition is removed by the subtraction below. For each peak, AlphaGenome was queried for the reference and scrambled sequences across multiple molecular outputs, including transcription-factor binding, chromatin accessibility (DNase and ATAC), histone modifications, transcriptional activity (RNA and CAGE) and three-dimensional contact, each restricted to the cell line’s assay ontology.

**Quantification**. AlphaGenome outputs differ in resolution by assay, and signal was extracted over a window centered on the perturbed position in each case. For tracks predicted at 128-bp resolution (transcription-factor ChIP and histone modifications), signal was taken from the single bin containing the perturbed site. For tracks predicted at 1-bp resolution (DNase, ATAC, RNA, CAGE and PRO-cap), signal was averaged over a 129-bp window centered on the perturbed site (±64 bp). For each peak and track, the motif effect was computed as the difference between the two perturbations,

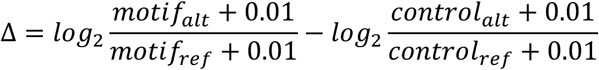

where a pseudocount of 0.01 was added to all predicted values and *ref* denotes the unperturbed prediction in the corresponding extraction window.

**Reader binding versus control factors.** Every three-route-resolved hidden-reader dependency whose reader had an AlphaGenome ChIP-TF track in the matching cell line was included, with no functional filter and no prior selection, yielding 180 dependencies. Where a reader comprised several factors (e.g. FOXA1/FOXA2/FOXA3), the reader value was the mean across that reader’s tracks. For each dependency, the same quantity was computed for twelve control TFs drawn without replacement from that cell line’s AlphaGenome coverage, excluding the target, the namesake partner, the reader and any factor sharing a DNA-binding-domain family with them. Controls were evaluated at the identical loci and bins as the reader, and the control value was the median across the twelve factors. Both are ChIP-TF tracks, so both are the 128-bp quantity defined above. Readers and their control medians were compared by one-sided Wilcoxon signed-rank test on the paired values.

**Reader function and transcriptional response.** This analysis additionally required a directional reader, annotated as an activator or a repressor from Gene Ontology molecular function terms independently of ARES. Human protein annotations to GO:0001228 (DNA-binding transcription activator activity, RNA polymerase II-specific) and GO:0001227 (the corresponding repressor term), including descendant terms, were retrieved from QuickGO.^76^ A reader annotated to both terms and multi-factor readers (ARES inferred more than one protein as motif reader candidate) whose factors disagreed were excluded because they make no directional prediction. This left 78 dependencies, 58 with an activator reader and 20 with a repressor reader. Predicted CAGE change was compared between the two groups by one-sided Mann–Whitney test.

### CAP-SELEX validation

To assess whether PROTEIN-route dependencies exhibit the biochemical cooperativity expected of protein-mediated interactions, we compared atlas dependencies against a published CAP-SELEX compendium of cooperative transcription-factor pairs.^17^ CAP-SELEX measures cooperative DNA binding of purified proteins in vitro and was not used during ARES report generation or route assignment.

An atlas dependency was considered testable when both the target TF and the TF owning the partner motif, including resolved heterodimer components, were present in the CAP-SELEX-tested TF universe (CAP-SELEX constructs together with JASPAR heterodimer components). A dependency was considered CAP-SELEX-supported when the target–partner TF pair, or any of its resolved heterodimer component pairs, appeared in the cooperative-interaction reference set (CAP-SELEX composites and JASPAR heterodimer motifs). Pairs were analyzed per cell line.

Fifty-eight dependencies with a resolved route were testable in vitro (38 PROTEIN, 10 SEQUENCE and 10 CONTEXT). Cooperative support was enriched among PROTEIN-route dependencies (23 of 38, 61%) compared with none of the 20 SEQUENCE or CONTEXT dependencies (Fisher’s exact test, P = 2.1 × 10^-6^). Validation rates for individual PROTEIN-route mechanism leaves were 16 of 23 for cooperative co-binding, 6 of 12 for tethering and 1 of 3 for co-occupancy.

### Reader centering analysis

To test whether inferred readers physically occupied the partner motifs they were assigned to interpret, we quantified reader ChIP-seq signal at motif-centered loci within target-TF peaks. For each dependency, occurrences of the partner motif were identified using FIMO (P < 1 × 10^-4^) within the target TF’s 3,000 highest-signal peaks. The highest-scoring motif occurrence in each peak defined the motif center, and dependencies with fewer than 30 motif-positive target peaks were excluded. Reader ChIP-seq signal was quantified across a ±500-bp window centered on each motif occurrence using 50 bins and averaged across motif-positive peaks to generate a metaprofile.

Two centering metrics were computed from the raw, unscaled metaprofiles. The center-to-flank ratio (CF) was defined as the mean reader signal within ±75 bp of the motif center divided by the mean signal in the 400–500-bp flanks. The summit offset was defined as the distance between the motif center and the position of maximum reader signal. A dependency was classified as centered when CF ≥ 2 and the absolute summit offset was no greater than 50 bp.

To estimate the motif-centered enrichment expected from generic occupancy at active regulatory elements, we constructed a matched random-TF null for each dependency. Four TFs with ChIP-seq data in the same cell type were sampled without replacement, excluding the inferred reader, target TF and partner-motif namesake. Each random TF was quantified at the identical motif-centered loci using the same procedure, and the median of the four random-TF profiles and CF values defined the matched null for that dependency. Reader and matched-null CF values were compared across dependencies using a one-sided paired Wilcoxon signed-rank test.

For display, each reader and matched-null metaprofile was divided by its own mean flank signal, yielding fold enrichment over the 400–500-bp flanks; this rescaling does not change the center-to-flank ratio. Figure 4c shows the median flank-normalized profile and interquartile range across dependencies.

### Reader-target tracking at namesake-unbound motif loci

For each hidden-reader dependency with available target, reader and namesake ChIP-seq in the same cell line, the top 3,000 target ChIP-seq peaks ranked by ENCODE signalValue were scanned for the partner motif using FIMO (P < 1 × 10^-4^). If multiple motif hits occurred in one peak, the highest-scoring hit was retained and its motif center was used as the locus coordinate. Namesake-unbound loci were defined as motif-positive target peaks whose motif center did not overlap a ChIP-seq peak for the motif-namesake TF.

Dependencies were retained if at least 30 namesake-unbound motif loci remained and at least 30 loci had finite target and reader bigWig signals. At each retained locus, target and reader ChIP-seq signal was quantified as the mean bigWig signal in a ±75-bp window centered on the motif. For each dependency, we computed the Spearman correlation between reader and target signal across loci. As a matched null, ten same-cell TFs with available ChIP-seq bigWigs were sampled at random, excluding the target, namesake and reader, and their correlations with the target were computed over the same loci. The median of these ten correlations was used as the null value for that dependency.

Reader correlations were compared with matched null correlations across dependencies by a one-sided paired Wilcoxon signed-rank test. Per-dependency reader-correlation significance, used only for point annotation in the figure, was estimated with a two-sided t approximation to Spearman’s correlation and corrected across dependencies by Benjamini–Hochberg. Reader ChIP-peak overlap was calculated as the fraction of tested motif loci falling inside a reader ChIP-seq peak.

### Perturbation of hidden reader GATA1

**Datasets.** ChIP-seq in K562 under RNAi was obtained from ENCODE^33^: TEAD4 in GATA1 knockdown (ENCSR985RPY) and in MAFF knockdown (ENCSR075FCS), the latter serving as the internal control. Control factors in a GATA1-depleted background were obtained from GEO: TAL1 and H3K27ac from GSE211293^61^, and CTCF from GSE146827^62^. Paired-end reads were processed as single-end using mate 1; the GATA1 knockdown comprised two biological replicates, all other conditions one.

**Peak set.** Analysis was restricted to unperturbed K562 TEAD4 IDR peaks (ENCFF501XJP; 35,971 peaks) containing at least one GATA5 motif (JASPAR MA0766.2; FIMO 5.5.5, --thresh 1e-4; hg38), yielding 8,094 peaks (22.5%). GATA1 occupancy was the mean signal of the unperturbed GATA1 bigWig (ENCFF334KVR) in a 300-bp window centered on the TEAD4 peak; baseline peak strength was the ENCODE signalValue of the same unperturbed TEAD4 peak.

**Alignment and quantification.** Reads were aligned with bowtie2 2.5.5 to hg38 and filtered at *MAPQ ≥ 30* (*-F 1804;* samtools 1.20). Per-peak counts were obtained with bedtools 2.31.1 multicov over the 300-bp window. H3K27ac was quantified over ±1-kb window, because a 300-bp window centered on a transcription-factor summit samples predominantly the nucleosome-depleted region.

**Signal change.** To avoid a peak-strength-dependent artifact from unequal sequencing depth, counts were depth-matched by binomial thinning (each read retained with probability D_min_/D_library_) within each factor to that factor’s smallest library, rather than scaled to CPM alone. The per-peak change was computed as

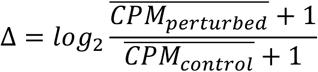

averaging over replicates and over 25 independent thinning draws.

**Dose–response.** Peaks were binned into quintiles of unperturbed GATA1 occupancy (∼1,619 peaks each), and the median Δ within each quintile computed per factor. Bands are 95% percentile intervals from 1,000 bootstraps over peaks. Factors were analyzed separately rather than in a single interaction model, because they were measured in different experiments and the absolute change of each factor after GATA1 knockdown could not be directly compared.

### Perturbation of a second hidden reader FOXA1

**Datasets and peak set.** HA–GATA3 ChIP-seq in T-47D comprised two siNT replicates (SRX5057619 and SRX5057620) and two FOXA1-knockdown replicates (SRX5057621 and SRX5057622), GEO GSE122847.^71^ ESR1 ChIP-seq comprised three control replicates (SRX8861813–SRX8861815) and three FOXA1-knockdown replicates (SRX8861819–SRX8861821), GEO GSE126004.^72^ Four control H3K27ac samples (SRX5323903, SRX5323907, SRX5323911 and SRX5323915) were used to measure baseline regional activity.^72^ Baseline FOXA1 occupancy and FOXA1 peaks were obtained from the ENCODE T-47D tracks ENCFF589DAO and ENCFF758GJL, respectively.

An independent GATA3 peak set was defined by pooling two untreated HA–GATA3 samples (SRX5057609 and SRX5057610) and calling peaks against pooled matched empty-vector anti-HA controls (SRX5057607 and SRX5057608) with MACS2 2.2.9.1 (−q 0.01 --nomodel --extsize 150 --keep-dup all).^77^ This yielded 5,105 GATA3 peaks without using the siNT or siFOXA1 samples subsequently used to quantify perturbation effects.

**Locus definitions.** The canonical GATA3 motif (JASPAR MA0037.4) and FOXP3 motif (JASPAR MA0850.1) were scanned with FIMO 5.5.5 (--thresh 1e-4; hg38) in 300-bp windows centered on GATA3 summits. Tethered loci were GATA3 peaks that overlapped a T-47D FOXA1 peak, contained the FOXP3 motif and lacked the canonical GATA3 motif. Direct loci overlapped a FOXA1 peak and contained the canonical GATA3 motif.

**Signal quantification and dose response.** The single-end T-47D libraries were aligned, filtered, quantified and depth-matched as described above for the K562 analysis. Transcription-factor signals were quantified in 300-bp summit-centered windows, whereas baseline H3K27ac was quantified in 2-kb windows. Perturbation effects were calculated using the same replicate-averaged log2 fold-change with a pseudocount of one and 25 binomial-thinning draws. The 482 tethered loci with both GATA3 and ESR1 perturbation measurements were divided into quintiles of baseline FOXA1 occupancy, measured as the mean ENCFF589DAO signal in the 300-bp window. Median GATA3 and ESR1 changes were calculated within each quintile, with 95% percentile intervals from 1,000 bootstrap resamples over loci. Empty-vector anti-HA ChIP-seq from the same experiment (siNT, SRX5057615–SRX5057616; siFOXA1, SRX5057617–SRX5057618) was processed in parallel as a background QC. EV signal was quantified in the same tethered-locus windows and FOXA1-occupancy quintiles, and the GATA3 dose-response was repeated after EV subtraction.

**Aggregation and weighting.** The aggregate quantities in Supplementary Fig. 7b,c are pooled ratios of summed CPM, not averages of ratios, and two levels of pooling apply. Across loci, CPM were summed within each quintile before the knockdown/control ratio was taken, so loci contribute in proportion to their signal; this is why aggregate gradients exceed the per-locus medians reported in Fig. 5g (Q5 − Q1 = −0.46 versus −0.20 across the same 482 loci). Across replicates, CPM were likewise pooled before the ratio was taken rather than averaging per-replicate ratios, so replicates contribute in proportion to their signal at these loci; the two replicates carry 39% and 61% of the empty-vector signal and 32% and 68% of the HA-GATA3 signal. This second weighting is consequential only for the empty-vector background level, where the replicates disagree in direction (+6.7% and −18.5%) and the pooled value is −8.7% against an unweighted mean of −5.9%. It is negligible for the HA-GATA3 gradient, where both replicates decline (−0.39 and −0.51) and the pooled value differs from the unweighted mean by less than the spread between replicates. Per-replicate values are plotted separately in Supplementary Fig. 7b.

### Pairwise annotation-sharing analysis

To evaluate whether partner-motif identity and reader identity encode distinct aspects of regulatory function, we performed pairwise comparisons across atlas dependencies. The analysis was restricted to PROTEIN- or CONTEXT-route dependencies whose reader is distinct from the partner-motif namesake (reader ≠ namesake). This decoupling is essential: across the full atlas the shared-reader and shared-motif indicators are strongly collinear (r = 0.91), because the reader sometimes is the factor that reads the motif; restricting to reader ≠ namesake reduces this collinearity (r = 0.69) and makes the two effects separately identifiable. The decoupled subset comprised 212 dependencies.

For each pair of dependencies, four binary relationships were recorded: whether they shared at least one reader protein (reader-set overlap), had similar partner motifs (Tomtom comparison of the two partner PWMs, q < 0.05), shared a target factor, and shared a cell type. Two outcomes were evaluated independently: whether the pair shared the same regulatory-region annotation and whether it shared the same transcriptional-outcome annotation, both recorded by ARES. Regulatory-region annotations comprised active enhancer, active promoter, active transcription state, poised enhancer and repressive heterochromatin, corresponding to the internal labels ENHANCER_ACTIVE, PROMOTER_ACTIVE, ACTIVE_TRANSCRIPTION_STATES, ENHANCER_POISED and REPRESSIVE_HETEROCHROMATIN. Transcriptional-outcome annotations comprised activating, repressive and no detectable effect, corresponding to ACTIVATING, REPRESSIVE and NO_EFFECT. After removing dependencies lacking the relevant annotation, 83 dependencies (3,403 pairs) were analyzed for regulatory region and 109 dependencies (5,886 pairs) for transcriptional outcome.

For each outcome, the four relationships were entered together into a single multivariable logistic regression predicting whether the pair shared the annotation, so that each feature’s effect was estimated after adjustment for the other three. Permutation P-values were computed for every feature in both models (the full 4 × 2 grid): the corresponding dependency-level label was permuted while the others were held fixed, the pairwise features were recomputed, and the model was refitted to build a null distribution for the focal coefficient (5,000 permutations per cell). Two associations were prespecified as the focal tests of the dissociation — shared reader in the transcriptional-outcome model and motif similarity in the regulatory-region model — and the remaining cells were exploratory. The shared-target coefficient was quasi-completely separated in both models (too few dyads share a target factor); its bootstrap confidence interval was degenerate and its estimate was not identifiable, so target is omitted from Fig. 4m and not interpreted.

Because both the inferred reader and the transcriptional outcome are produced within ARES, a shared-reader/shared-outcome association (Fig. 4m) could in principle be circular if the outcome were derived from the reader’s function rather than from data. It is not. The transcriptional-outcome annotation was computed from RNA-seq — comparing nearest-gene expression (TPM) at motif-positive versus motif-negative target-TF peaks, with fixed thresholds applied identically across dependencies — and used no information about the identity or canonical valence of the inferred reader, which was assigned from binding and motif data. The two annotations are therefore derived from independent assays. Empirically, the measured outcome frequently contradicted the reader’s canonical function: in 51 of 72 dependencies with a resolvable canonical valence (71%), the outcome was discordant with the reader’s known valence, including activator-family readers assigned REPRESSIVE or NO_EFFECT outcomes (Supplementary Note 4). For example, HEY1/M_GATA5_ (reader GATA2, canonical activators) was predicted to activate but showed significantly reduced expression at motif-positive genes (fold change 0.618, P = 2 × 10^-6^) and was annotated REPRESSIVE. The association in Fig. 4m therefore reflects a genuine relationship between reader identity and measured regulatory output, not a definitional consequence of the annotation procedure.

### Perturb-seq validation of reader assignments

**Perturbation data and knockdown z-scores.** We used the genome-wide CRISPRi Perturb-seq screen in K562 cells,^67^ taking the authors’ normalized pseudobulk expression matrix (perturbations × genes). Genes with an absolute normalized value exceeding 100 in any perturbation were discarded, and genes were further restricted to those whose mean expression exceeded the 10th percentile of all positive means. For each perturbation, the normalized value was converted to a variance-stabilized statistic z = x × √*n*, where n is the number of cells retained for that perturbation; z is used throughout as the signed effect of a knockdown on a gene. Where a factor was targeted by more than one guide set, the row with the strongest knockdown of its own transcript was retained. A knockdown was considered testable only if it achieved more than 20% knockdown of its own transcript (residual expression < 0.80) and that transcript was detectably expressed in control cells (≥ 0.05).

**Peak sets and partner-motif partition.** For every K562 factor we used the complete ENCODE4 narrowPeak set with no peak-number cap. For each dependency, the target’s peaks were scanned with the partner motif (JASPAR PWM) using FIMO at P < 1 × 10^-4^ within ±150 bp of each peak summit. A peak was called motif-positive if it contained at least one hit.

Dependencies with fewer than 20 motif-positive peaks were excluded (median 2,239 motif-positive peaks per dependency).

**Gene assignment.** Genes were assigned to a peak partition when their promoter window (transcription start site −2,000 to +500 bp, strand-aware) overlapped any peak of that partition. Genes whose promoters overlapped both partitions were removed from both, so that the motif-positive gene set is not contaminated by promoters also contacted by motif-negative peaks of the same target. From the motif-positive set we kept the genes the target’s own knockdown moved, |z_target| > 1.96; dependencies yielding fewer than 10 such genes were excluded.

These are referred to as target-moved genes (median 25 per dependency).

**Concordance.** For any factor with a testable knockdown, concordance with the target was defined as the percentage of target-moved genes for which sign(z_TF) = sign(z_target) — the fraction of the target’s own responsive genes that a second knockdown moves in the same direction. Because this is a direction-agreement statistic over independently perturbed genes, an unrelated factor is expected near 50%. Computed over a median of 25 genes, concordance is correspondingly noisy for any single dependency; inference therefore rests on the paired comparison across dependencies rather than on individual values.

**Occupancy-matched control factors.** The comparison set for each reader was drawn from factors that (i) are annotated TFs,^7^ (ii) are present in the CRISPRi library, (iii) pass the knockdown QC above and (iv) have their own K562 ENCODE ChIP-seq. For each such factor we computed its occupancy of the dependency’s motif-positive peaks, defined as the fraction of those peaks overlapped by at least one of its own peaks. The 10 factors whose occupancy was closest to the reader’s were retained as controls; the reader itself, the target, and any other candidate reader nominated for that dependency were excluded from the pool. This matching is what makes the test stringent — controls bind the same loci to a comparable extent (reader 47.6%, controls 42.0%, median matching error 3.6 percentage points), so a control cannot lose merely by being absent from the region. Each dependency contributes one point, plotting the reader’s concordance against the median concordance of its 10 controls.

**Dependency selection and statistics.** Dependencies were taken from the ARES atlas restricted to K562, excluding UNRESOLVED calls and dependencies in which the target was itself nominated as the reader, and requiring both target and reader to be testable. Where ARES nominated more than one candidate reader (16 of 50 dependencies), the candidate with the highest excess concordance was retained; because this selection uses the tested statistic, all analyses were repeated on the 34 dependencies with a single nominated reader, where no selection applies. The final set comprised 50 dependencies spanning 19 distinct readers.

Reader and control concordance were compared by a one-sided Wilcoxon signed-rank test paired within dependency, the direction (reader > control) having been specified in advance: +5.6 percentage points across all 50 dependencies (P = 3.6 × 10^-4^) and +4.3 percentage points across the 34 single-candidate dependencies (P = 3.4 × 10^-3^). Because dependencies are not independent, significance was additionally assessed by a cluster bootstrap resampling readers with replacement: +5.6 percentage points (95% CI +1.9 to +9.8, P = 0.007; 19 readers) and +4.3 percentage points (95% CI +1.5 to +8.8, P = 0.018; 15 readers), respectively.

### ADASTRA overlap analysis

Allele-specific binding variants (bQTLs) were obtained from the ADASTRA database,^68^ defined as variants associated with FDR-significant differential transcription-factor binding between alleles. ADASTRA data were not used during ARES report generation or route assignment.

For each atlas dependency, partner-motif instances were identified by a PWM log-odds scan (JASPAR motifs, both strands) in a ±15-bp window around each allele-specific-binding variant, calling a strong match when the log-odds score reached at least 0.80 of the motif’s maximal achievable score. A bQTL was assigned to a partner motif when a strong motif instance covered the variant position. A dependency was considered covered when its target factor had ADASTRA allele-specific-binding data in the relevant cell line, yielding 288 dependencies across the eight ADASTRA-covered cell lines (K562, HepG2, HEK293, MCF-7, GM12878, A549, SK-N-SH, HeLa-S3); 275 of these had a resolved route. We quantified the dependencies with at least one partner-motif bQTL.

### On-motif bQTL frequency analysis

We tested whether the on-motif rate of allele-specific-binding variants decreased across resolved routes. For each covered dependency i, we defined *y*_/_ = 1 if at least one allele-specific-binding variant fell within the partner motif and *y*_/_ = 0 otherwise. Dependencies assigned to UNRESOLVED or ARTIFACT were excluded.

To test for an ordered route trend, route was coded as SEQUENCE = 0, PROTEIN = 1 and CONTEXT = 2, and we fitted

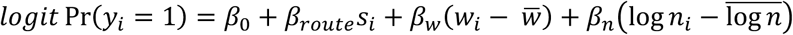

where *w*_/_ is motif width and *n*_/_ is the number of allele-specific-binding variants available for that dependency. The covariates account for differences in mutational opportunity and coverage.

The ordered trend was assessed from *β_route_*.

To estimate route-specific effects without assuming linearity, we refitted the model with SEQUENCE as the reference category and indicator terms for PROTEIN and CONTEXT. Odds ratios and Wald 95% confidence intervals were calculated by exponentiating the corresponding coefficients and coefficient intervals. As an unadjusted sensitivity analysis, we also performed a Cochran–Armitage trend test using route scores 0, 1 and 2.

### Motif-disruption dose–response analysis

For each partner-motif bQTL, the effect on motif strength was quantified as the difference between reference- and alternate-allele PWM log-odds scores, and the corresponding signed allele-specific binding effect was obtained from ADASTRA. Spearman rank correlations between motif-score change and measured allele-specific target binding were calculated separately for SEQUENCE-, PROTEIN- and CONTEXT-route dependencies. For visualization, variants were grouped into bins by motif-score change and average binding effects were plotted; all statistical tests for individual routes used the unbinned data.

### Mechanistic chain reconstruction

Before reconstructing mechanistic chains, we evaluated whether AlphaGenome reproduced the measured bQTL effects. For each partner-motif bQTL, AlphaGenome predicted the reference-and alternate-allele effects on target TF binding, which were compared with the measured ADASTRA allele-specific binding effects by Spearman correlation and sign concordance (Spearman r = 0.64, 86% of 273 variants sign-concordant; Supplementary Fig. 6a).

Chains are defined per dependency rather than per variant, so each dependency was represented by a single variant for this step. Where a dependency contained more than one on-motif bQTL (46 of 73), we took the variant with the strongest evidence of allelic imbalance.

ADASTRA reports, for each variant and target factor, two one-sided tests of the observed allelic read counts against the expectation set by the local background allelic dosage (one for preferential binding to the reference allele and one for the alternative; BH corrected). We ranked variants by –log_10_[min(FDR_ref_, FDR_alt_], that is, by the significance of whichever direction was supported, and took the maximum within each dependency; no dependency had a tie. The selection criterion was independent of both the variant’s effect on motif score and the direction of the allelic effect.

Of the 73 on-motif dependencies, 67 had a target factor with AlphaGenome ChIP coverage in the matching cell line. 60 of these 67 (90%) were direction-concordant at their representative variant and were retained as calibrated; only calibrated dependencies were carried to chain reconstruction.

Of the 60, the 25 SEQUENCE-route dependencies were set aside, as direct sequence recognition involves no trans-acting protein chain to test, and 3 unresolved dependencies were excluded. Among the remaining 32 PROTEIN- or CONTEXT-route dependencies, 22 were those for which AlphaGenome covered the chain components named in the ARES report (the target plus at least one reader or cofactor; Supplementary Fig. 6b). We classified these 22 by whether the report nominated a directional protein chain: eight did (a reader, bridge or heterodimeric partner whose allele response could be traced), and the remaining 14 inferred co-occupancy or context without a direct protein interaction and therefore no testable directional chain (Supplementary Table S5).

For the eight protein-chain candidates, we tested whether the predicted allele effect propagated through the ARES-named intermediate. For each, we computed a concordance ratio as the absolute predicted allele effect on the weakest (limiting) upstream node divided by that on the target; a large ratio indicates propagation through the chain comparable with the effect on the target. Because both quantities derive from the same predictive model, this is an in silico consistency check on ARES-nominated chains rather than an independent validation.

### Statistical analysis

All statistical analyses were performed in Python 3.11.5 using SciPy 1.16.1, statsmodels 0.14.6, NumPy 2.4.4 and pandas 2.2.2. Unless otherwise stated, tests were two-sided. Comparisons of an inferred reader against matched control factors test a directional prediction, that the reader exceeds matched controls, and were evaluated one-sided with the direction fixed before analysis. Mann–Whitney U tests were used for comparisons between independent groups and Wilcoxon signed-rank tests for paired comparisons. Fisher’s exact test was used for contingency-table analyses. Spearman correlation was used to assess monotonic relationships between continuous variables. Logistic regression was used for binary outcomes, including the bQTL enrichment analysis. Permutation tests were used for pairwise annotation-sharing analyses and cross-cell reproducibility analyses. Multiple-testing correction was performed using the Benjamini–Hochberg procedure where indicated. Exact sample sizes are reported in the Results and figure legends.

## Data availability

Transcription-factor and histone ChIP-seq, DNase-seq, RNA-seq, WGBS and in situ Hi-C datasets were obtained from ENCODE; chromatin-state segmentations are the Roadmap Epigenomics 18-state core model, downloaded as the hg38-lifted release distributed by the Roadmap Epigenomics Consortium.^33,73^ Assay availability differs by cell line, and every file used is listed with its ENCODE file and experiment accession in Supplementary Table S8.

Conservation scores were obtained from the UCSC hg38 100-way phyloP track.^47^ Protein–protein interactions were obtained from STRING v12.0.^48^ Motif position-weight matrices were obtained from JASPAR 2022 CORE vertebrate non-redundant v2, HOCOMOCO v12 CORE and CIS-BP build 2.00.^9,34,75^ PARM MPRA data were obtained from GEO accession GSE301246.^53^ Allele-specific binding variants were obtained from ADASTRA (BillCipher release, July 2022; cell-type × transcription-factor aggregation, 5% FDR; https://adastra.autosome.org).^68^ CAP-SELEX cooperativity calls were obtained from Jolma et al.^17^ Genome-wide CRISPRi Perturb-seq data in K562 were obtained from Replogle et al.^67^ Reader-perturbation ChIP-seq datasets were obtained from ENCODE and from GEO accessions GSE211293, GSE146827, GSE122847 and GSE126004;^61,62,71,72^ individual sample accessions are given in Methods.

Processed data to reproduce figures are deposited at <u>10.5281/zenodo.21806027</u>. The ARES atlas generated in this study, including all dependency-level reports, mechanistic narratives and route assignments, is available through an interactive browser at https://wanglab001-ares-atlas.static.hf.space/.

## Code availability

The ARES framework and all analysis scripts used in this study are available at https://github.com/Wang-lab-UCSD/ARES.

## Author contributions

H.X. and W.W. conceived the study and designed the research. H.X. and J.L. implemented the ARES framework. H.X. performed the computational analyses, with J.L. contributing to the MPRA analysis. H.X. curated the data and generated the figures. H.X. and W.W. wrote the manuscript. W.W. supervised the study and acquired funding. All authors read and approved the final manuscript.

