## Supplementary Data for "An atlas of transcription factor cooperation reveals how motif readers shape regulatory output"

### Supplementary Methods

#### Sequence encoding and motif scanning

For each retained target peak, the 200-bp sequence (midpoint-centered, as described in Methods) was one-hot encoded into a 200 × 4 matrix, with the four columns corresponding to A, C, G and T; positions containing an ambiguous base (N) were encoded as all-zero. Each JASPAR position-weight matrix was applied as a one-dimensional convolutional filter across the encoded sequence, producing a match score at every window position. For each motif and peak, the score was taken as the maximum over all window positions and over both the forward strand and the reverse complement (the best-hit score), yielding a peaks × motifs score matrix. These best-hit scores were used both for partner-motif discovery (as random-forest features) and as the partner-motif strength term (M) in the partner-dependence-coefficient model.

#### MPRA bidirectionality analysis for signed-non-significant dependencies

To test whether dependencies that were non-significant in the signed MPRA analysis nevertheless contained promoter subsets with strong motif-associated activation and repression, we re-analyzed the PARM promoter MPRA dataset (GEO GSE301246) using the same preprocessing as in the primary MPRA analysis.[1]

For each target–partner dependency, reporter fragments were first restricted to those overlapping the target-transcription-factor ENCODE ChIP-seq peaks, as in the primary MPRA analysis. Retained fragments were scanned for the partner motif using FIMO from MEME Suite v5.5.5[2] with the JASPAR 2022 CORE[3] vertebrates non-redundant v2 motif collection at $P < 1 \times 10^{-4}$. When the motif-positive rate exceeded 0.5, the threshold was tightened to $P < 1 \times 10^{-5}$. Fragments containing at least one partner-motif hit were labelled motif-positive, whereas fragments without a hit were labelled motif-negative.

To reduce confounding by basic sequence and positional features, each motif-positive fragment was matched to a motif-negative fragment by nearest-neighbor matching in standardized covariate space, using fragment length, GC content and distance to the transcription start site. Covariates were z-scored on the pooled motif-positive and motif-negative fragment distribution, and greedy cKDTree matching was performed without replacement using a fixed random seed.

Matched fragments were then grouped by parent promoter. For each promoter, the motif-associated activity difference was computed as the median activity of motif-positive fragments minus the median activity of their matched motif-negative fragments. This difference was standardized by the within-pair activity standard deviation, yielding Δ activity in SD units. Standardization was performed within each dependency so that dependencies with larger activity variance did not dominate the pooled strong-tail estimates. Promoters with Δ activity ≥ +1 SD were assigned to the strong-activation tail, whereas promoters with Δ activity ≤ −1 SD were assigned to the strong-repression tail.

For each dependency, observed strong-tail fractions were compared with a selection-matched shuffled-label null. Motif-positive and motif-negative labels were permuted, and the identical matching, promoter grouping and tail-fraction calculation were repeated to estimate the fraction expected by chance under the same selection procedure. Thus, the null controlled for the matching and selection steps used to define motif-positive and matched motif-negative fragments. Observed activation- and repression-tail fractions were each tested against their corresponding shuffled-label null using a one-sided paired Wilcoxon signed-rank test across dependencies, with dependency as the unit of analysis.

This analysis was restricted to 631 dependencies that were non-significant in the primary signed MPRA test (paired-locus BH q ≥ 0.05), were present in the canonical ARES atlas, and were assayed in one of the four PARM cell lines represented in the atlas: K562, HepG2, HEK293 and MCF-7. A dependency was classified as bidirectional when both its strong-activation and strong-repression fractions exceeded their corresponding per-dependency shuffled-label null fractions. This criterion requires only that each fraction exceed its null, without a per-dependency significance threshold; under independence of the two comparisons, approximately 25% of dependencies would satisfy it by chance. By this criterion, 438 of 631 signed-non-significant dependencies, or 69%, showed bidirectional motif-associated MPRA activity, against the approximately 25% expected under the chance baseline. This chance expectation is intended only as a heuristic baseline because activation- and repression-tail enrichments are not strictly independent.

**Baseline models for TF-binding prediction**

To assess whether motif-based feature representations are competitive with sequence-based deep learning for predicting transcription-factor occupancy, we compared RF-JASPAR, the random-forest model used for partner-motif discovery in ARES, against a linear motif baseline (Ridge-JASPAR) and three sequence-based neural networks (DeepCNN, DanQ[4] and ExplaiNN[5]) (Supplementary Fig. 8a).

RF-JASPAR is the random-forest regressor described above (Partner-motif discovery): for each target ChIP-seq dataset it predicts the log1p-transformed ChIP signal of a peak from the 841-dimensional vector of JASPAR 2022 motif scores for that peak. Ridge-JASPAR is a ridge regression on the same 841 motif features, included to isolate the contribution of the motif representation from that of the nonlinear model. The three neural networks take the one-hot-encoded 200-bp peak sequence as input, the same window from which the motif scores were computed: DeepCNN is a three-layer convolutional network, with each convolutional layer followed by batch normalization, ReLU activation and max-pooling; DanQ is a hybrid convolutional-recurrent network that combines a convolutional layer for local sequence features with a bidirectional LSTM layer for longer-range dependencies, run with the default parameters of the original implementation; and ExplaiNN is an interpretable convolutional network, run with 300 convolutional units and otherwise default parameters.

For every (cell line, target TF) ChIP-seq dataset, peaks were partitioned 70:10:20 into training, test and validation sets by a seeded shuffle (seed 1337). The same partition was used for all five models, so that every model was trained on the same peaks and scored on the same held-out test peaks. The validation partition was used for early stopping in the neural networks; RF-JASPAR and Ridge-JASPAR did not use it and were trained on the 70% training partition alone. Performance was quantified as the Pearson correlation between predicted and observed signal on the 10% test partition. RF-JASPAR and Ridge-JASPAR returned a result for all 1,552 atlas datasets; DanQ and DeepCNN for 1,551 and 1,550 respectively, the missing datasets reflecting occasional training instability; and ExplaiNN for 1,507, failing to train on the 45 datasets with the fewest peaks (all with 623 or fewer target peaks; the smallest dataset on which ExplaiNN trained successfully had 639). To keep the comparison paired, all five models were evaluated on the 1,504 atlas datasets scored by every model, and models were compared by paired two-sided Wilcoxon signed-rank tests across these datasets.

RF-JASPAR was used for partner-motif discovery despite DanQ's slightly higher accuracy because the analysis requires attributing occupancy to named motifs. With motif scores as features, the TreeSHAP importance of each feature is by construction the contribution of a specific JASPAR motif, scanned at its own length. A convolutional network trained on raw sequence learns filters that do not correspond to named motifs, and recovering which motif a filter represents requires a post hoc matching step with its own thresholds; a filter may also encode a blend of motifs or a pattern matching none. The convolutional layer could in principle be initialized with JASPAR PWMs so that each filter is a named motif, but filters within a layer share a fixed width whereas JASPAR motifs vary in length, so attribution would be to padded or truncated versions of each motif rather than to the motif as defined. ExplaiNN, which was designed to make filters interpretable, performed worst on this task and failed to train on the smallest datasets, so its interpretability was not available at acceptable accuracy. Notably, the gain from allowing nonlinear combinations of motif scores (Ridge-JASPAR to RF-JASPAR, $\Delta r$ = 0.07) exceeded the further gain from learning features directly from sequence (RF-JASPAR to DanQ, $\Delta r$ = 0.02), consistent with the motif-score representation capturing most of the sequence information relevant to predicting occupancy within a peak window. We therefore accepted a median difference of $\Delta r$ = 0.02 in exchange for feature importances that are, without further inference, the identities of motifs.

**Background-PC residualization of reader-target correlations**

To test whether reader-target tracking at namesake-unbound motif loci reflected generic ChIP occupancy rather than reader-specific signal, we repeated the reader-target correlation analysis after removing principal components of background TF signal. For each hidden-reader dependency, we used the same namesake-unbound partner-motif loci defined for the main reader-tracking analysis. Dependencies with more than 400 tested loci were randomly subsampled to 400 loci using a fixed seed.

At each locus, ChIP-seq signal was quantified for the target, the inferred reader, ten same-cell random null TFs and twenty additional same-cell background TFs. The null and background TF sets were sampled without overlap and excluded the target, motif namesake and inferred reader. All signal vectors were rank-transformed and z-standardized. Principal components were computed from the 20-background-TF signal matrix. Target, reader and null-TF signals were then residualized against an intercept plus the first k background PCs, where $k$ = 0, 1, 3 or 5. Correlations after residualization were computed between the residualized target signal and either the residualized reader signal or each residualized null TF signal. The median of the ten null-TF correlations was used as the matched null for that dependency.

Reader correlations were compared with matched null correlations across dependencies using a one-sided paired Wilcoxon signed-rank test. Per-dependency significance shown in the figure was computed from the residualized reader-target correlation using a two-sided t approximation with degrees of freedom $n - 2 - k$, followed by Benjamini–Hochberg correction across dependencies within each panel. Marker area indicates the fraction of tested loci that overlapped an inferred-reader ChIP-seq peak.

### Supplementary Figures

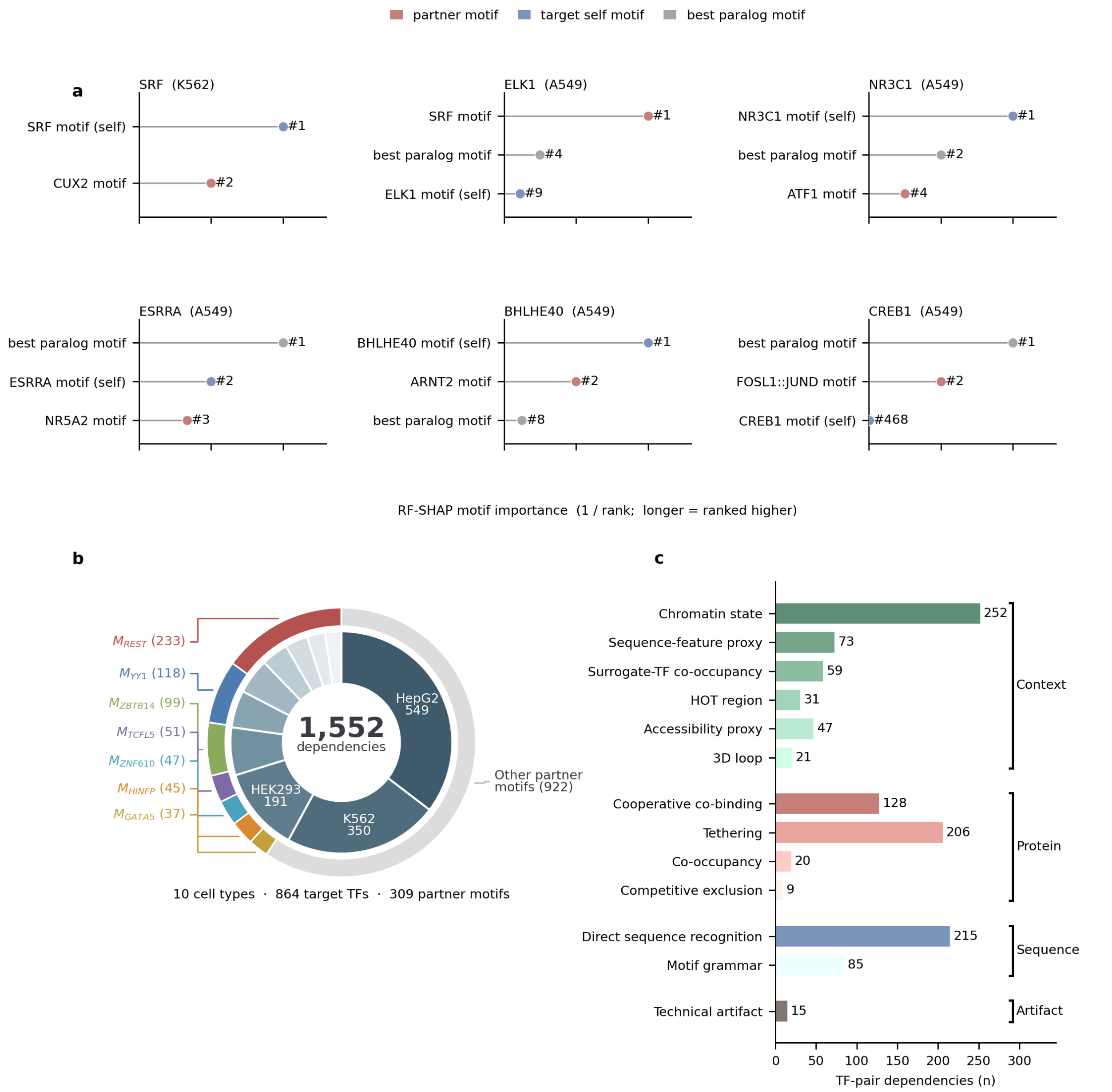


**Supplementary Fig. 1 | Motif-ranking examples and composition of the ARES dependency atlas.**

**a,** Representative random-forest SHAP motif-ranking examples for target TF ChIP-seq datasets. For each target dataset, motif scores were used to predict quantitative target occupancy, and motifs were ranked by RF-SHAP importance. The target self motif, best paralog motif and selected partner motif are highlighted. The x-axis shows inverse motif rank, so longer horizontal positions indicate higher-ranked motifs. These examples illustrate cases in which the selected partner motif ranked above, below or between the target self/paralog motifs, including cases where the target self motif was not among the top predictors.

**b,** Summary of the final ARES atlas. The inner ring shows the number of dependencies contributed by major cell types, and the outer ring highlights recurrent partner motifs, with remaining partner motifs grouped as “Other partner motifs”. The atlas contains 1,552 TF-motif-cell dependencies across 10 cell types, involving 864 target regulatory proteins and 309 partner motifs.
**c,** Distribution of curated ARES mechanism leaves across the final atlas. Mechanism leaves are grouped into Context, Protein, Sequence and Artifact classes. Context mechanisms were dominated by chromatin-state and sequence-feature-proxy explanations, Protein mechanisms by tethering and cooperative co-binding, and Sequence mechanisms by direct sequence recognition and motif grammar. Bar labels indicate the number of dependencies assigned to each mechanism leaf.

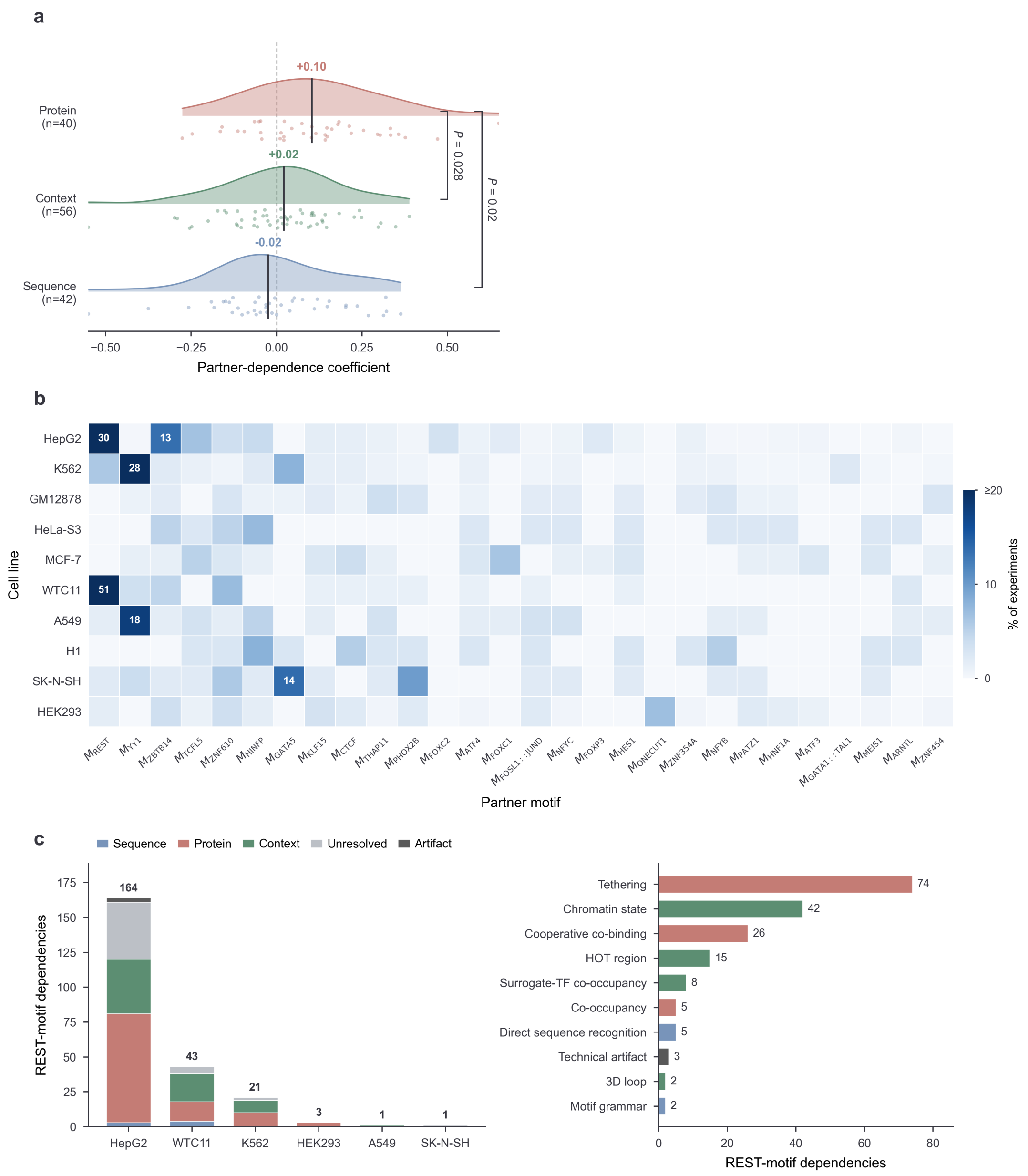


**Supplementary Fig. 2 | Route robustness and recurrent-motif structure.**
**a**, Partner-dependence coefficient by route after collapsing each partner motif to a single median within each route, controlling for hub-motif recurrence. Points are route × partner-motif combinations (42 SEQUENCE, 56 CONTEXT, 40 PROTEIN; 138 total), drawn from 94 distinct partner motifs, 33 of which are assigned to more than one route across cells or targets and therefore contribute a point to each. Vertical lines and colored values indicate medians. PROTEIN-route motifs retained significantly higher coefficients than both CONTEXT ($P = 0.028$) and SEQUENCE ($P = 0.02$; two-sided Mann–Whitney U), supporting that the elevated protein-route dependence in Fig. 2a is not driven by recurrent hub motifs. SEQUENCE and CONTEXT overlapped near zero, consistent with the coefficient measuring dependence on motif-namesake-TF occupancy rather than distinguishing these two protein-independent routes.
**b**, Recurrence of partner motifs across cell lines. Color and labelled value both give the percentage of a cell line's atlas dependencies carrying the indicated partner motif; cells ≥ 10%

are labelled and color saturates at ≥ 20%. A few motifs, notably REST motif and YY1 motif, dominate within specific lines, illustrating the hub structure that motivates the per-motif de-duplication used in **a** (Methods).

**c**, The REST motif as the largest recurrent hub. Left, dependencies with REST as partner motif per cell line, stacked by route (including unresolved and artifact); these concentrate in HepG2 and are predominantly PROTEIN or CONTEXT. Right, the ten most frequent resolved mechanism leaves among REST-motif dependencies, pooled across cell lines (182 dependencies; unresolved excluded), colored by parent route. Tethering dominates, consistent with REST recruiting proteins to RE1-bound loci through corepressor complexes rather than direct sequence recognition by those proteins.

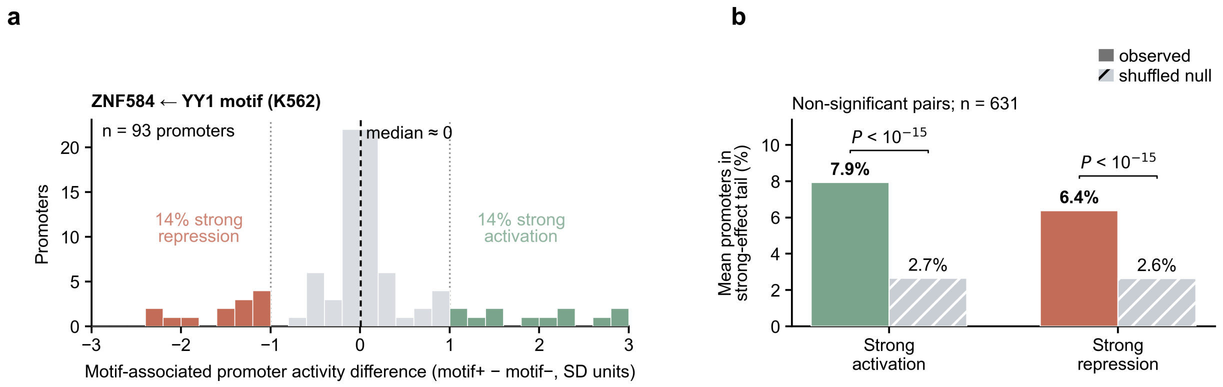


**Supplementary Fig. 3 | Direction-heterogeneous MPRA effects cause signed tests to undercount partner-motif cis-function.**

**a,** Representative bidirectional dependency, ZNF584 ← YY1 motif in K562. For each promoter carrying the YY1 partner motif, motif-associated reporter activity was defined as the median activity of motif-positive fragments minus that of covariate-matched motif-negative fragments, expressed in within-pair standard-deviation units ($motif^{+} - motif^{-}$; n = 93 promoters). Although the distribution is centered near zero and the signed paired test is non-significant, 14% of promoters fall in the strong-repression tail (≤ −1 SD) and 14% in the strong-activation tail (≥ +1 SD), indicating direction-heterogeneous cis-regulatory effects that cancel under signing.

**b,** Population analysis of 631 signed-non-significant dependencies from the primary MPRA test. Bars show the mean fraction of promoters in each strong-effect tail, observed versus a selection-matched shuffled-label null. Both tails are enriched over null: strong activation, 7.9% versus 2.7%; strong repression, 6.4% versus 2.6% (one-sided paired Wilcoxon signed-rank test across dependencies, $P < 10^{-15}$ for both). A dependency was classified as bidirectional when both strong-tail fractions exceeded their corresponding per-dependency null fractions; 438 of 631 dependencies met this criterion, against approximately 25% expected if the two tail comparisons were independent. Thus, motif-level cis-function is widespread even among dependencies called null by a signed test because promoter-specific activation and repression offset one another.

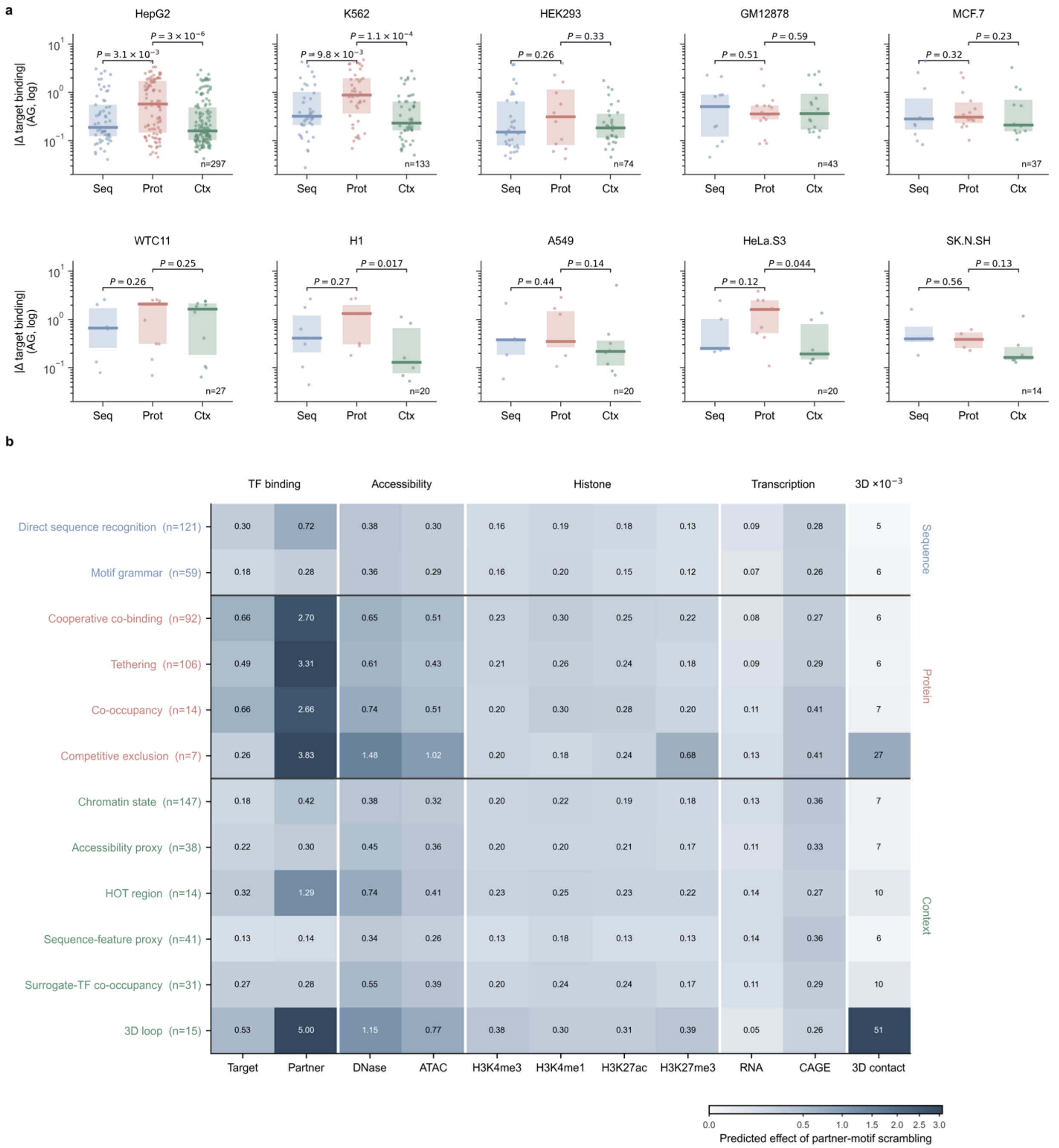


**Supplementary Fig. 4 | AlphaGenome partner-motif scrambling reveals route- and mechanism-specific regulatory signatures.**

**a,** Per-cell-line absolute predicted change in AlphaGenome target-binding output after partner-motif scrambling, where the change is defined as in Fig. 3d (Methods). Dependencies are grouped by ARES route. Points are dependencies; boxes show medians and interquartile ranges. P values are two-sided Wilcoxon rank-sum tests comparing PROTEIN-route dependencies with SEQUENCE and with CONTEXT dependencies separately; n is the number of scored dependencies per cell type. Absolute effect size alone does not assign a dependency to a route, but PROTEIN-route dependencies show larger predicted target-binding changes than the other routes in most cell lines.

**b,** Predicted response to partner-motif disruption across AlphaGenome regulatory outputs, stratified by ARES mechanism class. The plotted value is the median absolute Δ across a class's dependencies, where Δ is defined as in Fig. 3d (Methods). Rows are mechanism leaves grouped by ARES route, with the number of scored dependencies in parentheses. Columns group AlphaGenome outputs into transcription-factor binding, chromatin accessibility, histone modifications, transcription and three-dimensional contact. Color encodes effect size on a square-root scale; three-dimensional contact is summarized as the mean and displayed as $\times 10^{-3}$ owing to its distinct dynamic range. Perturbation fingerprints are mechanism-specific rather than strictly route-specific: PROTEIN-route mechanisms generally show prominent partner-binding responses and SEQUENCE mechanisms weaker partner dependence, whereas CONTEXT mechanisms are heterogeneous, with HOT-region and 3D-loop classes showing elevated accessibility effects and the 3D-loop class the strongest predicted change in contact.

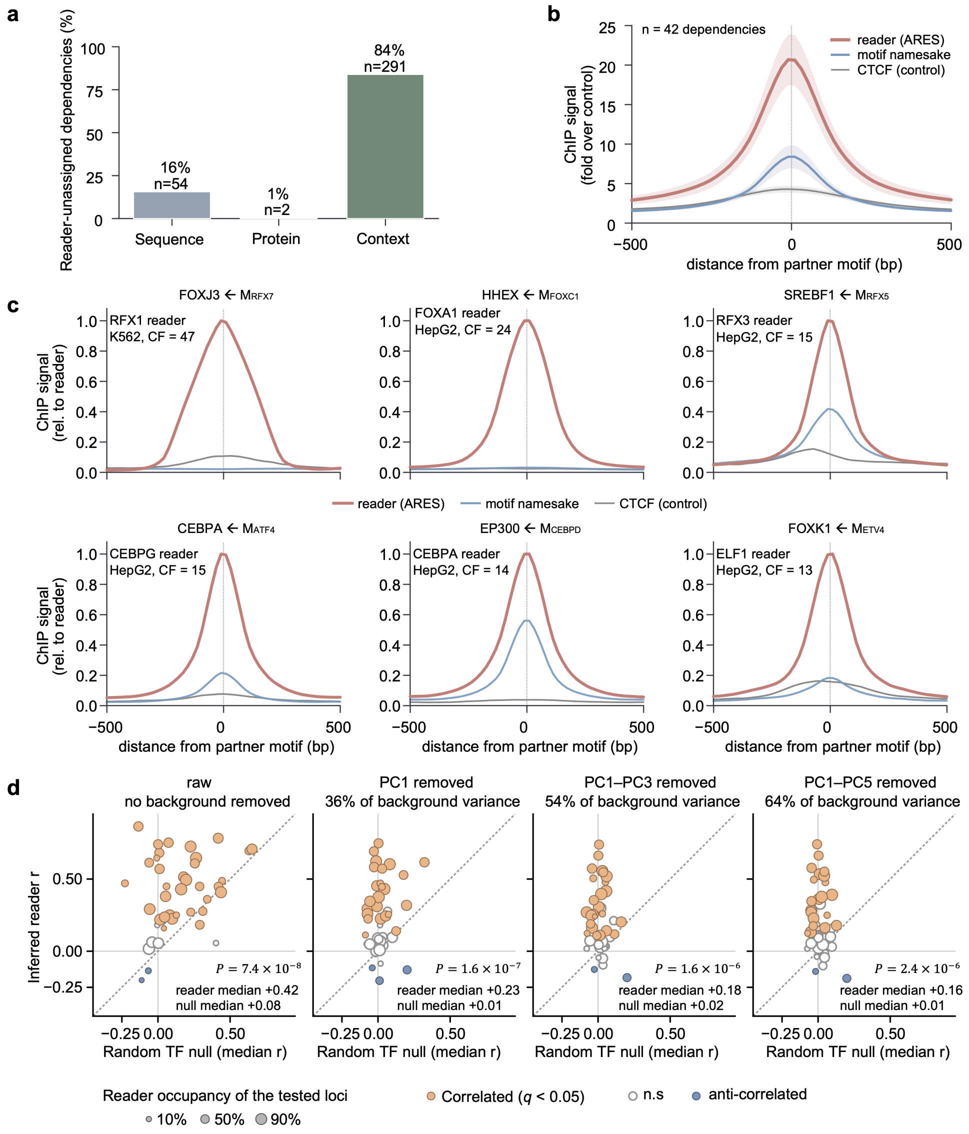


**Supplementary Fig. 5 | ARES-inferred readers, rather than motif namesake factors, support protein-route regulatory mechanisms.**

**a**, Cooperation-route composition of the resolved dependencies for which ARES assigned no reader (n = 347). Most are context-route (84%), in which the partner motif marks chromatin state, accessibility or higher-order architecture rather than recruiting a motif-reading protein, so the absence of a reader is an expected property of the mechanism. The sequence-route cases (16%) are predominantly composite or overlapping motif architectures, for which no single reader protein is named. Two protein-route dependencies remain reader-unassigned, both MNT/NPAS2 motif (K562 and MCF-7), in which the partner motif is a variant E-box engaged by the target through an obligate dimerization partner; these are boundary cases between the sequence- and protein-routes.

**b**, Aggregate ChIP-seq signal (fold enrichment over control) centered on the partner motif for the inferred reader, motif namesake factor, and CTCF (negative control) across 42 hidden-reader dependencies with expressed namesake (TPM > 0.5) and available ChIP-seq data.

Mean signal peaked at 20.7-fold for the inferred reader, versus 8.4-fold for the motif namesake and 4.3-fold for CTCF at the partner motif.

**c**, Aggregate ChIP-seq signal centered on the partner motif for six representative dependencies in HepG2 and K562. Profiles show the ARES-inferred reader, the motif namesake factor and a CTCF control, each normalized to the reader peak. Panel titles indicate the target and partner motif; annotations indicate the inferred reader, cell type and centering factor (CF), defined as the center-to-flank ratio of the reader signal around the motif. In each example, the inferred reader shows the strongest motif-centered signal. Examples are the six highest-CF dependencies with CF ≥ 10, selecting one example per partner motif for which namesake-factor and CTCF ChIP-seq were available.

**d.** Background-PC residualization of reader-target tracking. Reader-target correlations at namesake-unbound partner-motif loci were recomputed after removing principal components of generic same-cell TF occupancy. Each point is one hidden-reader dependency from the PCA-compatible reanalysis; loci were capped at 400 per dependency. x, median target correlation for ten random same-cell TFs; y, inferred-reader correlation, after residualizing target, reader and null-TF signals on 0, 1, 3 or 5 PCs from 20 disjoint background TFs. Panel titles show the median fraction of background-TF variance removed. Fill indicates reader-target significance after residualization (BH $q < 0.05$); marker area indicates the fraction of tested loci inside a reader ChIP-seq peak. P values are one-sided paired Wilcoxon signed-rank tests comparing reader correlations with matched null medians across dependencies.

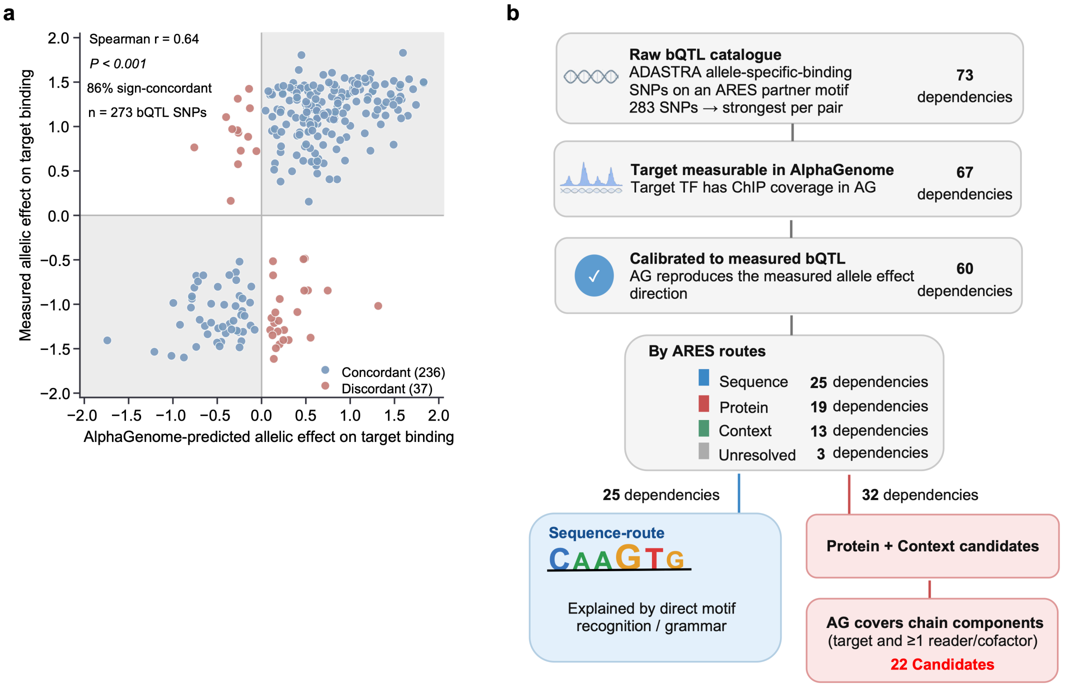


**Supplementary Fig. 6 | AlphaGenome calibration and filtering strategy for partner-motif bQTL analysis.**

**a,** Concordance between AlphaGenome-predicted and measured allelic effects on target-factor binding for partner-motif bQTL SNPs. Each point represents one bQTL SNP. The x axis shows the AlphaGenome-predicted allelic effect on target binding, and the y axis shows the measured bQTL allelic effect. Effects are shown as signed transformed allelic ratios, with positive values indicating stronger binding on the reference allele and negative values indicating stronger binding on the alternate allele. Shaded quadrants indicate sign concordance between prediction and measurement. AlphaGenome predictions were directionally concordant with measured effects for 236 of 273 SNPs (86%; Spearman r = 0.64, *P* < 0.001).

**b,** Filtering workflow used to define bQTL dependencies suitable for mechanism tracing. Starting from ADASTRA allele-specific binding SNPs overlapping ARES partner motifs, one representative SNP was retained per target–partner dependency, yielding 73 dependencies. Of these, 67 had target-factor ChIP coverage in AlphaGenome. AlphaGenome reproduced the measured bQTL effect direction for 60 dependencies, which were retained for downstream analysis. These calibrated bQTLs comprised 25 sequence-route, 19 protein-route, 13 context-route and 3 unresolved dependencies. Sequence-route bQTLs were interpreted as direct motif-recognition or grammar effects. Protein- and context-route candidates were further restricted to cases in which AlphaGenome covered the target and at least one inferred reader or cofactor, yielding 22 candidate chains for mechanistic reconstruction.

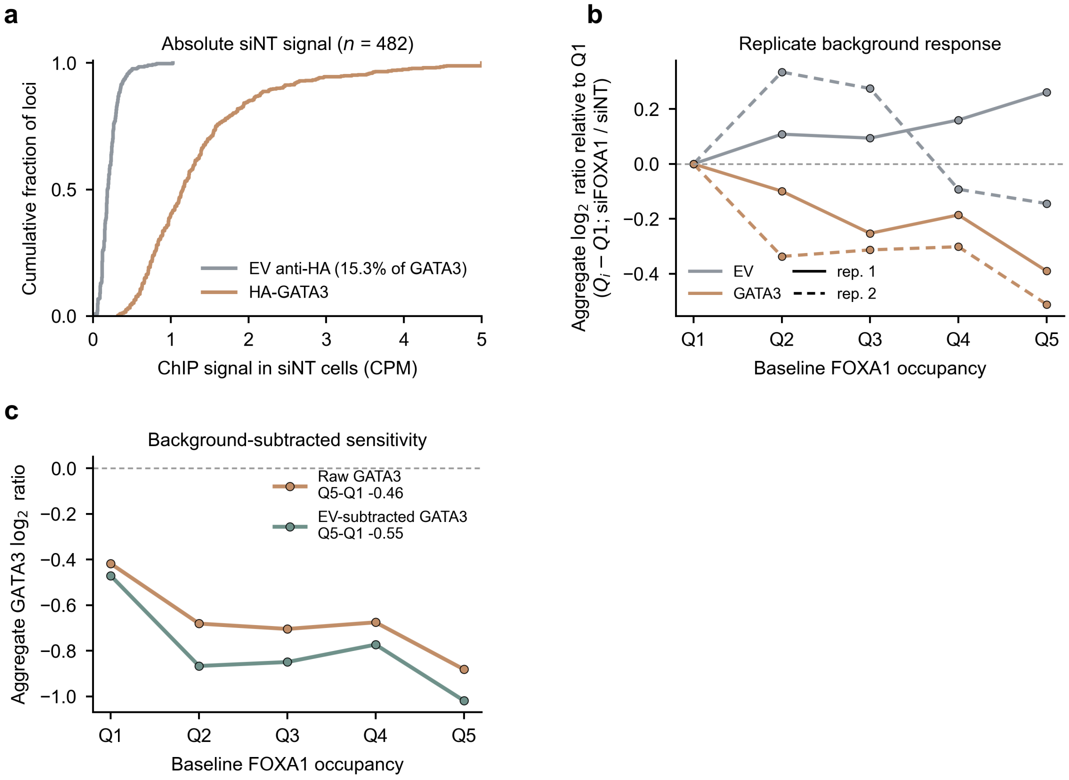


**Supplementary Fig. 7 | Controls for the FOXA1–GATA3 analyses.**
**a,** Cumulative distribution of ChIP signal in siNT cells across the 482 tethered loci of Fig. 5g. Empty-vector (EV) cells profiled with the same anti-HA antibody carry measurable but low background, 15.3% of HA-GATA3 signal in aggregate.
**b,** Response to FOXA1 knockdown by quintile of baseline FOXA1 occupancy, shown relative to Q1 so that gradients can be compared directly; replicates are plotted separately and quantification is as in Methods. Both HA-GATA3 replicates decline in the same direction (Q5 − Q1 = −0.39 and −0.51), whereas the two empty-vector replicates disagree in sign (+0.26 and −0.14) and their aggregate shows no gradient (+0.005). The overall anti-HA background level also changes after knockdown, but not reproducibly (aggregate −8.7%; the replicates disagree in direction). Q1 referencing removes the uniform component of any such change, so only a reproducible quintile-dependent background shift could produce the GATA3 gradient. Aggregate ratios exceed the per-locus medians reported in Fig. 5g (Q5 − Q1 = −0.46 versus −0.20 across the same 482 loci) because pooling weights loci and replicates by signal (Methods); the per-locus median is the more conservative summary used in the main figure.
**c,** Subtracting the quintile-matched empty-vector background from HA-GATA3 strengthens rather than weakens the gradient (Q5 − Q1 = −0.55 versus −0.46), confirming the gradient is not an artifact.

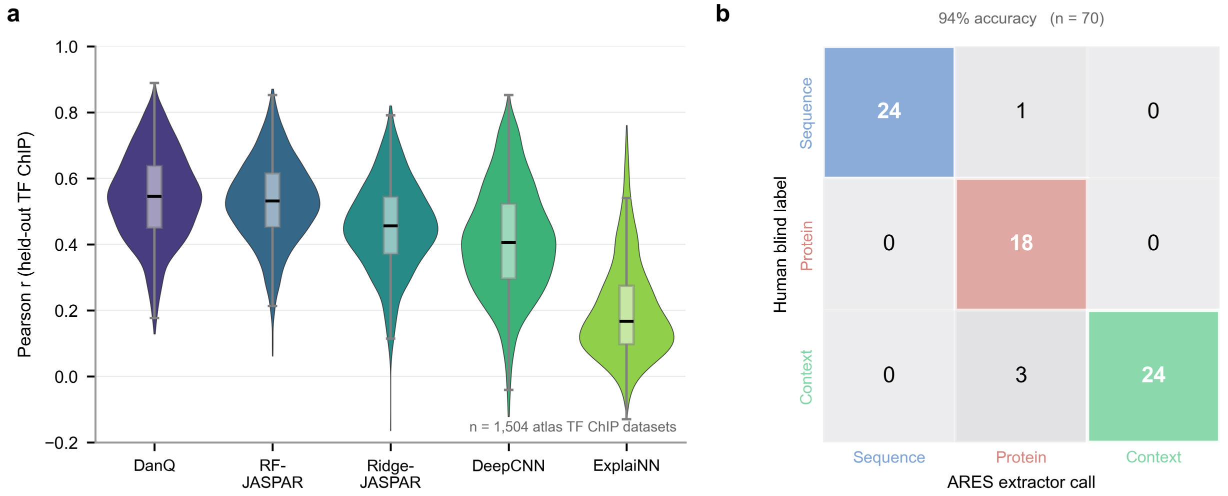


**Supplementary Fig. 8 | Motif-based random forests approach deep-learning accuracy for TF-binding prediction, and the ARES extractor reproduces route blind calls by a human expert.**

**(a)** Held-out prediction of transcription-factor ChIP-seq signal by five models trained on DNA sequence or motif features: RF-JASPAR (random forest on JASPAR motif scores, the model used by ARES), Ridge-JASPAR (ridge regression on the same motif features), DanQ (convolutional-recurrent network), DeepCNN and ExplaiNN. All five models were evaluated on an identical set of 1,504 TF ChIP-seq datasets, the subset of the 1,552 (cell line, target TF) datasets of the ARES atlas on which every model trained successfully (Supplementary Methods), so that the comparison is paired. Each violin summarizes the test-set Pearson correlation across datasets; boxes show the median and interquartile range, whiskers the 1.5× interquartile range. Median r was 0.55 (DanQ), 0.53 (RF-JASPAR), 0.46 (Ridge-JASPAR), 0.41 (DeepCNN) and 0.17 (ExplaiNN). DanQ exceeded RF-JASPAR by a median paired difference of $\Delta r = 0.02$, a systematic but small margin (DanQ higher in 861 of 1,504 datasets, 57%; paired two-sided Wilcoxon signed-rank $P = 1.5 \times 10^{-9}$). RF-JASPAR was used for partner-motif discovery because its feature importances map directly onto named motifs, whereas the sequence-based models achieve comparable accuracy but do not attribute predictions to named motifs without an additional inference step.

**(b)** Validation of the ARES route extractor against blind annotation by a human expert. An expert reviewed 72 dependencies while blinded to the automated call. Seventy were confidently assigned to Sequence, Protein or Context, whereas two could not be confidently classified and were excluded from the confusion matrix. Cells give the number of dependencies among the 70 with a confident human label; rows are the human blind label and columns the ARES extractor call. Diagonal agreement was 66 of 70 (94%). All four disagreements involved the extractor calling Protein: three dependencies the annotator labelled Context, and one labelled Sequence.

**Supplementary Note 1 | Deterministic safeguards in ARES.**

**Design principle.** ARES treats large-language-model outputs as proposals, not verdicts. Each agent—hypothesis generation, review, coding and summary—contributes language-model judgment, but the properties on which the atlas depends (fidelity of reported evidence to executed computation, separation of mechanism from co-occurrence, calibration of conclusions to the data, and the decision to terminate) are enforced by checks external to the generating model. These four properties were not chosen ad hoc: they correspond to the principal ways in which a language-model agent can produce a fluent but unsupported scientific report—asserting entities or numbers absent from the computation, relabeling a correlation as a mechanism, over-stating what the data establish, and stopping before the evidence warrants.[6] Rather than adding checks reactively as individual failures surfaced, we enumerated these failure modes and placed an external check on each, so that the set of safeguards is derived from the ways the inference can fail rather than assembled case by case. For each we state the strength of enforcement, distinguishing guards implemented as deterministic code from steps mandated in an agent's instructions, so that the guarantee is neither uniform nor overstated. Below we describe these checks and the strength of enforcement for each.

**Persistence as the audit foundation.** Every code block executed by the coding agent, together with its standard output, is written to disk before any downstream evaluation. The summary agent evaluates the coding result against this logged output, and the fidelity checks below operate on the log rather than on model recollection. Because a claim must trace to a log entry, a statistic or entity with no corresponding execution output has no support to point to. The coding loop executes any code block before a conclusion is accepted—running the block even when the model emits code and a solution together, and surfacing each result before the next step—so a solution cannot form without a corresponding execution log. A technical failure—including code that silently caught its own exception while the kernel reported success—is detected from the printed traceback and routed to a bounded retry rather than interpreted as a biological result. This persistence is the foundation on which the entity-fidelity check operates.

**Entity-fidelity check (deterministic).** Before a summary is accepted, a routine extracts gene-symbol-like tokens from the model's summary and findings and compares them against the concatenated standard output of the code that actually ran. Any token absent from the execution log is marked with a fabrication warning and the report's confidence is reduced. This runs on every evaluation with no model in the loop; it is the mechanism that would flag a factor named in prose but never analyzed—for example, a cohesin subunit asserted in a report whose enrichment was never computed.

**Number-verification step (required, prompt-enforced).** The coding agent is instructed to run a final verification cell before emitting a solution, re-deriving every reported number from the underlying variables and checking directionality, hardcoded constants and contingency-table orientation, and requiring that every named entity originate from a variable computed in the same block. This is a required step in the coding contract rather than a code-level gate: it enforces number-level fidelity by construction of the workflow, but is not independently verified to have executed. We therefore describe entity-level fidelity as deterministically checked and number-level fidelity as required of the coding agent. A misattribution in which a computed result is reported under the wrong factor's label—where the erroneous label is itself present in the execution log—falls to this prompt-level entity-from-variable rule rather than to the deterministic entity check. Reported magnitudes are held to the same standard: intensifiers are disallowed unless attached to a computed number, so that an un-quantified magnitude cannot propagate to the hypothesis agent, which sees only the summary and not the raw output.

**Discriminative rigor (three layers).** Separating a direct mechanistic effect from mere co-occupancy is enforced at three points. At generation, the hypothesis agent must complete a scratchpad contrasting the signal expected if the mechanism holds against the signal expected under co-occupancy alone, and must declare whether the two are distinguishable; if not, the predicted test is redesigned. At review, the screening agent re-checks discriminative value. At evaluation, a control comparison can override a mechanism call that is no stronger than a simpler baseline (below). The property the atlas depends on—that a named mechanism is not a relabeled correlation—is thus checked before, during and after testing.

**Interpretation ceiling and evidence discipline (required of the summary agent).** When a control or comparator shows that the mechanism-specific effect is no stronger than a simpler baseline, the summary agent is required to downgrade the named mechanism to INCONCLUSIVE or REJECTS, to record the simpler explanation that suffices, and to keep the report's wording within the supported interpretation. The summary agent is further required to report the outcome of every planned test, including null or inconvenient results, and to justify a SUPPORTS against each sub-prediction of a compound hypothesis; omitting a sub-result is defined as cherry-picking and disallowed, so a supported call cannot rest on a favorable subset of the evidence. These calibration steps guard against promoting a co-occurrence to a mechanism, and against reporting only the evidence that succeeded; they are enforced in the summary agent's instructions rather than as code-level gates.

**Causal-language calibration (required by the summary agent).** Conclusions are gated by data type. On observational functional-genomic data the pipeline forbids unqualified causal verbs—that a factor "drives," "causes" or "proves" an effect—and requires the report to state explicitly that causality is not established; unqualified causal phrasing is reserved for analyses that include a manipulation. Because the atlas is built from observational assays, this keeps every mechanistic report within the interpretation its data support.

**Commensurable measurement (required + reactively enforced).** Quantities that are compared across dependencies are computed through a shared library of measurement helpers with pinned parameters, so that the same operation yields comparable numbers in every run rather than reflecting per-run analytic choices. The comparability-critical helpers and the quantities they standardize include motif scanning (FIMO at $P < 1\times10^{-4}$, with masking disabled) for motif prevalence and enrichment; peak intersection with a fixed overlap denominator (overlaps / query peaks, zero flank) for co-occupancy rate; summit-centered signal extraction (250-bp window, mean statistic) for ChIP signal intensity; a fixed ±2-kb TSS window for promoter proximity and a fixed distance cutoff (50 kb) for peak-to-gene assignment; binned signal matching (fixed bin count and seed) for signal-matched controls; a STRING confidence threshold (score ≥ 400) for protein–protein interaction evidence; Tomtom (fixed threshold and distance metric) for motif similarity; and a case-insensitive, ambiguity-aware GC definition that removes a soft-masking artifact. Manifest categories are mapped to helper bundles through deterministic dispatch with no model in the routing path, and a coverage test fails if a category is unmapped. A larger set of shared routines handles data parsing and identifier reconciliation (for example, mixed gene-identifier namespaces and peak-to-sequence header matching); these fix correctness but do not carry a compared-quantity parameter and are not comparability anchors.

Enforcement is by contract and reactive rejection rather than a static guarantee: the coding contract requires these helpers in place of hand-rolled alternatives, and specific bypassing patterns are rejected on retry, but a run is not statically prevented from computing a quantity outside the library on a first attempt. Three parameters are standardized by documented rule rather than by an immutable helper default and are described as such: the motif threshold is pinned at $P < 1\times10^{-4}$ with a documented, rule-based relaxation to $P < 1\times10^{-3}$ for weak or DBD-less motifs; the expression cutoff used in the artifact check (TPM < 0.5) is a pipeline rule applied outside the shared library rather than a pinned helper parameter; and chromatin-state groupings are computed through a shared routine but with the promoter and enhancer state sets supplied per run rather than fixed. Commensurability therefore holds strongly for the pinned measurement quantities above and is approached, rather than guaranteed, for these three rule-governed parameters.

**Termination guards.** The decision to stop is removed from unilateral model control at several points:

- *Convergence veto (deterministic).* When the summary agent reports convergence, a deterministic check—evaluated in a fresh context independent of the reasoning that produced the hypotheses—verifies that at least one hypothesis (excluding the iteration-0 artifact check) reached SUPPORTS and that convergence rests on at least two genuinely independent measurement modalities; multiple analyses of the same assay do not count as independent support. Where a supported mechanism names a third-party factor that was not itself tested, convergence is refused until that factor is verified. If these conditions are unmet, convergence is reset and the run continues.
- *Guarded budget accounting (deterministic).* The iteration budget is charged only for hypotheses that pass review and data-availability checks; hypotheses rejected at review, starved of data, or failing technically are governed by separate circuit breakers and do not consume it. A run therefore ends in one of three recorded states: guarded convergence; a salvage stop that reports the supported evidence gathered when no further progress is possible; or exhaustion of the iteration limit.
- *Authority downgrade (deterministic).* The hypothesis agent may not declare convergence; any such outcome is rewritten as a new hypothesis, so that termination can occur only through the guarded summary step.
- *Outcome-table supersession (deterministic).* Where the report narrative and the recorded hypothesis outcomes conflict, a deterministic outcome table is appended that supersedes the narrative.
- *Artifact clamp (deterministic).* For the initial technical-artifact check, a claimed SUPPORTS is clamped to INCONCLUSIVE when expression data are absent from the manifest, preventing an artifact call from resting on self-motif enrichment alone.

Together, these safeguards implement the principle that model judgment is bounded by external verification. The model proposes hypotheses, writes and runs code, and interprets results, but key failure modes are constrained outside the generating model: entity provenance and termination are checked deterministically; cross-dependency measurements are routed through shared routines where possible; artifact calls are clamped by predefined rules; and causal language, mechanism calls and interpretation ceilings are governed by mandatory reporting rules. These safeguards do not make the inferred biology automatically correct, but they make each conclusion traceable to executed evidence and make unsupported reporting auditable rather than hidden in fluent model prose.

**Supplementary Note 2 | The REST motif predicts RCOR1 occupancy through CoREST tethering rather than direct recognition.**

RCOR1 is a core subunit of the CoREST corepressor complex and is not a sequence-specific DNA-binding protein: its Myb/SANT domain mediates protein and chromatin interactions rather than base-specific contacts, whereas REST binds RE1 elements through a C2H2 zinc-finger domain.[7] ARES nonetheless identified the REST motif as the strongest non-self predictor of RCOR1 ChIP-seq signal in HepG2.

The report resolved this to tethering. Across all RCOR1 peaks the REST motif score correlated positively with RCOR1 signal (Spearman $r = 0.39$, $P < 10^{-15}$), but within RCOR1 peaks lacking REST occupancy the correlation was abolished ($r = 0.03$, $P = 0.44$), indicating that the motif predicts RCOR1 binding only through REST occupancy. REST and RCOR1 have a maximum-confidence physical interaction (STRING score 999). Only 8.8% of RCOR1 peaks are co-bound by REST, but those peaks carry markedly higher RCOR1 signal (fold change 2.34, Cohen's $d = 1.13$; fold change 1.52 after matching for GC content and peak characteristics, $P < 10^{-15}$), so this minority subpopulation carries a disproportionate share of the signal variance the random forest models. Genes nearest to co-bound sites were enriched for neuronal and synaptic pathways (synaptic signalling, $q = 1.6 \times 10^{-13}$) and were expressed at lower levels than genes near RCOR1-only sites (median TPM 5.11 versus 17.09; fold change 0.30, $P < 10^{-15}$), consistent with CoREST-mediated silencing of lineage-inappropriate neuronal programmes in a hepatocyte background.

ARES therefore assigned RCOR1 ← REST motif (HepG2) to the PROTEIN route with a tethering mechanism, and REST as the motif reader. The evidence is observational and does not establish causality; the report recommends REST knockdown followed by RCOR1 ChIP-seq as the direct causal test.

**Supplementary Note 3 | Examples of target-as-reader assignments.**
In target-as-reader assignments, ARES inferred that the partner motif was best explained as a sequence read by the target factor itself rather than as the binding site of a distinct trans-acting reader. This can occur because motif databases assign profiles to specific factors or fine-grained families, whereas DNA-binding specificity is often shared across broader DNA-binding-domain classes.[8–10] Thus, a motif can escape the self/paralog exclusion filter because its JASPAR namesake belongs to a different annotated family, while still containing a sequence grammar recognized by the target.

**Shared E-box recognition across bHLH subclasses.** The most recurrent examples involved USF1 or USF2 targets paired with ARNTL/ARNT2-labeled motifs, observed across seven cell types. USF factors are bHLH-ZIP proteins, whereas ARNTL/BMAL1 and ARNT2 are bHLH-PAS proteins; both classes recognize E-box-like sequences centered on CACGTG.[11–14] Thus, although the ARNTL/ARNT2 motif labels were not removed by the self/paralog filter, the motif sequence itself is compatible with direct USF recognition. A similar pattern was observed for MAX paired with the HES1-labeled motif in multiple cell types, where both motifs contain an E/N-box-like CACGTG core despite distinct JASPAR family labels.[15,16]

**Shared GC-box recognition among zinc-finger factors.** A second class involved SP/KLF-family targets paired with the ZNF610-labeled motif, including SP1, SP4 and KLF12 dependencies. SP and KLF factors recognize GC-rich boxes, and the ZNF610-labeled motif is itself strongly GC-rich.[17] ARES therefore interpreted these dependencies as direct GC-box recognition by the SP/KLF target rather than as evidence that the motif namesake factor was the required reader.

**Shared nuclear-receptor half-sites.** A third example involved NR2F1 paired with a THRB-labeled motif in HepG2. Both factors belong to the nuclear-receptor class and recognize AGGTCA-based half-sites or repeats.[18] The THRB-labeled motif therefore contains a sequence grammar compatible with NR2F1 binding, explaining why ARES assigned the target itself as the reader despite the different motif namesake.

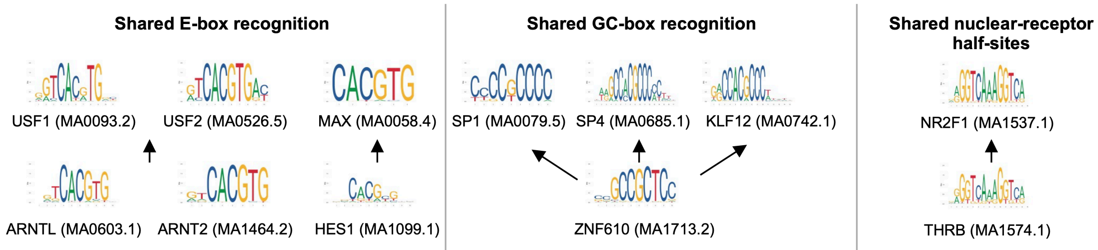


These examples illustrate why a target-as-reader assignment is not necessarily inconsistent with partner-motif selection. The partner motif is “non-self” with respect to the database label and exclusion rule, but it can still encode a sequence grammar recognized by the target factor. Thus, the target-as-reader category captures cases in which database labels and biological DNA-recognition space do not align perfectly. These examples above are representative rather than a complete manual taxonomy of all target-as-reader assignments.

**Supplementary Note 4 | RNA-seq-derived transcriptional outcomes are not determined by canonical reader's GO valence.**

To test whether ARES-transcriptional outcome annotation was independent of reader identity, we compared ARES annotation with canonical activator/repressor reader labels assigned from Gene Ontology. Of the 109 dependencies with ARES-transcriptional outcome annotation, 72 had a reader with a resolvable canonical GO valence (the other 37 had no clean activator/repressor annotation). Among these 72, ARES annotation matched the reader's canonical valence in only 21 (29%) and was discordant in 51 (71%), including 19 (26%) in which it was reversed. A function-derived label would be ~100% concordant; the observed 29% supports that ARES' outcome was measured with RNA-seq, not inferred from the reader.

| Reader canonical GO valence | ACTIVATING | NO_EFFECT | REPRESSIVE | Total |
|---|---|---|---|---|
| Activator | 20 | 27 | 5 | 52 |
| Repressor | 14 | 5 | 1 | 20 |

Representative discordant cases:

- HEY1/$M_{GATA5}$ (K562): ARES inferred reader as GATA2 (canonical activators); motif-positive genes lower-expressed (FC 0.618, $P = 2 \times 10^{-6}$); ARES annotated the transcriptional outcome as REPRESSIVE.
- EP300/$M_{CEBPD}$ (HepG2): reader CEBPA;CEBPB (GO annotated as activators); no significant expression difference; annotated NO_EFFECT.
- IRF4/$M_{SPIB}$ (GM12878): reader SPI1/PU.1 (GO annotated as activator); motif-positive genes not higher (mean TPM 35.9 vs 39.7); annotated NO_EFFECT.

**Supplementary Table S1 | Convergence of ARES analyses across atlas routes.**
Numbers of dependencies for which ARES converged or did not converge within a maximum of 15 hypothesis-evaluation iterations, stratified by final atlas route. An analysis was considered converged when the accumulated evidence satisfied all predefined evidential criteria for terminating the iterative evaluation; analyses that did not satisfy all criteria after 15 iterations were classified as not converged. Final route labels were assigned separately from convergence status. Dependencies were classified as SEQUENCE, PROTEIN or CONTEXT when the available evidence supported a corresponding mechanistic route, as UNRESOLVED when the evidence did not support any single route, and as ARTIFACT when the predictive dependency was attributed to a technical or analytical artifact. Percentages indicate the fraction of dependencies within each route that did not converge. The atlas row reports totals across all 1,552 dependencies.

| Route | n | Converged | Not converged | % not converged |
|:---:|---:|---:|---:|---:|
| UNRESOLVED | 391 | 34 | 357 | 91.3% |
| SEQUENCE | 300 | 220 | 80 | 26.7% |
| CONTEXT | 483 | 436 | 47 | 9.7% |
| PROTEIN | 363 | 330 | 33 | 9.1% |
| ARTIFACT | 15 | 15 | 0 | 0.0% |
| Atlas | 1,552 | 1,035 | 517 | 33.3% |

**Supplementary Table S2 | Route-class definitions used in the first-stage extraction.** Each ARES report was assigned to one of five route classes describing why the partner motif predicts the target TF's binding: SEQUENCE, PROTEIN, CONTEXT, UNRESOLVED or ARTIFACT. Assignment used a trace classifier (DeepSeek v4-pro) that reads the full report and follows the causal path from the partner motif to a reader (the protein that binds the motif), an actor (the protein whose binding directly moves the target's signal) and the target, deciding PROTEIN versus CONTEXT by the provenance of the actor (delivered by the reader's chain versus by the chromatin state; see the counterfactual in the criterion column). The reported class is the majority over five independent draws (temperature 0); ties were assigned UNRESOLVED, and a deterministic guard reassigned a call to UNRESOLVED when no hypothesis in the report supported a mechanism. The same call also recorded the reader, actor, actor provenance, protein sub-type, region of regulation and transcriptional outcome. Definitions and criteria are the operational instructions given to the classifier.

| Route class | Definition | Assignment criterion (trace) |
|---|---|---|
| SEQUENCE | The target's signal gain comes from a meaningful, target-specific regulatory sequence or grammar, with no protein functionally required to bring the target. | Any of: the target's own DBD recognizes the motif; the motif is a half-site nested in the target's own motif ("Trojan-horse"); or a constrained composite grammar with the target's motif facilitates binding (scrambling the grammar, not removing a protein, abolishes it). A protein may read the motif, but only if it does not act on the target. reader = target (or grammar-only reader); actor = none. |
| PROTEIN | A protein that reads the partner motif, or a cofactor it brings, directly moves the target's signal (recruits, tethers, stabilizes, co-binds or excludes it), and that protein is delivered by the reader's chain. | Provenance = reader_chain: removing the motif or its reader removes the actor's action on the target. Sub-types: tethering (target reads no DNA of its own), cooperative (target reads its own half-site but requires the partner), exclusion (the reader displaces the target; the motif negatively predicts binding). |
| CONTEXT | The partner motif marks a chromatin state or genomic region to which the target responds. Either no protein reads the motif, or a protein acts on the target but was delivered by the context rather than by a motif-reader. | Provenance = context: the actor would still act on the target if the motif were gone; the motif only marks the region (accessibility, active/repressive marks, HOT regions, compartments/loops). |
| UNRESOLVED | The report does not establish a clear mechanism to be confidently assigned to the three routes. | No reader/actor chain resolved; tied votes; or the deterministic guard when no hypothesis supported a mechanism. |
| ARTIFACT | The partner motif does not actually predict the target's binding. | The report shows the motif is not a genuine driver; most likely, the target factor is not expressed and the partner motif is not enriched. |

**Supplementary Table S3 | Mechanism-leaf taxonomy within each resolved route.**
Within the SEQUENCE, PROTEIN, CONTEXT and ARTIFACT routes, a second-stage classifier assigned each dependency a specific mechanism leaf chosen only from that route's menu. The reported leaf is the value receiving at least two of five independent votes; reports without a two-vote consensus were flagged for curator review. Definitions and defining criteria are condensed from the classifier schema. UNRESOLVED dependencies carry no mechanism leaf.

| Route | Mechanism leaf | Definition | Defining feature / criterion |
|---|---|---|---|
| SEQUENCE | Direct sequence recognition | The target's own DBD directly reads the partner motif, or a sub-sequence / nested ("Trojan-horse") motif it contains, often because target and partner families share a motif preference (KLF/SP on GC-boxes, ETS on GGAA, bHLH on E-boxes). | The target itself recognizes the motif. Also covers the case where the target reads and competes with motif namesake for binding at the sites. |
| | Motif grammar | The target reads a multi-motif DNA grammar that includes the partner motif: a fused/overlapping/embedded composite element, or two motifs at constrained spacing / rigid helical phase (ETS–GATA, EICE ETS–IRF). | The target reads the composite / grammar itself. |
| PROTEIN | Tethering | The target reads no DNA of its own at these sites; a motif-reading protein anchors it there by protein–protein interaction, either directly or through a bridging cofactor / complex / third factor. | Target’s own motif is absent at the sites, or the target cannot bind without the protein reading the motif; removing the motif evicts the reader and the target drops with it. |
| | Cooperative co-binding | The target binds its own DNA at these sites (own half-site or an adjacent / composite motif) and the partner-motif-reading protein contributes; both factors occupy DNA (heterodimer or cooperative adjacent binding). | Assigned only when the report shows the target reads its own sequence here; higher signal at co-bound sites alone is not sufficient. |
| | Co-occupancy | The partner-motif reader and the target occupy the same sites, but the report does not resolve whether the target contacts its own DNA (cooperative) or is recruited without DNA contact (tethering). | Co-occupancy or signal amplification shown, but the target's own DNA contact is unresolved. |
| | Competitive exclusion | A protein that reads the partner motif displaces or blocks the target from those sites; the target does not read the motif and is out-competed. | The motif negatively predicts (anti-correlates with) target binding. |
| CONTEXT | Chromatin state | The motif marks a histone-defined chromatin state or regulatory-element class (active/poised/bivalent promoter or enhancer; H3K4me3/me1, H3K27me3, H3K9me3) where the target binds with different intensity. | Chromatin state drives the association. |
| | Accessibility proxy | The motif marks accessible / open chromatin (DNase). | Controlling accessibility reduces association between motif and target. |
| | Sequence-feature proxy | The motif captures a DNA-intrinsic feature other than a binding site: CpG density / CpG islands, GC content, k-mer composition, low-complexity / repeat sequence, or DNA shape. | The association is explained by that intrinsic feature. |
| | Surrogate-TF co-occupancy | The partner motif is actually a binding site for a different TF (a partner-paralog or an abundant factor with similar motif preference, e.g. SP1/SP2 on a GC-rich motif) that co-occupies target peaks but does not recruit or stabilize the target. | No recruitment of the target is established; the surrogate is a passive reader of the motif, often when the named partner is not expressed. |
| | HOT-region co-occupancy | The locus is a hyper-occupied (HOT) region, or a shared master / lineage TF (AP-1/MYC/GATA/SPI1) independently drives both partner and target occupancy; co-binding is not pair-specific and the target's own binding logic is intact. | The relationship weakens after HOT / shared-driver filtering. |
| | 3D-loop context | The motif marks an architectural compartment (CTCF/Hi-C loop anchors, TAD boundaries) where the target binds; protein-level cooperation is rejected. | Associative co-localization via architecture, not a causal partner to target loop. |
| ARTIFACT | Technical artifact | Antibody cross-reactivity, peak-calling noise, mappability bias, or a feature proxy the ML model latched onto with no biological basis. | The association is technical or modelling in origin, with no genuine biological dependency. |

### Supplementary Table S4 | Faithfulness audit of the one-shot baseline (GPT-5.5) across the 50-dependency benchmark.

| Target/Partner | Cell | Category | Failure mode | What the baseline did | HOW ARES safeguard |
|---|---|---|---|---|---|
| ATF3/USF2 | GM12878 | Major | Phantom analysis | Reported a standardized linear model ($\beta$=0.398; $R^2$ 0.266→0.112), a chromatin-adjusted partial Spearman (0.388, p=$3\times10^{-144}$) and Hi-C loop-anchor %s — no regression, partial-correlation or Hi-C code exists | Forced execution before conclusion: a solution cannot form without a corresponding execution log, so an analysis that was never run produces no output to report. The entity-fidelity check flags any factor named but absent from the log |
| ATF2/ATF3 | HEK293 | Major | | TSS-distance %s (28.3% vs 17.9%; 9.4 kb) — GENCODE never loaded, no distance computed | |
| CTCF/PHOX2B | H1 | Major | | Multivariable regression coefficient (PHOX2B 0.109) — no model code saved | |
| HOXA10/MEIS2 | HepG2 | Major | | Hi-C loop-anchor overlap 30.2% — loops file defined but never used | |
| ZNF169/EGR2 | HEK293 | Major | | GC medians (0.60 vs 0.51) + motif↔GC r=0.488, called "directly computed" — no GC code exists | |
| SP1/HNF4A | HepG2 | Major | Fabricated number | Cohesin SMC3 25.9% vs 11.3% (2.30-fold) — no SMC3 row in the saved table; never computed (2.30 ≈ EP300's 2.307) | Entity-fidelity check (deterministic): SMC3 is absent from the execution log and is flagged. Number-verification step re-derives every reported statistic from a variable |
| ZNF571/REST | HepG2 | Major | | Five numbers absent from any saved output | |
| ZEB2/GATA5 | K562 | Major | Direction reversed | Claimed motif-positive windows have higher signal + dominate top decile; saved output shows the opposite; top-decile columns relabeled "motif-positive" | Number-verification step (directionality and denominator checks); interpretation-ceiling / control-override at evaluation |
| ZNF317/KLF12 | K562 | Major | Mislabeled entity | An AHR-derived result reported under the KLF12 label | Entity-from-variable rule (required, prompt-enforced): every named entity must originate from a variable computed in the same block. Not caught by the deterministic entity check, since the KLF12 label is itself present in the log |
| RORB/FOXE1 | WTC11 | Minor | Imprecise wording | Aside says "only one high-confidence STRING edge"; saved output lists three. Central conclusion unaffected | Summary-agent wording and magnitude discipline (required; low impact) |
| AEBP2/ONECUT1 | HEK293 | Unsupported | Number lacks saved computation | Log-linear $R^2$ deltas (0.343→0.431; 0.029→0.114) with no traceable computation; core claims verified | Number-verification step (required): every reported number re-derived from a data variable before a solution |
| FOS/ISL1 | GM12878 | Unsupported | | Secondary cofactor signal medians + ISL1/GC Spearman with no saved code for those numbers; conclusion-driving claims verified | Number-verification step (required) |
| TRIM22/CTCF | MCF-7 | Unverifiable | No saved code or output | Saved neither code nor outputs; none of the numbers (74.4% overlap, 60.3-fold, 18 bp) checkable | Persistence foundation (deterministic): every executed block and its output are persisted, so a report with no saved computation cannot be produced |
| BRF2/TBP | HeLa.S3 | Unverifiable | | Saved nothing; entire report (16/32 overlap; Spearman $\rho$=0.804, p=$3.1\times10^{-8}$) uncheckable | |

**Supplementary Table S5 | Classification of the 22 covered bQTL dependencies by the mechanism type inferred in their ARES reports.**

Each mechanistic conclusion is quoted from the dependency's ARES report. Dependencies were assigned to two groups by whether the report nominates a directional protein chain, not by any ratio. For protein-chain candidates (n = 8), the report nominates a specific directional protein complex (a reader, bridge or heterodimeric partner), and an AlphaGenome concordance ratio testing whether the allele effect propagates through the named chain is defined and reported (Fig. 5e). For the remaining 14 dependencies, the report infers co-occupancy without a direct protein interaction, so no directional chain is available to test and a concordance ratio is not defined. Assignment is by mechanism type, not by ratio magnitude: a proxy path constructed for several of the non-chain dependencies would exceed the concordance threshold (for example LEF1/GATA5 would give 2.33), yet these dependencies remain excluded from the concordance analysis because their reports do not specify a directional protein chain.

| Target/Partner | Cell | Mechanistic conclusion from ARES report |
|---|---|---|
| **Protein-chain candidates (n=8)** | | |
| CEBPG/ATF4 | K562 | CEBPG heterodimerizes with ATF4 and is physically recruited or stabilized at shared stress-response loci, so the ATF4 motif marks the sites the two proteins co-occupy. |
| FOSL1/JUN | K562 | FOSL1 and JUN directly interact to form the AP-1 heterodimer, so JUN-motif loci recruit and stabilize FOSL1, giving higher FOSL1 signal at co-occupied sites. |
| IKZF1/RUNX1 | GM12878 | RUNX motifs predict IKZF1 indirectly: RUNX3 and IRF4 co-occupy active enhancers, forming a ternary complex that recruits NuRD/CHD4 and creates a favorable environment for IKZF1 binding. |
| ZNF143/THAP11 | HepG2 | ZNF143 and THAP11 form a direct protein complex that extensively co-occupies the genome, so the THAP11 motif marks co-bound loci where ZNF143 binding is cooperatively enhanced. |
| GATA3/ FOXP3 | MCF-7 | The FOXP3 motif is a proxy for FOXA1 sites; GATA3 lacks its own motif and is tethered to FOXA1 through an ESR1-bridged co-occupancy complex. |
| CEBPB/ATF4 | K562 | At sites lacking a CEBPB motif, ATF4 binds its own motif in accessible chromatin and recruits or stabilizes CEBPB, so CEBPB occupancy tracks ATF4 protein rather than sequence. |
| NFE2/MAFK | K562 | NFE2 and MAFK physically heterodimerize and co-occupy shared regulatory elements, so the MAFK-motif sites mark their co-bound loci. |
| ATF2/ATF3 | HepG2 | ATF2 and ATF3 form a heterodimeric complex that co-occupies a subset of stress-response loci, where the ATF3 motif facilitates recruitment or stabilization of ATF2-ATF3 complex. |
| **No directional chain (n=14)** | | |
| ETV1/GATA5 | K562 | GATA factors facilitate ETV1 binding through pioneer-like opening of local accessibility plus sub-threshold GATA co-occupancy that tethers and stabilizes ETV1, not direct interaction or sequence affinity. |
| MEIS2/NRF1 | K562 | Not a direct interaction: NRF1 co-occupies a distinct subset of actively transcribed regulatory elements (Pol II, elevated metabolic-gene expression) that the NRF1 motif marks. |
| MIXL1/FOXL1 | HepG2 | MIXL1 and FOXA2 co-occupy accessible enhancers where their co-binding recruits EP300 and elevates H3K27ac, rather than a direct motif-driven cooperative complex. |
| TEAD4/GATA5 | K562 | GATA1/2 read the motif and co-occupy shared regulatory elements with TEAD4, where co-occupancy is associated with elevated EP300 independent of baseline H3K27ac; direct cooperative binding was inconclusive. |
| LEF1/GATA5 | K562 | LEF1 is indirectly tethered to active enhancers by the erythroid GATA-TAL1 complex rather than binding its own motif; this tethering dominates at LEF1-motif-negative, high-H3K27ac sites. |
| TFE3/ARNTL | HepG2 | The ARNTL motif marks ARNTL co-occupancy with TFE3, which recruits EP300 and deposits H3K27ac to epigenetically prime these shared enhancers (observational, non-causal). |
| SP2/ZNF610 | HEK293 | The ZNF610 motif is a sequence proxy for KLF1 scaffold binding at GC-rich active elements (H3K27ac / H3K4me3), where KLF1 co-occupies with SP2 rather than interacting directly. |
| ZNF660/ATF3 | HEK293 | The ATF3 motif is a proxy for ATF2 occupancy (the two share bZIP CRE/TRE-like motifs); ZNF660 co-occupies these ATF2-bound regulatory regions. |
| NFIC/GATA5 | K562 | GATA motifs mark dense, highly accessible erythroid multi-TF hubs where NFIC binds opportunistically at lower affinity, not through direct cooperation, competition, or tethering. |
| EBF1/RELB | GM12878 | The RELB motif predicts EBF1 because its paralog RELA, sharing the NF-kB motif, occupies these sites and co-occupies conserved stress-response enhancers alongside EBF1. |
| MAX/TCFL5 | A549 | The TCFL5 motif is a variant E-box that MAX preferentially co-occupies as part of shared MYC and E2F6 complexes at transcriptionally active metabolic-gene loci, rather than binding it directly. |
| NRF1/TCFL5 | K562 | The TCFL5 motif mimics the E-box and acts as a sequence proxy for MAX binding, whose co-occupancy at these sites is associated with elevated NRF1 signal (independent of GC and accessibility). |

| | | |
|---|---|---|
| E2F6/TCFL5 | A549 | The TCFL5 motif is a canonical E-box directly bound by USF1/USF2, whose co-occupancy raises local chromatin accessibility that correlates with stronger E2F6 binding, not a direct E2F6-TCFL5 interaction. |
| NRF1/ZBTB14 | HepG2 | The ZBTB14 motif is a proxy binding site for the cofactor GABPB1, whose co-occupancy at these motifs interacts with NRF1 and is associated with elevated NRF1 ChIP signal. |
